# Systemic and kinematic properties of *Amoeba proteus* migration correlate strongly across environments

**DOI:** 10.64898/2026.10.20.759940

**Authors:** José Carrasco-Pujante, Iker Malaina, Gorka Pérez-Yarza, Borja Camino-Pontes, Jesús M. Cortés, José I. López, Ildefonso M. De la Fuente

## Abstract

Recent work has shown that cell migration is a systemic property of the whole cell, yet how the different properties of a migratory trajectory relate to one another remains largely uncharacterized. Here we quantified the migration of 150 Amoeba proteus cells under three experimental scenarios (absence of stimuli, a direct-current electric field, and a chemotactic peptide gradient) with eleven metrics that describe either the systemic dynamics of the trajectory (long-range correlations, long-term persistence, super-diffusive exploration, regularity of the move steps and memory of the velocity) or its kinematic output (net displacement, straightness and speed). Screening all 55 metric pairs for statistical dependence with distance covariance identified six couplings that held in all three experimental scenarios, besides the pairs linked by definition. Long-term persistence and super-diffusive exploration were coupled with each other, in agreement with the scaling relation expected for self-similar trajectories, and each of them was coupled with the net displacement and the straightness of the trajectory: more persistent and more super-diffusive cells moved farther and along straighter paths. In addition, the long-range correlations of the move steps were coupled with the memory of speed between consecutive steps. All six couplings were positive and monotonic, with no trend reversal over the observed range. The stimuli shifted six of the eleven metrics, the electric field mainly increasing the straightness and the net displacement of the trajectories and the chemical gradient mainly increasing the speed memory, but they left the six couplings intact and reorganized part of the dependence structure around speed. These results show that several systemic and kinematic properties of *Amoeba proteus* migration are not independent observables but vary together as parts of an integrated system whose couplings are preserved across environments.

## Introduction

Cellular migration is an essential property of myriad prokaryotic and eukaryotic organisms. It plays a critical role in the survival of free-living unicellular organisms while navigating complex microenvironments, foraging and avoiding predators. In multicellular organisms, it is also fundamental for developmental and physiological processes including organogenesis, embryogenesis, adult tissue remodeling, angiogenesis, cell differentiation, morphogenesis, immune responses, and wound healing [1–3].

Dysregulated cell motility is closely linked to the onset and development of human disease, including metastatic cancer progression [4–8], congenital brain disorders [9–11], atherosclerosis [12], congenital cardiomyopathies [13,14], skeletal and cranial anomalies [14], chronic obstructive pulmonary disease [15], osteoarthritis [16], endometriosis [17,18], multiple sclerosis [19,20], diabetic wound [21], asthma [22–24], progeria [25], rheumatoid arthritis [26,27], Crohn’s disease [28,29], auditory impairments [30], fibrosis [31–33], psoriasis [34,35], immune-related actinopathies [36], and other immunodeficiencies [37].

Directional cell migration has been the subject of extensive research aimed at elucidating the molecular components that drive motility. Experimental studies revealed that locomotion is a highly complex process involving virtually all cellular components. Essentially, directed movement is driven by the cytoskeleton, a dynamic architecture composed of actin microfilaments, microtubules, and intermediate filaments that interact through sophisticated regulatory networks [38]. Integrative functional coordination through these networks underlies the emergence of systemic behaviors that drive cell functionality [39–44].

Recent research has shown that cell migration does not depend on any specific individual cellular component or molecular-metabolic process, but rather emerges as a systemic property of the interaction of a vast number of cellular molecular components, organelles and metabolic processes [45]. In that study, researchers used computational and physico-mathematical tools to quantitatively analyze the trajectories of 700 individual cells from three species of free-living amoebae (*Amoeba proteus*, *Metamoeba leningradensis* and *Amoeba borokensis*) under four different experimental conditions: absence of stimuli, a direct-current electric field (galvanotaxis), a chemotactic gradient (chemotaxis), and simultaneous galvanotactic and chemotactic stimuli. All cells exhibited a complex migratory structure regardless of species and experimental conditions. This structure, characterized by robust long-range correlations, super-diffusive behavior, strong long-term persistence, and highly regular move-step displacement sequences, reflecting highly coordinated and information-rich locomotion, disappeared when the migration trajectory time series were randomized. This finding revealed that cell locomotion is characterized by coherent integrative responses operating at a global cellular scale, driven by cooperative, nonlinear interactions of myriad cellular molecular components.

Additionally, these integrative locomotion responses were verified in enucleated cells through a quantitative study comparing the migratory dynamics of 200 enucleated (cytoplasts) *Amoeba proteus* and 200 intact cells. Both cytoplasts and nucleated cells displayed similar complex migratory behaviors, characterized by long-range correlations, anomalous super-diffusive migration, long-term trend-reinforcing behavior (persistence), and highly ordered move-step sequences under the four different experimental conditions tested (absence of stimuli, a direct-current electric field, a chemotactic gradient, and simultaneous galvanotactic and chemotactic stimuli). These findings suggest that self-organized cytoplasmic processes predominantly govern cell migration [46].

*Amoeba proteus* is a free-living amoeba species that has been used as a model organism in cell migration studies for over a century [47–50]. While *Amoeba proteus*’ reactions to galvanotaxis [51], chemotaxis [52], and mechanical shock [53] are well studied, more recent work has quantitatively established that cell migration under different stimuli is characterized by complex integrative dynamics [45], and that these dynamics are preserved in the motility of enucleated cells (cytoplasts) [46]. Further studies have demonstrated its ability to anticipate periodic stimuli [54] and to display motility patterns consistent with associative conditioning [55]. Quantitative analyses of Amoeba proteus locomotion have a substantial precedent in trajectory, stochastic, and nonlinear-dynamics studies [56–58], complemented by biophysical measurements of intracellular pressure, calcium-dependent contractility, and cytoplasmic rheology [59–61]. After this earlier work, systematic statistical characterization of whole-cell migratory trajectories was revisited in 2019 and expanded in multi-condition and enucleation studies [45,46,62]. These research efforts establish this organism as a model for the study of the integration of internal mechanical state and environmental cues in single-cell motility.

Building upon these findings, the present study aims to advance the understanding of the dynamic aspects underlying cellular migration, which, despite the abundance of studies on the properties of migrating cells, remain insufficiently understood. Cell migration has been extensively modeled using persistent random-walk theory [63–67]. Maiuri et al. (2015) [68] reported exponential coupling between cell speed and persistence in multiple cell types, with mechanistic models linking actin dynamics to this coupling [69], directional persistence itself emerging from protrusion dynamics and polarized intracellular trafficking [70], and the coupling enhancing search efficiency in immune cells [71]. Several computational frameworks have been developed to systematically quantify and compare migration dynamics [72–74]. Entropy-based measures have recently been applied to cell migration: the Shannon entropy of velocity power spectra and of angular-displacement distributions correlates with persistence in mammalian cells [75,76], and has been used to quantify migration persistence across multiple cell types including amoebae. Morphological entropy characterizes strategy switching across length scales [77], and the Shannon entropy of the distribution of step directions has been correlated with migration persistence and cell shape in ovarian cancer cells [78]. These applications all rest on Shannon-type entropies, which measure the spread of a distribution and are therefore insensitive to the order in which values occur. Approximate Entropy instead measures the regularity of the sequence itself, quantifying how reliably a short pattern of move steps is followed by a similar one. That distinction is what makes ApEn the appropriate entropy here, because the systemic organization of migration is expressed in the ordering of the move steps rather than in their marginal distribution. Single confined cells can exhibit nonlinear dynamics not captured by linear models [79], a perspective extensively reviewed in the recent literature on data-driven modeling of cell migration [80].

To our knowledge, no previous study has quantified the dependence between the systemic (RMSF α, DFA γ, MSD β) and the kinematic (IR, DR, AS) metrics (defined below) of the same trajectories across environments; the closest published works report exponential coupling between purely kinematic quantities [68]. A scaling exponent has also been used on its own: the persistence of random amoeboid motion in *Dictyostelium* was quantified with the Hurst exponent of fractional Brownian motion [81], to which MSD β and DFA γ are tied for a self-similar trajectory. Approximate Entropy has been applied to amoeboid migration trajectories [45,46], but its dependence on persistence has not been tested.

Here, we study the relationship between systemic metrics commonly used in cell migration studies and the kinematic descriptors of the resulting trajectory. The systemic set comprises the Root Mean Square Fluctuation (RMSF) exponent α and the RMSF correlation time, which measure respectively the scaling of the fluctuations in the displacement signal and the time over which those fluctuations stay correlated; the Mean Squared Displacement (MSD) exponent β, which separates diffusive from super-diffusive exploration; the Detrended Fluctuation Analysis (DFA) exponent γ, which quantifies long-term persistence; and Approximate Entropy (ApEn), which measures the regularity of the move-step sequence. The kinematic set comprises the Intensity of the Response (IR), the Directionality Ratio (DR) and the Average Speed (AS), which capture the magnitude, the straightness and the pace of the displacement respectively. The systemic set also includes the memory retained in the velocity itself, separated into a speed memory, a direction memory and a coupling between the two [82]. The two sets are distinguished by a single criterion: the kinematic metrics depend only on the step increments of the trajectory and not on their order, whereas every systemic metric depends on that order (Materials and Methods).

Before presenting the results, we outline the analysis as a whole, so that each section can be read against the overall design. We recorded the migration trajectories of 150 Amoeba proteus cells, 50 in each of three experimental scenarios: the absence of stimuli, a direct-current electric field, and a chemotactic peptide gradient. For every trajectory we computed eleven metrics, comprising systemic descriptors of the migratory dynamics (the two RMSF descriptors, the Detrended Fluctuation Analysis and Mean Squared Displacement scaling exponents, Approximate Entropy and three descriptors of velocity memory) and kinematic descriptors of trajectory geometry (Intensity of the Response, Directionality Ratio and Average Speed). We then screened all 55 pairs among those eleven metrics for statistical dependence with a distance covariance permutation test, corrected for multiple comparisons, and retained as coupled the pairs that were dependent in all three scenarios and not linked by definition. Six pairs met this criterion. We quantified their strength with the bias-corrected squared distance correlation and their direction with the Spearman correlation, and examined their shape with tests for nonlinearity and trend reversal and with a comparison of candidate functional forms. Finally, we tested every metric for stimulus effects and compared the couplings across the three scenarios.

## Results

The migration trajectories of 150 *Amoeba proteus* cells were recorded under three different experimental scenarios (50 cells per scenario): free movement in the absence of external stimuli, galvanotactic behavior under an electric field, and chemotactic behavior in the presence of a peptide gradient (nFMLP, N-Formylmethionyl-leucyl-phenylalanine, also known as chemotactic peptide). Fig 1 shows 25 representative cell migration trajectories for each experimental scenario and a prototypical *Amoeba proteus* cell migration trajectory in detail.

**Fig 1.**
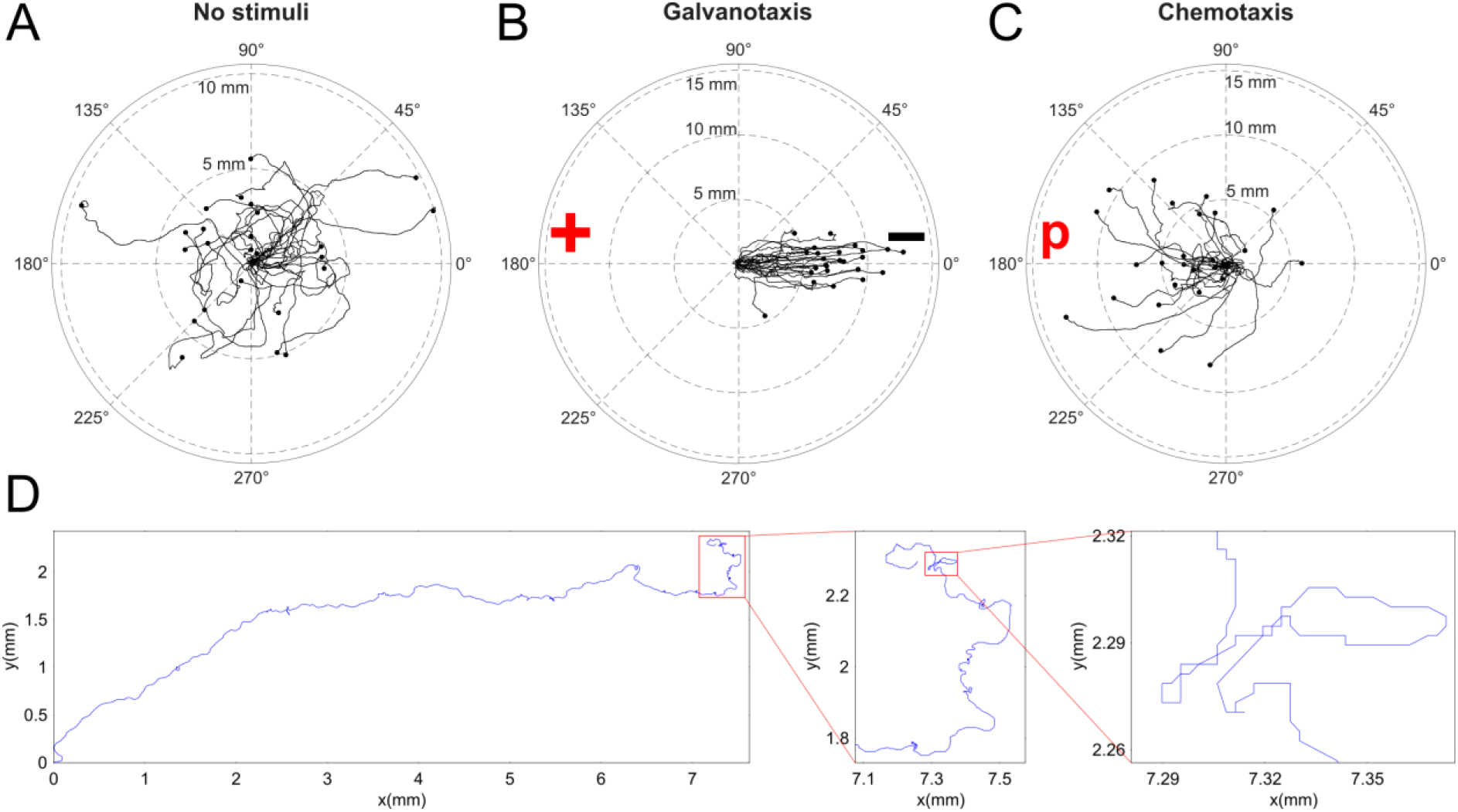
Representative migratory trajectories under the three experimental scenarios considered. **(A)** Migratory trajectories of 25 representative cells in the absence of stimuli, **(B)** under a direct-current electric field (−, cathode; +, anode) and **(C)** in the presence of a chemotactic peptide gradient (p). **(D)** Digitized cell trajectory with two insets displaying regions of interest in the cellular displacement. In (A–C), the angle and the radius give the direction and the distance (mm) from each cell’s starting point. In (D), the x and y axes give the position in mm. The electric field straightens the trajectories and orients them toward the cathode, and the peptide gradient orients most of them toward its source.

All the galvanotactic experiments were performed in a custom experimental set-up consisting of two standard 17.5 cm electrophoresis blocks, two agar bridges, a power supply, and an experimental chamber made from a standard glass slide and coverslips to house the cells (Fig 2 and Materials and Methods). One electrophoresis block was connected to the power supply, while the second block was connected to the first using the agar bridges. This arrangement effectively isolated the electrodes from the cell medium. The amoebae were placed in the center of the experimental chamber to monitor their migratory displacements. The glass chamber facilitated laminar flow, allowed for the passage of the electric current, and enabled the generation of an nFMLP peptide gradient (Materials and Methods). For migration in the absence of stimuli and chemotaxis experiments, where there was no need to isolate the electrodes from the cell medium, a single electrophoresis block was utilized.

**Fig 2.**
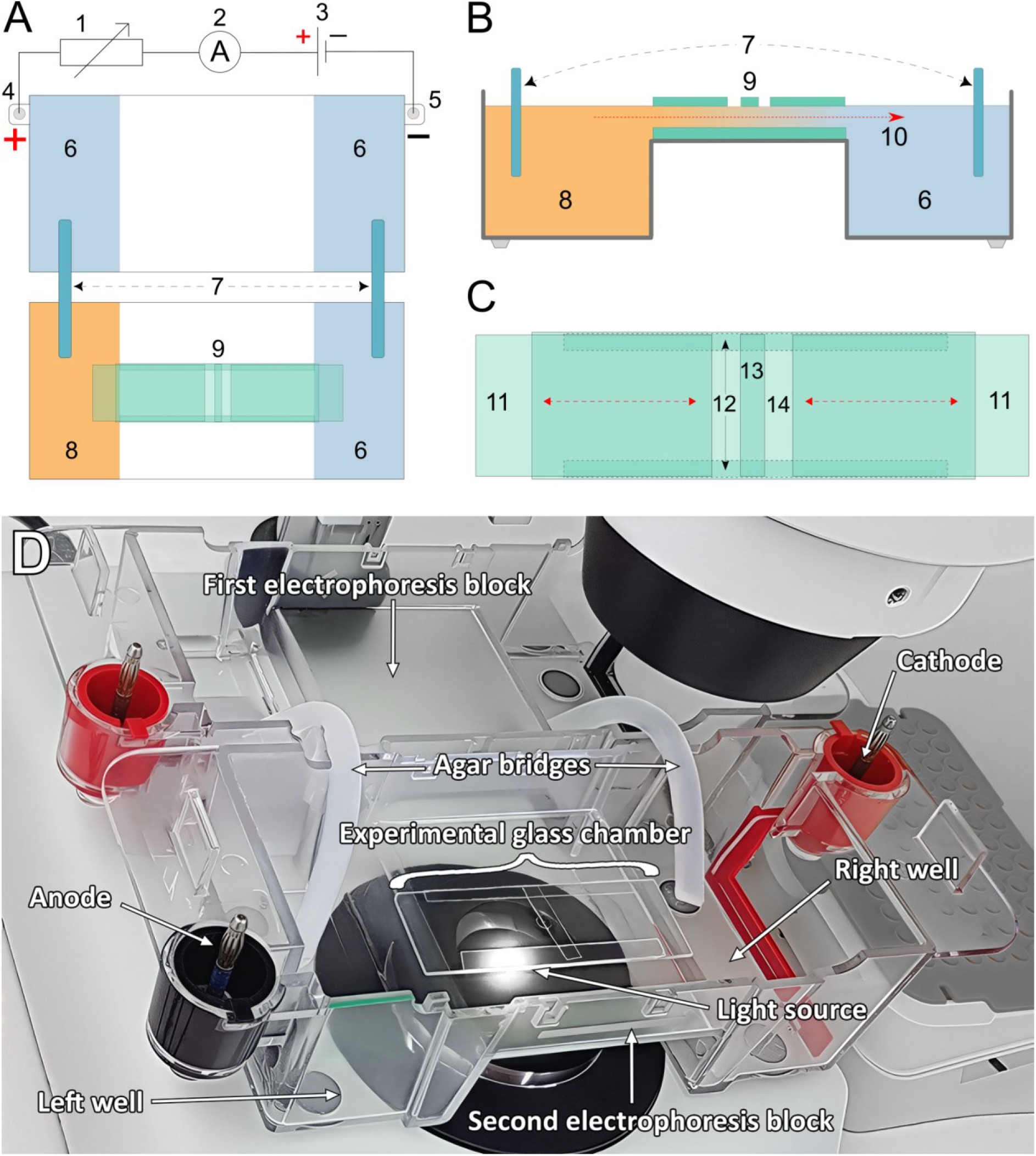
**Experimental set-up**. **(A)** and **(B)** Top and lateral views of the experimental set-up. 1: 1MΩ variable resistor; 2: ammeter; 3: DC voltage source; 4: anode; 5: cathode; 6: well containing fresh Chalkley’s simplified medium; 7: agar bridges; 8: well where the peptide was added; 9: experimental glass chamber; 10: direction and trajectory of the laminar flow (red arrow). **(C)** Top view of the experimental glass chamber. 11: sliding lateral pieces; 12: longitudinal glass strips; 13: central glass piece where the cells were placed beneath; 14: standard microscope slide. **(D)** All the elements of the set-up described in (A-C) are shown under experimental conditions.

Cells were starved for 24 h before each experiment, and their migratory trajectories were recorded using a digital camera attached to a stereomicroscope. A maximum of 8 cells were used per experiment. Finally, all cellular trajectories were analyzed as time series using computational tools (Materials and Methods). The values of the eleven metrics in every cell are given in Tables S1 to S11 in S1 File.

### Cell migration in the absence of stimuli

The trajectories of 50 cells (7 experimental replicates, 7–8 cells per replicate) were recorded in the absence of any external stimuli. Under these conditions, cell migration is described as correlated random motion, exhibiting a progressive decline over time of the original direction of migration [64]. Fig 1A shows a representative example of 25 migration trajectories of *Amoeba proteus* recorded under these conditions; cell motility exhibited significant directional changes, allowing the exploration of virtually all orientations within the experimental chamber within 30 minutes.

### Cell migration under an electric field

Migration trajectories of 50 cells (7 experimental replicates, 7–8 cells per replicate) were recorded under a controlled external direct-current electric field of 334–343 mV/mm for 30 minutes (Materials and Methods). Amoebae exhibit robust galvanotaxis in response to electric fields between 300 and 600 mV/mm [51]. Under these conditions, the vast majority of amoebae migrate toward the cathode [51]. In our experiment, as expected, all cells showed directional migration toward the cathode. A representative example of this migratory behavior is shown in Fig 1B.

### Cell migration under a peptide gradient

The migratory behavior of 50 amoebae was recorded in the presence of a chemoattractant nFMLP peptide gradient (8 experimental replicates, 5–7 cells per replicate). Amoebae displayed chemotactic responses. Specifically, the bacterial peptide nFMLP indicates the possible presence of nutrient sources nearby, provoking a significant chemotactic response [52]. In our experiment, the majority of cells (80%) showed directional migration toward the peptide source for 30 minutes. Fig 1C shows a representative example of these locomotion movements.

### Systemic migratory-dynamics metrics

Cell migration is characterized by the high information content found within the move-step sequences in migratory trajectories, indicating the high complexity that emerges from the cellular system [45]. This pattern complexity can be estimated by calculating the Approximate Entropy (ApEn), a robust approximation of the Kolmogorov–Sinai (K–S) entropy [83,84] developed by Pincus et al. in 1991 [85]. ApEn is a statistical measure used to assess the regularity of fluctuations within time-series data. Higher ApEn values indicate a time series with greater irregularity and unpredictability, whereas lower ApEn values are characteristic of time series that exhibit more repetitive and predictable patterns, less likely to happen by chance (Materials and Methods). ApEn was computed on the move-step series of each trajectory subsampled every 5.5 s (Materials and Methods). The median/IQR values were 1.175/0.076 in the absence of stimuli, 1.159/0.054 under an electric field, and 1.158/0.065 within a chemotactic peptide gradient (Table S1 in S1 File), and across the 150 trajectories ApEn spanned 1.033 to 1.302. ApEn did not differ between the scenarios (Kruskal–Wallis effect size ε^2^ = 0.029, q = 0.141, where q is the p-value adjusted for multiple comparisons by the Benjamini–Hochberg procedure; Table S12 in S1 File), and it was not dependent on any other metric in all three scenarios (Table S13 in S1 File).

Another property that characterizes migratory cell displacement is anomalous diffusion dynamics, a phenomenon indicative of efficient exploration of the cellular environment [86,87]. To assess the amount of space explored by the amoebae during their migratory displacements, we calculated the Mean Squared Displacement (MSD). This methodology, rooted in Statistical Mechanics and pioneered by Einstein [88] and von Smoluchowski [89], allowed us to quantify the spatial displacement of cells during their locomotion. The anomalous diffusion exponent (β) is the key parameter in this approach, used to distinguish between normal (Brownian) diffusion (β≅1) and anomalous diffusion (β≇1). Furthermore, values of 0 < β < 1 are indicative of sub-diffusion dynamics, while 1 < β < 2 means super-diffusion and β = 2 indicates ballistic diffusion (Materials and Methods). In this analysis, all values of coefficient β fell within the super-diffusion range, with median/IQR values of 1.863/0.099 for cells migrating in the absence of external stimuli, 1.891/0.067 for cells migrating under an electric field, and 1.874/0.100 for cells migrating within a chemotactic peptide gradient (Table S2 in S1 File). Values reached close to the ballistic limit (β = 2), consistent with persistent, directed migration. Across the 150 trajectories β spanned 1.408 to 1.986, so every cell was super-diffusive and the largest values approached the ballistic limit without reaching it; no trajectory was Brownian or sub-diffusive.

Cellular migratory displacement is also characterized by long-term persistence. To assess the persistence in cellular migratory trajectories, we used Detrended Fluctuation Analysis (DFA), a tool introduced by Peng and colleagues in 1994 [90] that has been widely applied to physiological time series [91]. In this method, the scaling exponent γ is used to detect and measure long-term persistence effects. Specifically, a fractional Brownian process shows anti-persistence when 1 < γ < 1.5, no persistence when γ ≈ 1.5, and persistence when 1.5 < γ < 2 [92] (Materials and Methods). The scaling exponent γ values obtained reflect long-term persistence across all studied trajectories, with median/IQR values of 1.953/0.078 for cells migrating without external stimuli, 1.974/0.046 for cells migrating under an electric field, and 1.952/0.045 for cells within a peptide gradient (Table S3 in S1 File). These findings show that all the experimental series exhibit positive, trend-reinforcing persistence (γ > 1.5), with most values close to the upper end of the persistent range (γ = 2), as expected for the radial distance of cells that keep moving away from their starting point.

Systemic behavior in complex systems is also characterized by the presence of dynamics with strong long-range correlations [93]. “Long-range” refers here to decay in the autocorrelation function slower than exponential decay, where a single scale dominates the decay. A classic tool to detect and study these correlations in time series is the root mean square fluctuation (RMSF) analysis, a method rooted in Statistical Mechanics and based on ideas initially proposed by Gibbs [94] and Einstein [95]. Long-range interdependence can be detected by a power-law relationship such that *F*(*l*)∼*l^α^*, where l is the number of steps in the time series. For uncorrelated data, the fluctuation exponent *α* is about 0.5, whereas *α* > 0.5 or *α* < 0.5 indicate respectively the presence of positive or negative long-range correlations (Materials and Methods). The values obtained for the fluctuation exponent *α* ranged from 0.499 to 0.806 and exceeded 0.5 in all but one trajectory, suggesting the presence of long-range correlations in the move-step displacement sequences of practically all cell migration trajectories studied. Median/IQR values were 0.654/0.110 in the absence of stimuli, 0.664/0.086 in the electric field, and 0.699/0.065 under chemotaxis (Table S4 in S1 File). Additionally, we calculated the time duration of the correlations regime and found that all trajectories present long-range correlation regimes spanning from 1.042 to 16.667 minutes, with median/IQR values of 9.896/9.375 minutes in the absence of stimuli, 10.417/8.333 minutes in the electric field and 10.417/5.208 minutes under chemotaxis. This indicates a strong influence of previous trajectory values on each cellular move step (Table S5 in S1 File).

The memory retained in the velocity itself provides a further characterization of the migratory dynamics. It can be separated into speed-memory and direction-memory components together with a coupling between speed and direction [82] (Materials and Methods). The three components were computed on trajectories subsampled every 5.5 s (Materials and Methods). The speed memory, given by the lag-one autocorrelation of the instantaneous speed AC_v_(1), was positive in every one of the 150 trajectories, with median/IQR values of 0.500/0.220 in the absence of stimuli, 0.577/0.184 under an electric field and 0.620/0.123 within the peptide gradient (Table S6 in S1 File). Across the 150 trajectories AC_v_(1) spanned 0.169 to 0.844, above the ±0.108 bound expected at the 95% level for temporally independent speeds over the 327 steps of each subsampled trajectory, so the speed of a migrating amoeba is never memoryless. The direction memory, expressed as the persistence time t_θ_ derived from the mean cosine of the turning angle, gave median/IQR values of 94.0/71.3 s, 105.0/66.2 s and 103.9/57.3 s respectively (range 33.4 to 384.8 s; Table S7 in S1 File), and the speed-direction coupling C_θ-v_, a dimensionless cross-correlation bounded in [−1, 1], gave 0.010/0.201, 0.067/0.214 and 0.000/0.290 (range −0.361 to 0.476) (Table S8 in S1 File). Unlike the speed memory, neither the direction memory nor the speed-direction coupling differed between the scenarios (Kruskal–Wallis q = 0.999 and 0.141 for |C_θ-v_|, respectively; Table S12 in S1 File).

### Kinematic metrics

In addition to the systemic analyses, we calculated three key metrics to measure the cytokinematic properties of the 150 cellular locomotion trajectories studied: Intensity of the Response (IR), Directionality Ratio (DR) and Average Speed (AS). The IR is defined as the modulus of the cell trajectory (net distance traveled from the starting point to the ending point of the trajectory) in mm and reflects the strength of the migratory response. IR values had a median/IQR of 4.998/4.808 mm in the absence of stimuli, 6.852/3.448 mm under an electric field, and 5.329/3.368 mm in the presence of a peptide gradient (Table S9 in S1 File). The DR quantifies straightness of the trajectory, with values ranging from 0, for a completely curved trajectory, to 1, for a completely straight trajectory. It was calculated as the ratio between the modulus of the trajectory and the total distance traveled by the cell (cumulative sum of move steps). DR values had a median/IQR of 0.426/0.265 in the absence of stimuli, 0.705/0.207 under an electric field, and 0.519/0.157 within a peptide gradient (Table S10 in S1 File). The AS is the total distance traveled by the cell along its trajectory divided by the duration of the recording, expressed in micrometers per second, and measures how fast the cell moves irrespective of the direction taken. Average speed values were 6.797/1.995 µm/s in the absence of stimuli, 6.099/2.392 µm/s in the electric field and 5.779/1.888 µm/s under the chemotactic peptide gradient (Table S11 in S1 File). Across the 150 trajectories the three descriptors spanned 0.608 to 13.411 mm for the intensity of the response, 0.052 to 0.875 for the directionality ratio, and 3.111 to 11.001 µm/s for the average speed. The directionality ratio is informative in that it reached neither bound: no cell followed a perfectly straight path, and none moved with the near-zero directionality of a memoryless random walk, so every trajectory carried some directional persistence while none was fully committed to a single heading. These three descriptors are not mutually independent by construction. Writing L for the total path length traveled along the trajectory and T = 1800 s for the fixed duration of the recording, the definitions above give DR = IR/L and AS = L/T, so that the three metrics are bound by the exact identity IR = DR · AS · T, in which IR is expressed in mm, DR is dimensionless, AS in µm/s and T in seconds, which makes the numerical constant 1.8 in these units. The identity is three-way, so no two of the metrics are perfectly correlated and DR and AS remain mutually unconstrained, but IR is fully determined by the other two. The IR-DR and IR-AS pairs are therefore arithmetic rather than biological, and were treated as definitional and excluded from the coupling analysis throughout. No such identity links IR or DR to any of the systemic metrics.

### Dependence among the eleven metrics

To identify the metrics that vary together, every one of the 55 metric pairs was tested for statistical dependence within each scenario with a distance covariance permutation test, which detects linear, nonlinear and non-monotonic dependence alike [96,97], and a pair was considered dependent when the test remained significant after Benjamini–Hochberg correction (q < 0.05) in all three scenarios (Materials and Methods; Table S13 in S1 File). Eight pairs met this criterion. Two of them, IR-DR and IR-AS, follow from the identity that links the three kinematic descriptors (see Kinematic Metrics section above) and were treated as definitional. The remaining six pairs are the couplings reported in the following sections: the DFA exponent γ with the MSD exponent β, with IR and with DR; the MSD exponent β with IR and with DR; and the RMSF exponent α with the speed memory AC_v_(1). The strength of each coupling was measured with the bias-corrected squared distance correlation *R*\* [97], which is close to 0 for independent metrics and increases toward 1 as the dependence strengthens (for two normally distributed metrics the squared distance correlation is an increasing function of their Pearson correlation [96], so that *R*\*_n_ values of 0.2, 0.4 and 0.6 correspond to Pearson correlations of about 0.49, 0.68 and 0.82), and is reported with its bias-corrected and accelerated (BCa) bootstrap 95% confidence interval, together with the Spearman rank correlation ρ (Table 1). All six couplings were positive and monotonic in every scenario (ρ between 0.52 and 0.87), and none reversed its trend within the observed range, either in the Lind–Mehlum test [98] or in the two-lines test [99], which assumes no functional form (Materials and Methods). Five further pairs were dependent in two of the three scenarios only (Table S13 in S1 File) and are compared between scenarios below.

**Table 1.** Dependence between the metrics of the six coupled pairs under the three experimental scenarios. *R*\*n: bias-corrected squared distance correlation, with its bias-corrected and accelerated (BCa) bootstrap 95% confidence interval; ρ: Spearman rank correlation. Every pair was dependent in all three scenarios (distance covariance permutation test, Benjamini–Hochberg q < 0.05; Table S13 in S1 File).

| Coupled pair | No stimuli |  | Galvanotaxis |  | Chemotaxis |  |
| --- | --- | --- | --- | --- | --- | --- |
| | $R_n^*$ [95% CI] | $\rho$ | $R_n^*$ [95% CI] | $\rho$ | $R_n^*$ [95% CI] | $\rho$ |
| DFA γ-MSD $\beta$ | 0.54 [0.32, 0.69] | 0.78 | 0.67 [0.42, 0.82] | 0.87 | 0.60 [0.36, 0.75] | 0.84 |
| DFA γ-IR | 0.41 [0.19, 0.58] | 0.67 | 0.36 [0.13, 0.51] | 0.68 | 0.46 [0.26, 0.61] | 0.74 |
| DFA γ-DR | 0.32 [0.12, 0.49] | 0.62 | 0.57 [0.31, 0.74] | 0.77 | 0.39 [0.14, 0.59] | 0.66 |
| MSD β-IR | 0.39 [0.19, 0.52] | 0.71 | 0.34 [0.08, 0.49] | 0.65 | 0.43 [0.17, 0.58] | 0.73 |
| MSD β-DR | 0.30 [0.10, 0.45] | 0.64 | 0.60 [0.36, 0.73] | 0.86 | 0.21 [0.04, 0.39] | 0.58 |
| RMSF α-$AC_v(1)$ | 0.27 [0.06, 0.46] | 0.58 | 0.19 [0.03, 0.34] | 0.52 | 0.36 [0.11, 0.54] | 0.63 |

Linearity was rejected by Ramsey’s RESET test [100] in at least one scenario for three of the six couplings (DFA γ-MSD β, DFA γ-IR and MSD β-IR; Table S13 in S1 File). To describe the shape of these relations, four two-parameter candidate forms (straight line, one-term exponential, power law and logarithm) were fitted to each coupling in both directions and ranked by the corrected Akaike Information Criterion (AICc) [101–103]. The one-term exponential ranked first in 22 of the 36 contests (six couplings × three scenarios × two fit directions) and obtained the highest Borda count [104], the sum over the contests of the points awarded by rank (Materials and Methods), whereas the power law, first in only 2 contests, stayed closest to the best-ranked model on average (mean AICc difference from the best-ranked model, ΔAICc, of 1.23 against 1.85 for the exponential; Table S14 and Fig S1 in S1 File). Because no single form was decisively preferred, the couplings are reported by the strength and direction of their dependence rather than by the parameters of a fitted curve.

### Correlations among the systemic migratory-dynamics metrics

First, to assess the relation between two systemic hallmarks of migratory dynamics, long-term persistence and spatial-exploration efficiency, we analyzed the DFA scaling exponent γ together with the MSD diffusion exponent β (Fig 3). In Fig 3A, violin and box plots show the distribution, median and mean of both exponents in the three scenarios. In Fig 3B, the DFA γ (left axis) and MSD β (right axis) values of the 50 cells of each scenario are plotted against cell index, and the two traces rise and fall together. In all three scenarios, cells with stronger trend-reinforcing persistence (higher γ) were also more super-diffusive (higher β) (Fig 3C; Spearman ρ = 0.78 in the absence of stimuli, 0.87 under galvanotaxis and 0.84 under chemotaxis), and this was the strongest of the six couplings (*R*\*_n_ = 0.54 [0.32, 0.69], 0.67 [0.42, 0.82] and 0.60 [0.36, 0.75], respectively; Table 1). The coupling also has a precise theoretical reference. For an ideal self-similar signal with Hurst exponent H, which ranges from 0 to 1 and exceeds 0.5 for a persistent signal, the MSD exponent is β = 2H [105] and the DFA exponent of the radial distance is γ = H + 1 [92], so that β = 2(γ − 1) (dashed line in Fig 3C). This relation is not an identity: it holds only for a trajectory that scales in the same way across all the time scales analyzed, and the two exponents are estimated independently, DFA γ from the fluctuations of the radial distance and MSD β from the mean squared displacement. The cells lay close to this line: γ departed from the self-similar prediction β/2 + 1 by a median absolute difference of 0.026, 0.022 and 0.017 in the absence of stimuli, under galvanotaxis and under chemotaxis, respectively, smaller than the spread of γ between cells (interquartile range 0.078, 0.046 and 0.045). The DFA γ-MSD β coupling therefore shows that the long-term persistence and the super-diffusive exploration of the trajectories are two expressions of a common scale-free organization.

**Fig 3.**
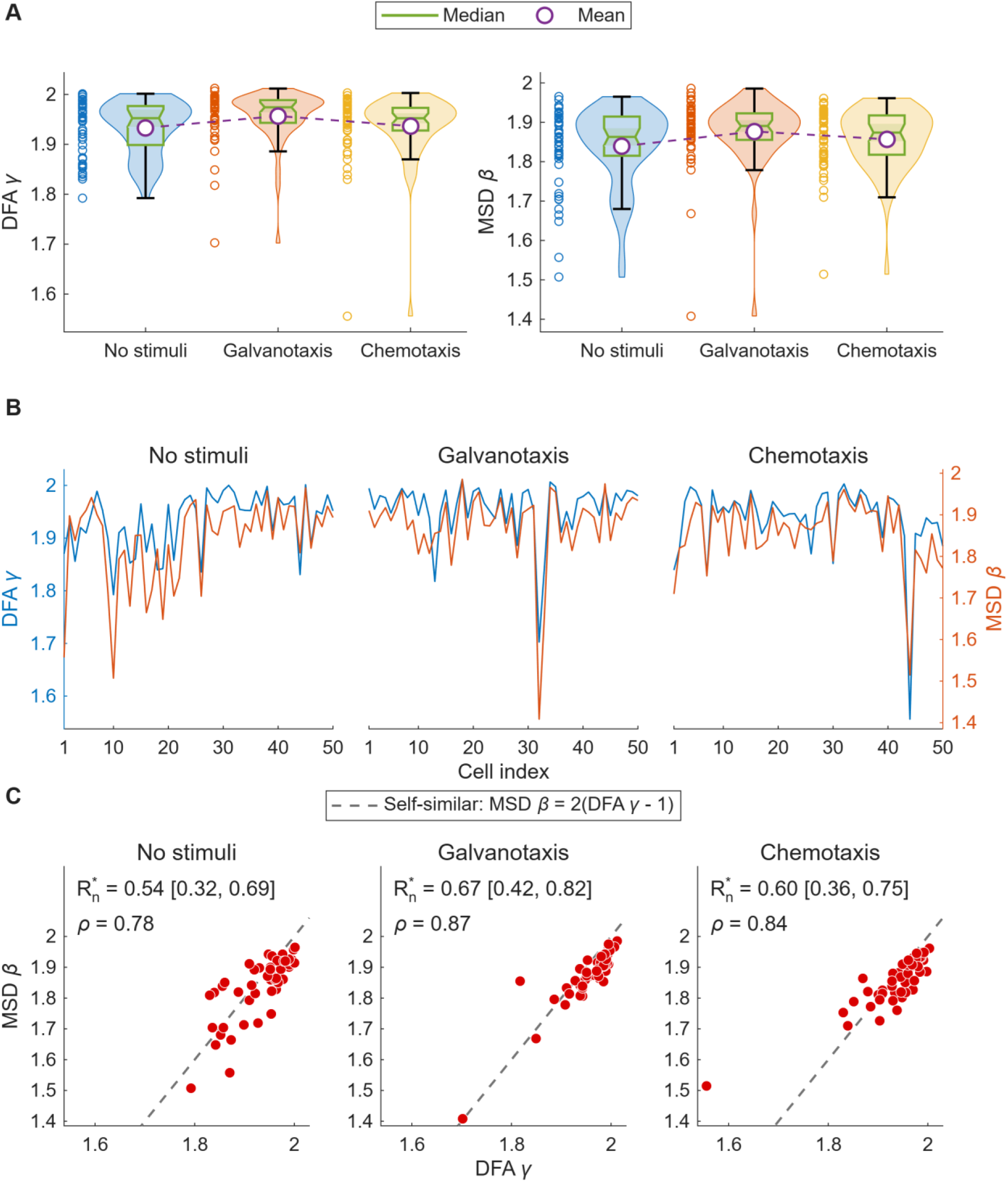
Relationship between the Detrended Fluctuation Analysis (DFA) scaling exponent γ and the Mean Squared Displacement (MSD) diffusion exponent β. **(A)** Violin plots (kernel density estimates within the observed range) and notched box plots of DFA γ and MSD β for the 150 cells studied under the three experimental scenarios; each box spans the interquartile range, its notch marks the 95% confidence interval of the median and its whiskers reach the most extreme values within 1.5 interquartile ranges of the box, the purple circles joined by a dashed line mark the means, and the open circles to the left of each violin are the individual cells. **(B)** DFA γ (left axis) and MSD β (right axis) of the 50 cells of each scenario plotted against cell index. **(C)** MSD β against DFA γ for each scenario, with the bias-corrected squared distance correlation *R*\*n (BCa 95% confidence interval) and the Spearman correlation ρ; axes are common across scenarios. The dashed line is the self-similar prediction MSD β = 2(DFA γ − 1).

Second, the RMSF exponent α was coupled to the speed memory AC_v_(1) (Fig 4). Cells whose move-step sequences carried stronger long-range correlations (higher α) also kept a more similar speed from one 5.5 s step to the next (higher AC_v_(1), stronger speed memory) in every scenario (Spearman ρ = 0.58 in the absence of stimuli, 0.52 under galvanotaxis and 0.63 under chemotaxis; *R*\*_n_ = 0.27 [0.06, 0.46], 0.19 [0.03, 0.34] and 0.36 [0.11, 0.54], respectively; Table 1). This coupling links a long-range property of the move-step sequence, which extends over minutes, to a short-term memory of speed between consecutive steps. It held at every sampling interval from 5.5 to 60 s (*R*\*_n_ ≥ 0.16 in every scenario; Fig S4E in S1 File).

**Fig 4.**
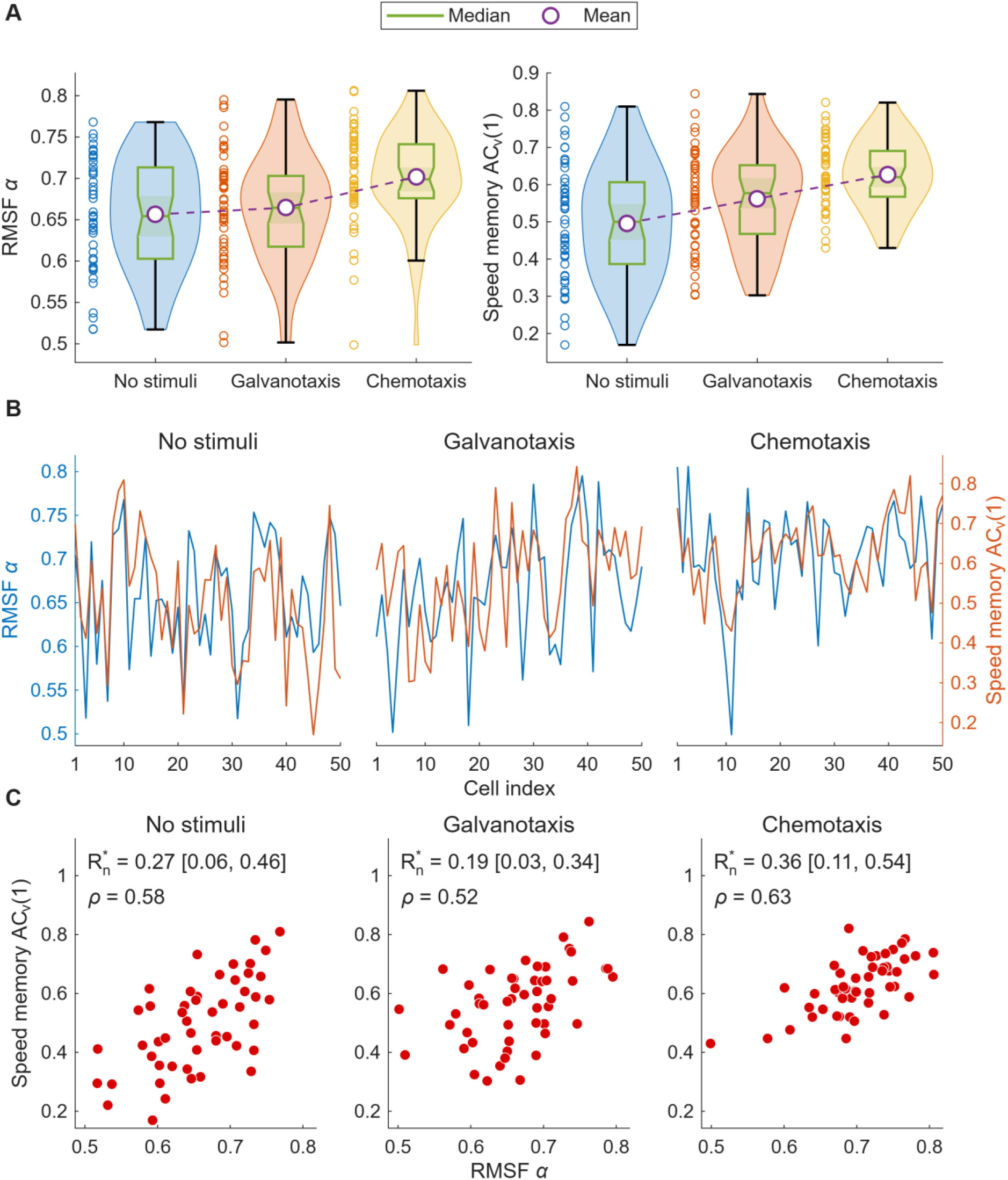
Relationship between the Root Mean Square Fluctuation (RMSF) exponent α and the speed memory ACv(1). **(A)** Violin and box plots of RMSF α and ACv(1) for the 150 cells studied under the three experimental scenarios, drawn as in Fig 3. **(B)** RMSF α (left axis) and ACv(1) (right axis) of the 50 cells of each scenario plotted against cell index. **(C)** ACv(1) against RMSF α for each scenario, with the bias-corrected squared distance correlation *R*\*n (BCa 95% confidence interval) and the Spearman correlation ρ; axes are common across scenarios.

### Correlations between systemic and kinematic dynamics

Both scaling exponents were coupled to the kinematic performance of the cells. First, to assess the relation between long-term persistence and the strength and straightness of the migratory response, we analyzed DFA γ together with IR (Fig 5) and with DR (Fig 6). In every scenario, cells with stronger trend-reinforcing persistence (higher γ) moved farther from their starting point (DFA γ-IR: Spearman ρ = 0.67 in the absence of stimuli, 0.68 under galvanotaxis and 0.74 under chemotaxis; *R*\* between 0.36 and 0.46) and along straighter paths (DFA γ-DR: ρ = 0.62, 0.77 and 0.66; *R*\* between 0.32 and 0.57) (Table 1).

**Fig 5.**
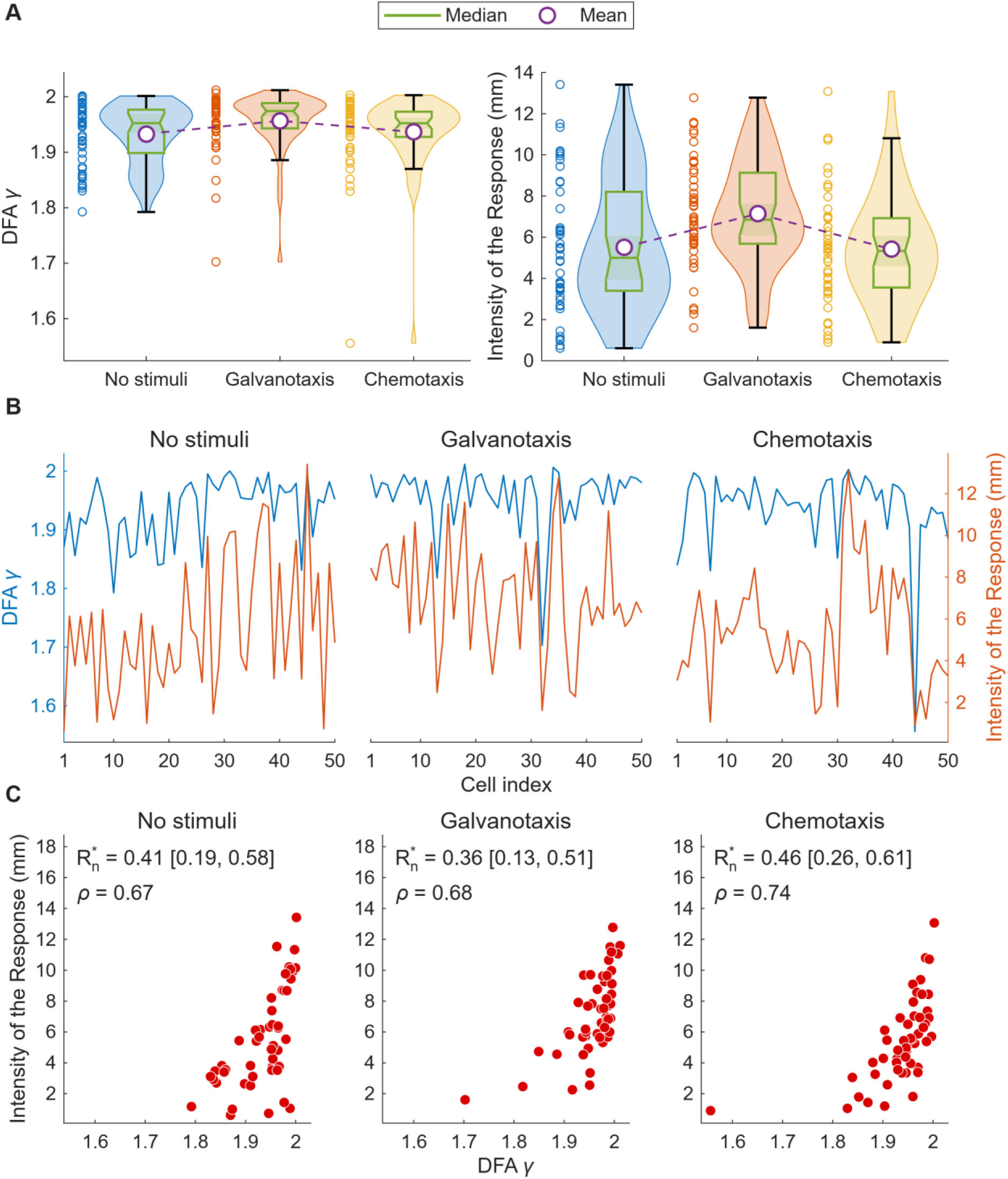
Relationship between the Detrended Fluctuation Analysis (DFA) scaling exponent γ and the Intensity of the Response (IR). **(A)** Violin and box plots of DFA γ and IR for the 150 cells studied under the three experimental scenarios, drawn as in Fig 3. **(B)** DFA γ (left axis) and IR (right axis) of the 50 cells of each scenario plotted against cell index. **(C)** IR against DFA γ for each scenario, with the bias-corrected squared distance correlation *R*\*n (BCa 95% confidence interval) and the Spearman correlation ρ; axes are common across scenarios.

**Fig 6.**
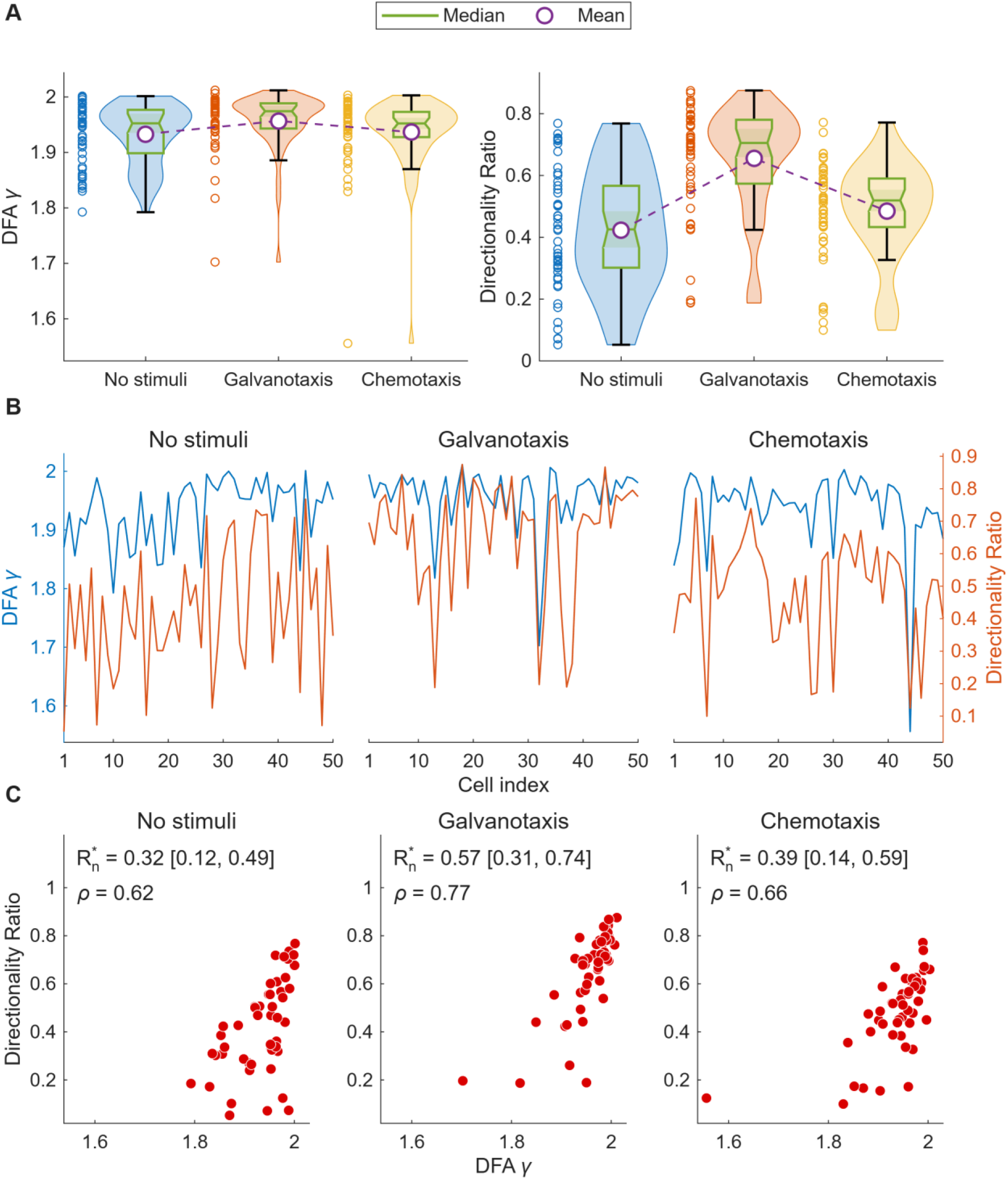
Relationship between the Detrended Fluctuation Analysis (DFA) scaling exponent γ and the Directionality Ratio (DR). **(A)** Violin and box plots of DFA γ and DR for the 150 cells studied under the three experimental scenarios, drawn as in Fig 3. **(B)** DFA γ (left axis) and DR (right axis) of the 50 cells of each scenario plotted against cell index. **(C)** DR against DFA γ for each scenario, with the bias-corrected squared distance correlation *R*\*n (BCa 95% confidence interval) and the Spearman correlation ρ; axes are common across scenarios.

Second, to assess the relation between spatial-exploration efficiency and kinematic performance, we analyzed MSD β together with IR (Fig 7) and with DR (Fig 8). More super-diffusive cells (higher β) likewise produced a stronger (MSD β-IR: ρ = 0.71, 0.65 and 0.73; *R*\*_n_ between 0.34 and 0.43) and more directed response (MSD β-DR: ρ = 0.64, 0.86 and 0.58; *R*\*_n_ between 0.21 and 0.60) in every scenario (Table 1). In Figs 5–8, panels A show the distributions of the two metrics, panels B their values cell by cell, and panels C their joint distribution on axes common to the three scenarios.

**Fig 7.**
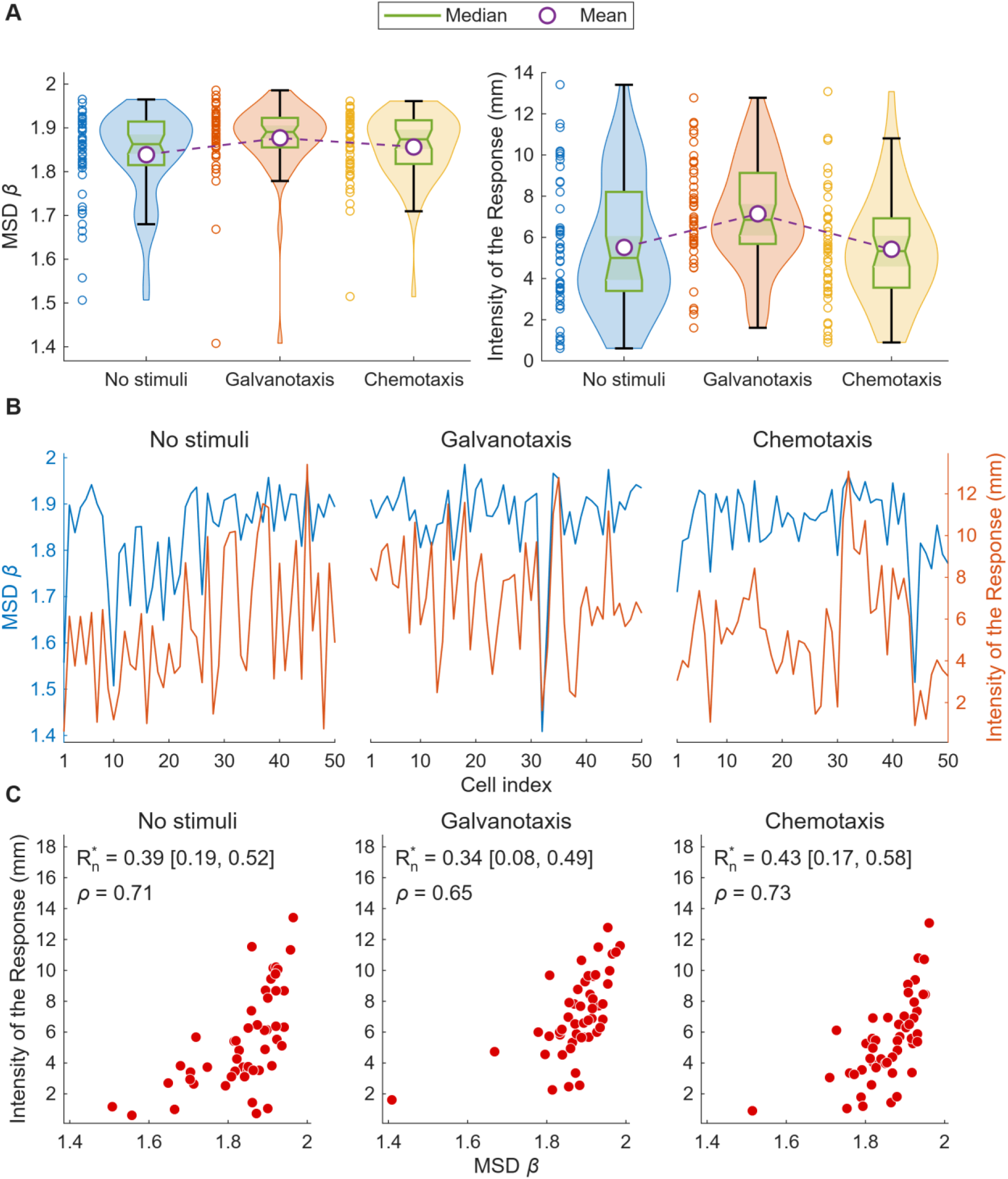
Relationship between the Mean Squared Displacement (MSD) diffusion exponent β and the Intensity of the Response (IR). **(A)** Violin and box plots of MSD β and IR for the 150 cells studied under the three experimental scenarios, drawn as in Fig 3. **(B)** MSD β (left axis) and IR (right axis) of the 50 cells of each scenario plotted against cell index. **(C)** IR against MSD β for each scenario, with the bias-corrected squared distance correlation *R*\*n (BCa 95% confidence interval) and the Spearman correlation ρ; axes are common across scenarios.

**Fig 8.**
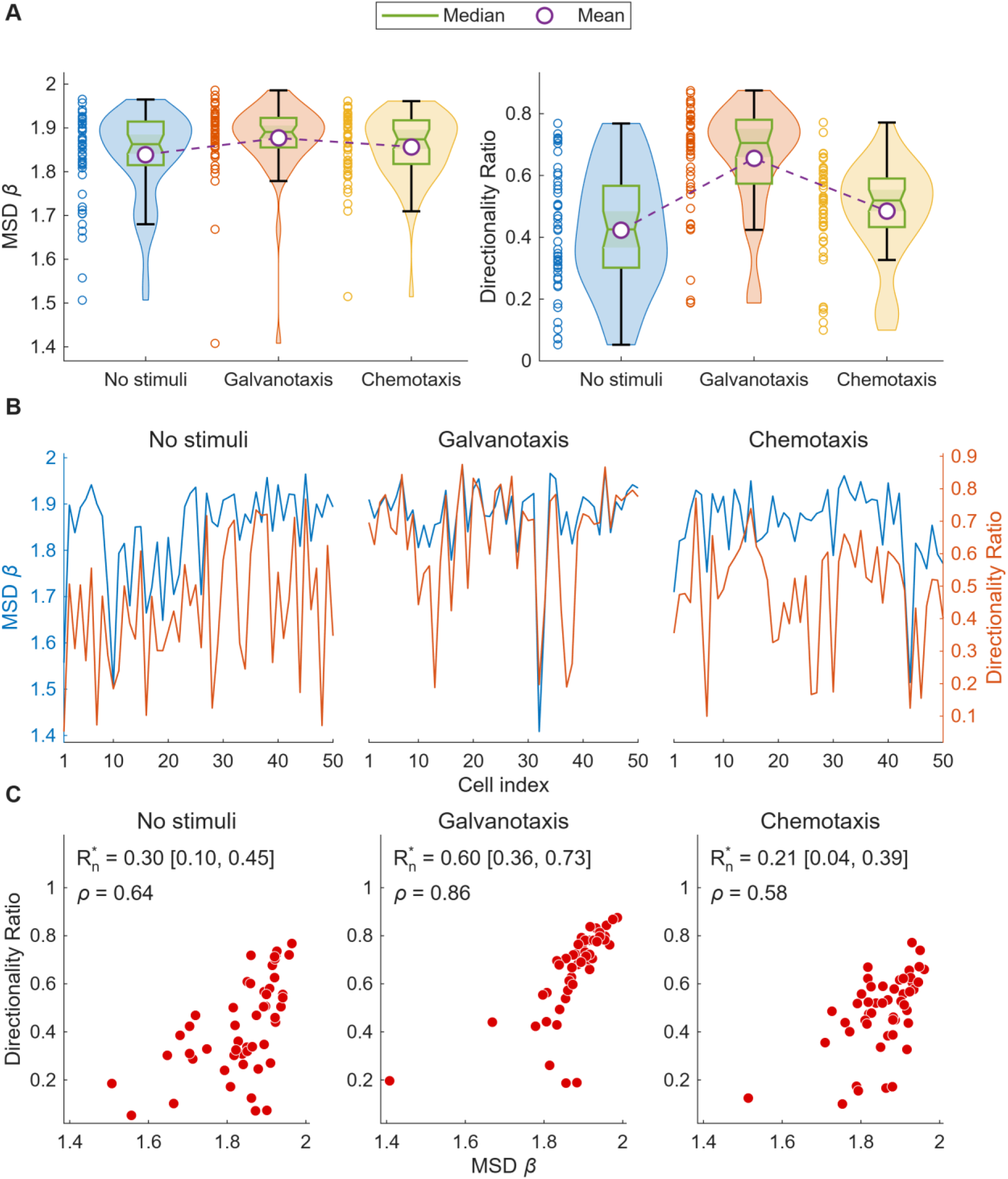
Relationship between the Mean Squared Displacement (MSD) diffusion exponent β and the Directionality Ratio (DR). **(A)** Violin and box plots of MSD β and DR for the 150 cells studied under the three experimental scenarios, drawn as in Fig 3. **(B)** MSD β (left axis) and DR (right axis) of the 50 cells of each scenario plotted against cell index. **(C)** DR against MSD β for each scenario, with the bias-corrected squared distance correlation *R*\*n (BCa 95% confidence interval) and the Spearman correlation ρ; axes are common across scenarios.

Thus, more persistent and more super-diffusive trajectories were also longer in net displacement and straighter in all three scenarios.

### External stimuli shift the metric values and part of their dependence structure, and preserve the six couplings

The stimuli shifted the values of six of the eleven metrics (Kruskal–Wallis test, q < 0.05; Table S12 and Figs S2 and S3 in S1 File). Effect sizes are given as ε^2^ for the Kruskal–Wallis test across the three scenarios and as Cliff’s δ for each pair of scenarios, δ being the probability that a cell of the first scenario has a higher value than a cell of the second minus the probability of the reverse, so that a negative δ indicates higher values in the second scenario. The largest effect was on the Directionality Ratio (ε^2^ = 0.268, q = 2 × 10^−8^), which was higher under galvanotaxis than both in the absence of stimuli (δ = −0.634, q = 2 × 10^−6^) and under chemotaxis (δ = 0.606 for galvanotaxis against chemotaxis, q = 3 × 10^−6^). The speed memory AC_v_(1) showed the second largest effect (ε^2^ = 0.138, q = 2 × 10^−4^) and was higher under chemotaxis than both in the absence of stimuli (δ = −0.515, q = 1 × 10^−4^) and under galvanotaxis (δ = −0.295, q = 0.037). Average Speed was highest in the absence of stimuli (ε^2^ = 0.099, q = 0.002), the RMSF exponent α was highest under chemotaxis (ε^2^ = 0.095, q = 0.002), and IR and DFA γ were highest under galvanotaxis (ε^2^ = 0.087, q = 0.003, and ε^2^ = 0.054, q = 0.032, respectively). MSD β, Approximate Entropy, the RMSF correlation time, t_θ_ and |C_θ-v_| did not differ between scenarios (q ≥ 0.141). The two cues therefore act preferentially on different features of locomotion: the electric field straightens the trajectory and lengthens the net displacement, whereas the chemical gradient makes the speed more persistent from step to step.

In contrast to these shifts, the six couplings were present in every scenario (Table 1). Their strength varied moderately between scenarios: the couplings of DR with the two scaling exponents were strongest under galvanotaxis (*R*\*_n_ = 0.57 for DFA γ-DR and 0.60 for MSD β-DR, against 0.21 to 0.39 in the other two scenarios), the scenario in which DR itself was highest, although their confidence intervals overlapped across scenarios (Table 1). A permutation test of the difference in *R*\*_n_ between scenarios (Materials and Methods) detected no difference for any of the six couplings (smallest q = 0.16, for MSD β-DR under galvanotaxis against chemotaxis; Table S15 in S1 File).

The same permutation test, applied to the 53 metric pairs not linked by definition in each of the three pairs of scenarios (159 contrasts; Table S15 in S1 File), detected three differences in dependence (q < 0.05), all of them involving the direction-memory time t_θ_. Under galvanotaxis, AS was independent of MSD β (*R*\*_n_ = −0.02, against 0.41 in the absence of stimuli and 0.37 under chemotaxis), whereas in the other two scenarios faster cells were more super-diffusive (ρ = 0.63 and 0.65), with between-scenario q = 0.052 and 0.080; the dependences of AS on DFA γ and on DR followed the same pattern without reaching significance (q between 0.087 and 0.15). Under chemotaxis, t_θ_ depended more strongly on MSD β than in the absence of stimuli (*R*\*_n_ =0.51 against 0.03, q = 0.016; 0.24 under galvanotaxis) and was coupled to AS (*R*\*_n_ = 0.44, against 0.00 in the absence of stimuli and −0.02 under galvanotaxis, q = 0.016 and 0.032), faster cells keeping their direction for longer (ρ = 0.72), a positive relation between speed and persistence of the kind reported across cell types by Maiuri et al. [68]. The contrasts that involve t_θ_ hold for the 5.5 s interval used throughout, because the dependences of t_θ_ change with the interval over which it is computed: at intervals of 15 s and longer, the *R*\*_n_ of t_θ_ with MSD β reached 0.3 to 0.7 in all three scenarios (Fig S4F in S1 File), and the *R*\*_n_ of t_θ_ with AS reached 0.28 to 0.38 in the absence of stimuli and 0.28 to 0.49 under chemotaxis but did not exceed 0.11 under galvanotaxis at any interval (Fig S4G in S1 File). The stimuli therefore left the six couplings intact but reorganized part of the dependence structure around speed: under the electric field, speed was independent of super-diffusive exploration and at most weakly dependent on the persistence of direction at every time scale, whereas the chemical gradient made speed and the persistence of direction dependent already at the 5.5 s interval, at which they were independent in the absence of stimuli.

## Discussion

The central result of this study is that the systemic and kinematic descriptors of Amoeba proteus locomotion are not independent observables but vary together in every environment. Of the 55 metric pairs, six were dependent in all three scenarios besides the pairs linked by definition, and all six were positive and monotonic: cells with stronger long-term persistence (higher DFA γ) were more super-diffusive (higher MSD β), both exponents rose with the net displacement (IR) and the straightness (DR) of the trajectory, and cells whose move-step sequences carried stronger long-range correlations (higher RMSF α) also retained more memory of their speed from one step to the next (higher AC_v_(1)). The strongest of these couplings, between DFA γ and MSD β, followed the relation expected for a self-similar trajectory, in which both exponents are set by the same Hurst exponent. Because DFA γ and MSD β were estimated independently, from the fluctuations of the radial distance and from the mean squared displacement, their agreement with this relation indicates that long-term persistence and super-diffusive exploration are two expressions of a common scale-free organization of the trajectories.

The couplings of the two scaling exponents with IR and DR have no such exact theoretical relation. A persistent, super-diffusive trajectory accumulates net displacement faster than a diffusive one, but the scaling exponents are computed from how fluctuations and displacements grow across time scales and do not depend on the overall magnitude of the displacement, whereas IR measures that magnitude directly and DR the fraction of the path that contributed to it. That cells with higher scaling exponents also traveled farther and straighter in every scenario therefore links the organization of the trajectory across time scales to its migratory output. This link is consistent with the theoretical prediction that temporal correlation increases the persistence factor, the ratio of net displacement to path length that we call DR, at every level of angular diffusion tested [106]. Approximate Entropy, in contrast, was not dependent on any other metric in all three scenarios; it depended only on the speed and direction memories, and only in the absence of stimuli (Table S13 in S1 File). Because ApEn was not dependent on any scaling exponent or kinematic descriptor in any scenario, carried no weight on the first principal component, which DFA γ, MSD β, IR and DR dominate, and joined the other metrics only near the top of the dependence dendrogram, the tree that joins first the metrics with the highest mean *R*\*_n_, so that a metric joining near the top depends only weakly, on average, on the others (Materials and Methods; Table S13 and Figs S7 and S8 in S1 File), the regularity of the move-step sequence is a property of migration that persistence, exploration and displacement do not capture.

The coupling between RMSF α and the speed memory AC_v_(1) links two properties of the speed of the cell that are measured on different time scales: the long-range correlations of the move-step sequence, which extend over minutes, and the persistence of speed between consecutive 5.5 s steps. Both can arise from a single source. When the fluctuations of the move steps are positively correlated, as in the fractional Brownian motion that drives translational motion in the model of Montenegro-Rojas et al. [106], a single parameter, the Hurst exponent H, sets the correlation between steps at every lag, and for H > 0.5 that correlation increases with H both between consecutive steps and between steps far apart in time [105]. A stronger long-range correlation of the move steps (higher RMSF α) is then accompanied by a stronger short-term memory of speed (higher AC_v_(1)), as observed in every scenario. Of the three components into which the velocity memory can be separated [82], only the speed memory was coupled to another metric in all three scenarios, which supports treating speed memory and direction memory as distinct properties of migration, as proposed by Shaebani et al. [82].

Maiuri et al. (2015) [68] reported exponential coupling between cell speed and persistence; however, both quantities in their analysis are kinematic. The present work extends the study of couplings between migratory properties to systemic-kinematic pairs, in which one variable measures the scaling of the trajectory dynamics (DFA γ, MSD β) and the other a kinematic property of the trajectory (IR, DR), and to a pair of systemic properties measured on different time scales (RMSF α and AC_v_(1)). The one-term exponential, the form of the speed-persistence coupling [68], ranked first most often among the candidate forms fitted to these couplings, but no form was decisively preferred; we have therefore described the couplings by their strength and direction rather than by fitted parameters.

The three scenarios are also informative when compared with one another. Directional cues do not shift every descriptor in the same way: the electric field produced the largest change in trajectory straightness, with the directionality ratio highest under galvanotaxis (Cliff’s δ = −0.634 against the absence of stimuli), whereas the chemical gradient produced the largest change in the speed channel, with the lag-one speed autocorrelation highest under chemotaxis and significantly higher than in the absence of stimuli (δ = −0.515) and under galvanotaxis (δ = −0.295). The two cues therefore act preferentially on different components of the locomotion output, one on direction and the other on the regularity of speed. Where coupled metrics shifted, they shifted together, in the direction of their positive coupling: DFA γ, IR and DR were all highest under galvanotaxis, and RMSF α and AC_v_(1) were both highest under chemotaxis. The couplings themselves were present in all three scenarios and only their strength varied, the couplings of DR with the two scaling exponents being strongest under galvanotaxis, the scenario in which the trajectories were straightest, with overlapping confidence intervals and no significant difference in the between-scenario permutation test of *R*\*_n_ (Materials and Methods; smallest q = 0.16; Table S15 in S1 File). Directional cues therefore move cells along the couplings rather than altering the couplings themselves. Outside the six couplings, however, the cues did change how speed depended on the other descriptors: under galvanotaxis the Average Speed was independent of MSD β, to which it was coupled in the other two scenarios, consistent with the independent regulation of speed and directionality reported for electrotaxis in *Dictyostelium* [107], and the coupling between speed and persistence that Maiuri et al. described as universal [68] was present under chemotaxis already at the 5.5 s interval and in the absence of stimuli at intervals of 15 s or more, but remained weak under galvanotaxis at every interval (*R*\*_n_ ≤ 0.11; Table S15 and Fig S4G in S1 File). In the model of Montenegro-Rojas et al., a directional cue acts only on the orientation of the cell and leaves the temporal correlation of its motion unchanged, yet orients migration toward the target [106]. The scaling exponents measured under a cue may therefore include directionality imposed by the cue, as the higher DFA γ under galvanotaxis suggests; because the same couplings were present in the absence of any stimulus, however, they cannot be attributed to the cue.

Beyond the specific pairs analyzed, the picture that emerges is one in which locomotion behaves as an emergent systemic property governed by the functional integration of most, if not all, cellular components, and not as a behavior attributable to any singular molecular part, partial mechanism or individual process. Cells sustain this organization far from thermodynamic equilibrium, where irreversible metabolic processes and the permanent dissipation of energy allow the system to become spatially and temporally self-organized, giving rise to the ordered collective patterns and the long-range correlations that characterize dissipative structures, and the same principles have been shown to govern the cellular metabolic structure and its information-processing activity [43,44]. The present results are a further step along that line of inquiry: having established that migratory trajectories are regulated at the systemic level in nucleated cells, and that such regulation survives even the removal of the nucleus [45,46], we now find that the descriptors of those trajectories are themselves bound to one another by statistical dependences that hold in every environment tested. These couplings provide a quantitative signature of the complex functional integration that emerges in the mechanisms driving cell migration. Overall, the systemic and kinematic signatures of migration constitute an integrated system whose couplings are preserved across environments.

## Materials and Methods

### Cell cultures

*Amoeba proteus* (Carolina Biological Supply, Burlington, NC, USA; #131306) were grown in Ø100 × 20 mm Petri dishes (Corning® CLS430167) filled with Chalkley’s simplified medium [51] (NaCl, 1.4 mM; KCl, 0.026 mM; CaCl2, 0.01 mM) and sterilized wheat grains.

### Cell preparation

Before all experiments, amoebae were starved for 24 hours in clean, axenic Chalkley’s simplified medium without any external stimuli. For each experiment, only actively moving cells that displayed a healthy spindled or oval morphology were selected. Selected cells were washed in fresh Chalkley’s medium before being placed into the experimental chamber.

### Experimental set-up

Experiments were conducted using a custom-designed device composed of one or two standard electrophoresis blocks (Mini-Sub cell GT, Bio-Rad, Hercules, CA, USA) and a custom-made glass observation chamber (see Fig 2).

For galvanotaxis experiments, two electrophoresis blocks were used. The first block was connected to a power supply (BIO-RAD Model 1000/500), and the second block (from which the wires reaching from the electrode pins down to the wells were removed) was connected to the first via two 12 cm agar bridges (2% agar in 0.5 N KCl). A custom-made glass observation chamber was placed on the elevated platform of this second block. This two-block configuration prevented direct contact between any electrodes or metallic wires and the experimental medium, avoiding contamination of the medium where amoebae migrated.

For migration in the absence of stimuli and chemotaxis experiments, a single electrophoresis block was utilized. The glass observation chamber was placed directly onto its central elevated platform. For both galvanotaxis experiments and experiments in the absence of stimuli, an electrophoresis block that had never been in contact with the chemotactic peptide was used to prevent contamination.

### Experimental chamber

The observation chamber was constructed from a modified glass slide and carefully trimmed standard glass coverslips. Two glass strips (24 × 40 × 0.17 mm each) were glued with silicone to a standard (75 × 25 × 1 mm) glass slide to act as the lateral walls of the chamber. A central glass piece (3 × 24 × 0.1 mm) and two lateral sliding pieces (40 × 24 × 0.1 mm) were placed on top. This configuration allowed for the creation of a laminar flow when closed and for the placement and extraction of cells when open. The modified glass slide was carefully washed and new trimmed coverslips used for each experiment.

### Migration in the absence of stimuli

Amoebae were placed in 30 µL of clean Chalkley’s medium under the central glass piece within the glass chamber positioned on a clean electrophoresis block. The cells were allowed to adapt and adhere to the glass surface for approximately 2 minutes before the chamber was sealed to create a laminar flow. Cell movements were then recorded.

### Galvanotaxis

Amoebae were placed in the glass chamber, which was positioned on the second (the one not in direct contact with the power supply unit) of the two electrophoresis blocks connected by agar bridges. An external direct-current electric field was applied to the first block, generating a voltage that oscillated between 58.5 and 60 V (334–343 mV/mm) inside the experimental glass chamber. To ensure precise control, a 1 MΩ variable resistor and a microammeter were connected in series with the power supply (see Fig 2A). The current was maintained between 70 and 74 µA by adjusting the variable resistance. The power was turned off after 30 minutes of cell exposure.

### Chemotaxis

The experimental chamber was placed atop the central platform of an electrophoresis block. After establishing a laminar flow, 750 µL of 2 × 10^−4^ M N-Formylmethionyl-leucyl-phenylalanine (nFMLP) peptide solution (Sigma-Aldrich, St. Louis, MO, USA; #F3506) was added to the 75 ml of Chalkley’s medium in the well corresponding to the positive pole (left well in Fig 2D and the recordings), achieving a final working concentration of 2 × 10^−6^ M. The medium was gently mixed until amoebae began to respond, at which point cell movements were recorded for 30 minutes.

### Chemotactic gradient

We evaluated how the nFMLP gradient concentration evolved over time. The assay used 4 µM fluorescein-tagged peptide (formyl-Nle-Leu-Phe-Nle-Tyr-Lys, fluorescein derivative; Invitrogen, #F1314), loaded in place of nFMLP. In this assay, 60 µL aliquots of medium were withdrawn at a series of time points (0, 2, 5, 10, 15, 20, and 30 min) after the laminar flow was established (Fig S5 in S1 File). Each aliquot was drawn from the middle of the experimental glass chamber through a narrow opening created by slightly displacing one of the side glass pieces surrounding the central one. To establish the nFMLP concentration corresponding to every time point, we compared the readings against a standard curve built from established concentration values of fluorescein-labeled peptide. At each time point we collected two aliquots, and because the whole experiment was run three separate times, the assay yielded a total of six measurements per time point. We quantified fluorescence on a SynergyHTX plate reader (BioTek) using black, glass-bottomed 96-well plates (Cellvis, #P96-1.5H-N), with excitation and emission set to 460 and 528 nm, respectively, in accordance with the protocol described by Green and Sambrook [108].

### Track recording and digitizing

Cellular trajectories were recorded using a digital camera attached to an AmScope SM-2T stereomicroscope (AmScope, Irvine, CA, USA). Videos were captured for a duration of 30 minutes, resulting in 3600 frames per recording (two frames per second). Acquiring frames more often than the analysis needs and skipping the extra ones afterwards is the safer strategy for tracking cell migration, since frames can be dropped but not added [72]. The movements of individual amoebae were manually tracked from the recorded videos using the TrackMate software plugin for ImageJ [109] as described by Hilsenbeck et al. [110], as automated cell tracking is prone to errors [110], requiring post-hoc manual curation [111].

### Quantitative analyses

All analyses were run in MATLAB R2026a [112] with the Statistics and Machine Learning Toolbox [113] and the Curve Fitting Toolbox [114], except the two-lines test, which was run in R 4.6.1 [115] with the author’s R package statuser 0.4.0 [99,116] and the packages mgcv [117], sandwich [118] and lmtest [119].

### Root Mean Square Fluctuation (RMSF)

The root mean square fluctuation (RMSF) analysis is a traditional approach in Statistical Mechanics built upon the concepts introduced by Gibbs in 1902 [94] and Einstein in 1909 [95]. It was later expanded and used to measure physiological signals (as per [120,121]). We employed the RMSF method to evaluate the existence of power-law autocorrelation in the time-series of move-step fluctuations, adhering to the procedure outlined by Viswanathan et al. in 1996 [122]. To calculate the RMSF in a given two-dimensional trajectory *P*(*t*) = [*x*(*t*), *y*(*t*)] with values equidistant in time, we first defined the displacement time series as: 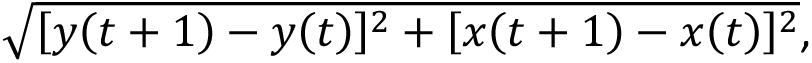, for *t* = 1,…,*t_max_* − 1. From here, the distance traveled after l move steps was computed as 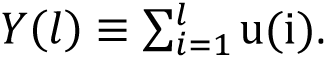. Finally, the RMSF of the average displacement is calculated as 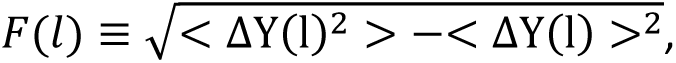, where we have defined *ΔY*(*l*) ≡ *Y*(*l* + *l*_0_) − *Y*(*l*_0_) and the brackets denote the average over all possible values of *l_0_*. Therefore, *F*(*l*) is defined as the square root of the difference between the average of the square of ΔY(l) minus the square of its average. Persistence can be identified when the fluctuations follow a power law, i.e., *F*(*l*)∼*l^α^*. For uncorrelated data, the fluctuation exponent *α* is equal to 0.5, whereas a value between 0.5 and 1 signifies the presence of positive persistence [122]. The RMSF correlation time, the time during which the regime of long-range correlations is preserved, was assessed as the maximum value of l at which the scaling F(l) started to curve in a log-log representation [62].

### Mean Squared Displacement (MSD)

The Mean Squared Displacement (MSD) method was first introduced by Einstein [88] and has been used in multiple fields, including the quantification of cell migration [72,123]. MSD calculates the average squared displacement in a migration path over increasingly longer time scales [72], and provides a metric for the spatial extent and efficiency of cell migration [87]. For a two-dimensional trajectory *P*(*t*) = [*x*(*t*), *y*(*t*)], the MSD is defined as

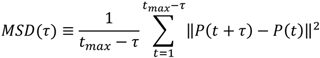

where ‖·‖ denotes the Euclidean norm and *τ* represents the time scale. Here, diffusion was studied up to a maximum time scale equal to one quarter of the data size. Random walks are characterized by their power-scaling relationship MSD(*τ*) ∝ *τ^β^*, where *β* characterizes diffusion regimes: *β* = 1 corresponds to classical Brownian motion, *β* > 1 indicates super-diffusive (persistent) migration, and *β* < 1 denotes sub-diffusive (restricted) motion.

### Detrended Fluctuation Analysis (DFA)

Detrended Fluctuation Analysis (DFA) is a widely adopted method for uncovering power-law autocorrelations in nonstationary time series, introduced by Peng and colleagues in 1994 to detect power-law autocorrelations in time series [90]. It has been intensively used in the quantitative analysis of physiological signals [91]. For the radial distance of the cell, R(t) = √(x(t)^2^ + y(t)^2^), measured from the cell’s initial position (*x*_0_, *y*_0_) such that each trajectory effectively starts at the origin ((0,0)), the signal profile is calculated by determining the series’ cumulative sum, represented by 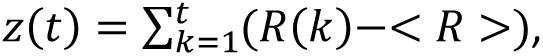 where brackets indicate the average of R(k). Next, the time series *z*(*t*) is divided into boxes of length *n*, and the local trend *z*_n_(*t*) is subtracted from each box. The boxes were cut from both ends of the series, so that no part of the record is left out [124].

Detrended signal fluctuation is then calculated by 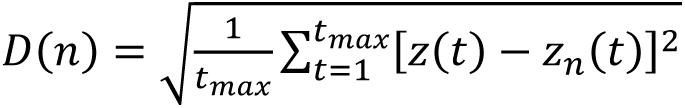 for all box sizes. D(n) is the fluctuation function and n the box size. The existence of power-law autocorrelations is revealed by a linear relationship on a log-log graph *D*(*n*)∼*n^γ^*.For fractional Brownian motion, the non-stationary class of signals to which the radial distance *R*(*t*) belongs, with Hurst exponent *H* = γ – 1 [92], the exponent γ denotes anti-persistence when 1 < γ < 1.5, absence of persistence when γ ≈ 1.5, and positive persistence (trend-reinforcing behavior) when 1.5 < γ < 2 [92].

### Approximate Entropy

Approximate Entropy (ApEn) is a metric introduced by Pincus and colleagues in 1991 [85] to measure time series regularity. Here it was computed on the move-step lengths of each trajectory subsampled every 5.5 s (Velocity-memory metrics). Low ApEn is indicative of a highly regular signal that can be stored in fewer bits of information, while a highly unpredictable and irregular signal has high ApEn and requires more bits of information to be stored. Given a time series of *N* equally spaced values *S* = *s*(1), *s*(2),…,*s*(*N*), which can be of relatively small size (N > 100) [83], fix a positive number *r* (specifies a filtering level, a common choice is *r* = 0.2 · *SD*, where *SD* is the standard deviation of the signal) and an integer number *m*. We used *m* = 2, the length of compared runs at which ApEn can be estimated from relatively few points [83]. Form a sequence of vectors *v*(1), *v*(2),…,*v*(*N* − *m* + 1) in ℝ*^m^*, real m-dimensional space, defined by *v*(*i*) = [*s*(*i*),…,*s*(*i* + *m* − 1)]. For each *i* = 1,…,*N* − *m* + 1, the sequence *v*(1), *v*(2),…,*v*(*N* − *m* + 1) is used to construct

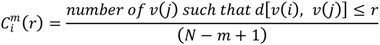

where *d* [*v*(*i*), *v*(*j*)] is defined as 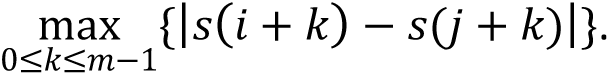.

Thus, *d* represents the distance between the scalar components of the vectors *v*(*i*) and *v*(*j*). Then, the variable 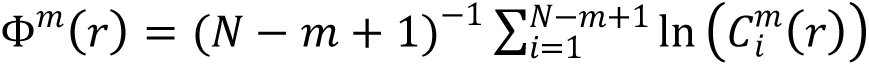 is established, where ln is the natural logarithm. Finally, Approximate Entropy is defined by *ApEn*(*S*, *r*, *m*) = Φ*^m^*(*r*) − Φ*^m^*^+1^(*r*) for fixed *m* and *r*.

### Kinematic metrics

Three complementary descriptors quantified each trajectory’s geometry. Let *P*(*t*) be the cell’s position vector at time *t*, sampled at equally spaced points *P*_0_,…,*P_N_* over the trajectory duration *T*, and let ∥⋅∥ denote the Euclidean norm (modulus). The intensity of the response (IR) reflects the space covered by the cell; we computed the modulus of the trajectory, that is, the net distance (in mm) between its origin *P*(0) and endpoint *P*(*T*), *IR* =∥ *P*(*T*) − *P*(0) ∥, as a measure of the magnitude of the response. The directionality ratio (DR) is the net displacement divided by the total path length, *DR* = *IR*/*L*, where 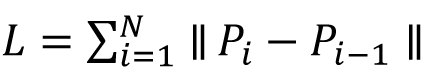 is the path length summed over consecutive sampled positions *P_i_*; DR ranges from 0 (highly tortuous) to 1 (perfectly straight). The average speed (AS) is the path length per unit time, *AS* = *L*/*T*, expressed in micrometers per second. Because *IR* = *T* ⋅ *DR* ⋅ *AS* up to unit conversion (here *IR* = 1.8 *DR* ⋅ *AS*, with *T* = 1800 s and the millimeter-to-micrometer factor), the pairs IR-DR and IR-AS are definitional and were excluded from the coupling analysis. IR, DR and AS depend only on the set of step increments of the trajectory, the displacement vectors between consecutive recorded positions, and not on their order: any order of the same increments gives the same endpoint, and hence IR, and the same path length L, and hence DR and AS. Every other metric, including the velocity-memory metrics computed after subsampling, depends on the order in which the steps occur; this is the criterion by which the metrics are classed as kinematic or systemic.

### Velocity-memory metrics

Following Shaebani et al. (2022) [82], we separated the velocity autocorrelation into speed-memory and direction-memory components plus a speed-direction coupling, each evaluated at a coarse-graining interval *Δt*. For each step *t* we defined the instantaneous speed *v_t_* =∥ *P_t_* − *P_t_*_−1_ ∥/*Δt* and the local heading *θ_t_* (the direction angle of the step); ⟨⋅⟩ denotes the average over the trajectory, *σ_v_* and *σ_θ_* the standard deviations of speed and heading, and *σ*^2^ the speed variance. The speed memory is the lag-one speed autocorrelation, 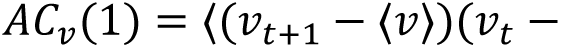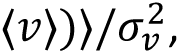 bounded in [−1,1]. The direction memory is a persistence time *t*_θ_ obtained from the mean cosine of the turning angle *α_i_* = *θ_i_* − *θ_i_*_−1_: assuming exponential directional decay over the interval Δ*t* between consecutive headings [125,126], ⟨⟩ = *e*^−*Δt*/*t*^*^θ^*, we solved *t_θ_* = −*Δt*/*ln*⟨⟩ (in seconds). The speed-direction coupling is the cross-correlation *C_θ_*_−*v*_ = ⟨(*θ* − ⟨*θ*⟩)(*v* − ⟨*v*⟩)⟩/(*σ_θ_σ_v_*), where ⟨*θ*⟩ is the circular mean heading [127], *v* − ⟨*v*⟩ is wrapped to [−π, π), and *σ*_θ_ is the standard deviation of *v* − ⟨*v*⟩; it is bounded in [−1,1]. A mirror reflection of a trajectory reverses the sign of C_θ-v_ and leaves the other metrics unchanged; the distance-covariance test detects dependence on the magnitude of C_θ-v_ as well as on its sign, and the scenarios were compared on |C_θ-v_|. In manual tracking the recorded position is that of the clicked pixel, so a cell that moves less than one pixel between frames, or idles, keeps its coordinates and yields a zero-length step, the pixel grid effect [72]. ApEn, *AC_v_*(1), *t_θ_* and *C_θ_*_−*v*_ were computed on trajectories subsampled to one position every 5.5 s (the subsampling procedure of Gorelik and Gautreau [72]), the shortest interval at which more than 99% of steps were displacements; all other metrics use every frame. At 0.5 s, 56 to 58% of the steps have zero length; the metrics that use every frame were nevertheless robust to them: recomputed from the positions every 5.5 s, the path length was a median 13 to 15% shorter but ordered the cells in the same way for DR and AS (Spearman ρ ≥ 0.97), the dependence of DR on DFA γ and on MSD β changed by at most 0.03 in *R*\*_n_, and RMSF α correlated with its value at 0.5 s (ρ = 0.79 to 0.91) and kept its dependence on AC_v_(1) (*R*\*_n_ = 0.36, 0.18 and 0.35, against 0.27, 0.19 and 0.36). Zero-length steps have no heading and were left out of t_θ_ and C_θ-v_ [72], so that each turning angle joins two consecutive steps that moved. The dependence of ApEn, AC_v_(1), t_θ_ and C_θ-v_, and of the dependences of AC_v_(1) and t_θ_, on the sampling interval Δt is shown in Fig S4 in S1 File, and the per-trajectory values of AC_v_(1), t_θ_ and C_θ-v_ are given in Tables S6, S7 and S8 in S1 File.

## Statistical analysis

Because the Lilliefors test rejected normality in 12 of the 33 metric and scenario combinations (p < 0.05), and in 10 after Benjamini–Hochberg correction (p-values in the last row of Tables S1 to S11 in S1 File; values beyond the tabulated range of 0.001 to 0.5 are given as bounds), we reported all results as the median and interquartile range (median/IQR). The Spearman correlation ρ is reported as the descriptive effect size of each coupling (Table 1), and the Pearson correlation only in the Supporting Information (Table S13 and Fig S6 in S1 File); neither was used to decide statistical dependence. Statistical dependence was tested for each of the 55 pairs among the eleven metrics, within each scenario, with the distance covariance test of independence [96], whose population statistic is zero only under independence, so that linear, nonlinear and non-monotonic dependence are all detected. Its null distribution was obtained from 9,999 random permutations of one metric of the pair, and the p-value was computed as p = (1 + B)/(9,999 + 1), where B is the number of permuted statistics at least as large as the observed one [128]. The 165 p-values (55 pairs × 3 scenarios) were corrected with the Benjamini–Hochberg procedure [129], and a pair was considered dependent when its corrected value q was below 0.05 in all three scenarios (Table S13 in S1 File). The strength of each dependence was measured by the bias-corrected squared distance correlation *R*\*_n_ [97], an estimator of the squared population distance correlation that is close to 0 under independence and can take small negative values, with its bias-corrected and accelerated (BCa) bootstrap 95% confidence interval from 2,000 resamples of cells [130,131]. The pairs IR-DR and IR-AS are fixed by the identity IR = 1.8 × DR × AS (Kinematic metrics) and were treated as definitional.

The dependence structure of the eleven metrics is also displayed in the Supporting Information by Pearson and Spearman correlation matrices (Fig S6 in S1 File), principal component analysis (Fig S7 in S1 File), a dendrogram of the metrics (Fig S8 in S1 File), matrices of *R*\*_n_ and of the Spearman correlation (Fig S9 in S1 File), and a forest plot of *R*\*_n_ with its BCa intervals for the 15 non-definitional pairs with the largest median *R*\*_n_ (Fig S10 in S1 File). Differences in each metric between the three scenarios were tested with the Kruskal–Wallis test [132], with the effect size ε^2^ = H/(n − 1), where H is the Kruskal–Wallis statistic and n = 150 the number of cells [133], and between pairs of scenarios with the Mann–Whitney U test [134] and Cliff’s δ [135], the probability that a cell of the first scenario has a higher value than a cell of the second minus the probability of the reverse, with BCa bootstrap confidence intervals, from 2,000 resamples of the cells of each scenario. P-values were corrected with the Benjamini–Hochberg procedure separately over the 11 Kruskal–Wallis and the 33 pairwise tests (Table S12 and Figs S2 and S3 in S1 File). The confidence intervals of δ were computed at the level 1 − 0.05 × 12/33 = 98.2%, 12 being the number of the 33 pairwise tests with q < 0.05; this level keeps the false coverage rate, the expected proportion of intervals that miss the true δ among those of the contrasts with q < 0.05, at or below 0.05 [136].

### Differences in dependence between scenarios

For each of the 53 metric pairs not linked by definition and each of the three pairs of scenarios, the difference between the *R*\*_n_ values of the two scenarios was tested by permutation. Each metric was first standardized to zero mean and unit variance within each scenario; this leaves *R*\*_n_ unchanged, because *R*\*_n_ does not depend on the location or the scale of either metric [96], but removes the differences in metric values between scenarios (Table S12 in S1 File) from the permuted sample. The 100 cells of the two scenarios were then pooled and randomly split into two groups of 50 in each of 9,999 permutations, and the p-value was computed as p = (1 + B)/(9,999 + 1), where B is the number of permutations in which the absolute difference between the *R*\*_n_ values of the two groups was at least as large as the observed one [118]. The 159 p-values (53 pairs × 3 pairs of scenarios) were corrected with the Benjamini–Hochberg procedure [119] (Table S15 in S1 File). Standardization removes differences in location and scale but not in the shape of the distributions, so a significant contrast can also reflect a change in the shape of a metric between scenarios.

### Principal component analysis (PCA)

To assess the effective dimensionality of the metric set, the eleven metrics were z-scored across the 150 cells (pooled over the three experimental scenarios) and decomposed by PCA on their correlation matrix R. Standardization is required at this step because the eleven metrics carry different units and variances, so an unscaled decomposition would be dominated by whichever metric happens to have the largest spread. PCA solves the eigenproblem R vₖ = λₖ vₖ, where λₖ is the variance carried by the k-th principal component and vₖ is its loading vector, the vector of weights of the eleven metrics on that component. The proportion of variance explained by component k is λₖ / Σⱼ λⱼ. Effective dimensionality was summarized by the participation ratio PR = (Σₖ λₖ)² / Σₖ λₖ², which is 1 when a single component dominates and equals the number of metrics when all contribute equally [137]. The contribution of metric m to component k was taken as the squared loading vₘₖ², which sums to one over the metrics within each component, and its overall contribution to the retained subspace as the variance-weighted mean of these squared loadings over the first three components (Fig S7 in S1 File). PCA was used descriptively: it is linear, and pooling the scenarios mixes differences between cells with differences between scenarios, so the dependence results rest on the distance covariance tests. The participation ratio was also computed within each scenario.

### Hierarchical clustering of the metrics

To display the dependence structure as a tree, the metrics were grouped by their pairwise statistical dependence rather than by linear correlation. For each metric pair (m, n) the bias-corrected squared distance correlation *R*\*_n_ was computed in each experimental scenario and averaged across the three experimental scenarios to give *R̄*ₘₙ, from which a distance dₘₙ = 1 − max(*R̄*ₘₙ, 0) was defined, so that strongly dependent metrics lie close together (dₘₙ → 0) and independent metrics far apart (dₘₙ → 1). Metrics were then agglomerated by average linkage (UPGMA) [138], in which the distance between two clusters A and B is the mean of dₘₙ over all metric pairs with m in A and n in B; the merge height drawn in the dendrogram (Fig S8 in S1 File) is this between-cluster distance. A short merge height therefore marks a strong, possibly nonlinear association, not measurement redundancy.

### Functional form

For every pair, the shape of the relation was examined in each scenario and in both regression directions, so that neither metric was cast as the predictor. Linearity was tested with Ramsey’s RESET [100], the F test of whether the second to fourth powers of the fitted values improve a straight-line fit [139], with Benjamini–Hochberg correction over the 330 tests (55 pairs × 3 scenarios × 2 directions). A reversal of trend within the observed range was tested with the U-shape test of Lind and Mehlum [98], which asks whether the slope of a quadratic fit changes sign between the smallest and the largest observed predictor value, corrected in the same way, and with the two-lines test of Simonsohn [99], which assumes no functional form, run with the author’s code and corrected in the same way. Four two-parameter candidate forms, all monotone, were then fitted by least squares to each of the six coupled pairs in both directions: the straight line y = b_0_ + b_1_x, the one-term exponential y = a·exp[b(x − x̃)], the power law y = a·x^b^ and the logarithm y = a·ln x + b, where y is the response metric, x the predictor metric, x̃ the median of the predictor pooled over the three scenarios, and a, b, b_0_ and b_1_ the fitted coefficients. In each contest, one combination of pair, scenario and direction, the candidates were ranked by the corrected Akaike Information Criterion (AICc) [101–103], computed from the residual sum of squares RSS of a fit to n cells with p estimated parameters, its two coefficients and the error variance (p = 3 for every candidate), assuming Gaussian errors, where ln is the natural logarithm:

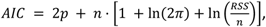

with the small-sample correction

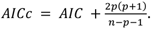

For each candidate i in a contest we computed the AICc difference Δ_i_ = AICc_i_ − min(AICc) and the Akaike weight

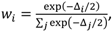

where j runs over the candidates of the contest [140]. The rankings of the 36 contests were aggregated by Borda count: in each contest c, candidate i, with rank r_ic_ from 1 for the lowest AICc to K = 4 for the highest, receives K − r_ic_ points, that is, 3, 2, 1 and 0 points from the best-to the worst-ranked candidate [104,141], and the Borda score of candidate i, B_i_ = Σ_c_ (K − r_ic_), sums these points over the 36 contests, with a maximum of 108. No two candidates tied in AICc in any contest. Because B_i_ = 36(K − r̄_i_), where r̄_i_ is the mean rank of candidate i over the 36 contests, the score rewards consistent placement across contests rather than first places alone. Because it ignores the size of the AICc gaps, which between the first two candidates of a contest ranged from 0.002 to 3.8 units, it was read together with the mean AICc difference and the mean Akaike weight of each candidate [141] (Table S14 and Fig S1 in S1 File).

## Acknowledgments

We acknowledge Dr. Fyodor Morozko for fruitful discussion and suggestions regarding the methodology of quantitative analyses.

## Author contributions

**JC-P:** conceptualization, data curation, formal analysis, investigation, methodology, software, validation, visualization, writing – original draft. **IM:** funding acquisition, methodology, writing – review & editing. **GP-Y:** resources, writing – review & editing. **BC-P:** investigation, writing – review & editing. **JMC:** funding acquisition, writing – review & editing. **JIL:** investigation, writing – review & editing. **IMDF:** project administration, supervision, writing – review & editing.

## Data and code availability

The cell trajectories and the MATLAB and R code that reproduces every result and table of the paper, together with the files it writes, are available on GitHub (https://github.com/JoseCarrascoPujante/proteus-migration-correlations/releases/tag/bioRxiv) and Zenodo (https://doi.org/10.5281/zenodo.23508490), the code under the MIT license and the data under CC BY 4.0. The per-cell values of all metrics are also given in Tables S1 to S11 in S1 File.

## Supporting Information S1 File. Supporting figures and tables

Figs S1–S10 and Tables S1–S15. (PDF)

