## Supplementary material for "Systemic and kinematic properties of *Amoeba proteus* migration correlate strongly across environments": S1 File

### Supporting Information S1 File. Supporting figures and tables

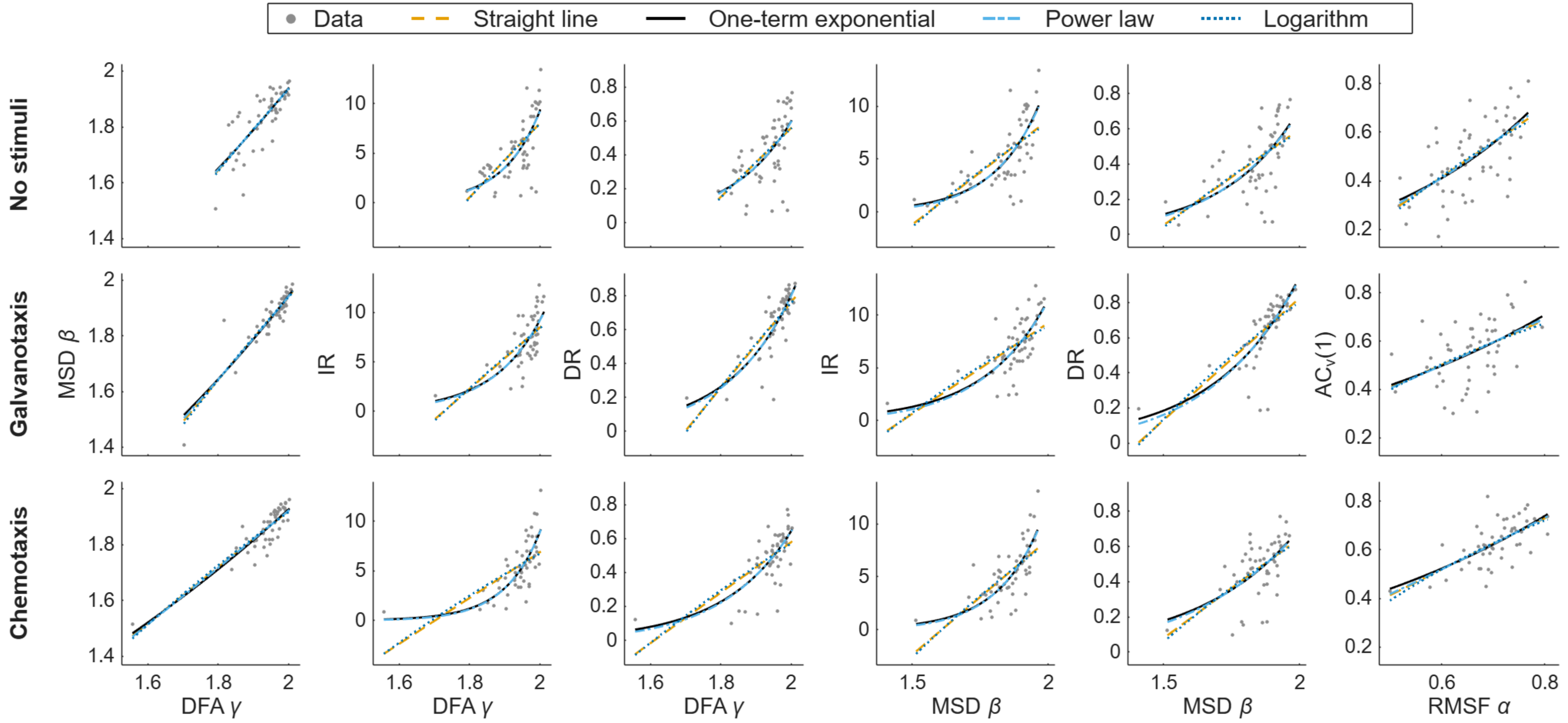


**Fig S1. Candidate functional forms for the six couplings.** One column per coupling, in the order of Table 1, and one row per scenario. The second metric of each coupling is plotted against the first, the regression of B on A in the notation of Table S13. The three panels (one panel per experimental scenario tested) of a coupling share their axes. Each panel shows the cells and the four two-parameter candidates fitted by least squares (straight line, one-term exponential, power law and logarithm). The one-term exponential and the power law nearly coincide over the observed range, as do the straight line and the logarithm; AICc separates none of the four decisively (Table S14).

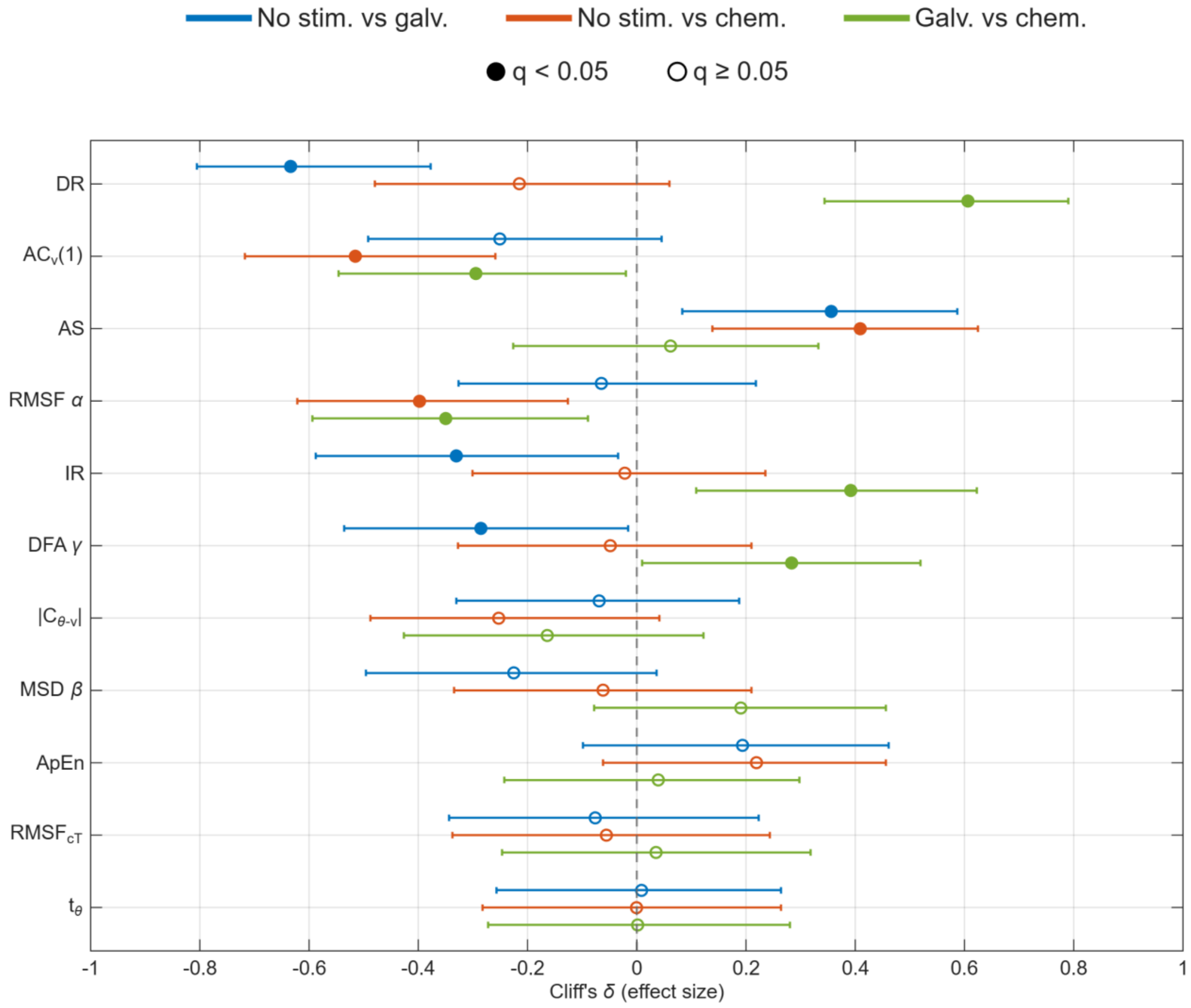


**Fig S2. Stimulus effect sizes: forest plot.** Cliff's δ with its BCa confidence interval for each metric and each pair of scenarios. Rows are ordered by the Kruskal–Wallis effect size $\varepsilon^2$, and filled markers denote contrasts with Mann–Whitney $q < 0.05$ (q, the p-value adjusted by the Benjamini–Hochberg procedure over the 33 pairwise tests). The confidence level, 98.2%, keeps the false coverage rate of the intervals of these contrasts at or below 0.05 (Materials and Methods). The values are those of Table S12.

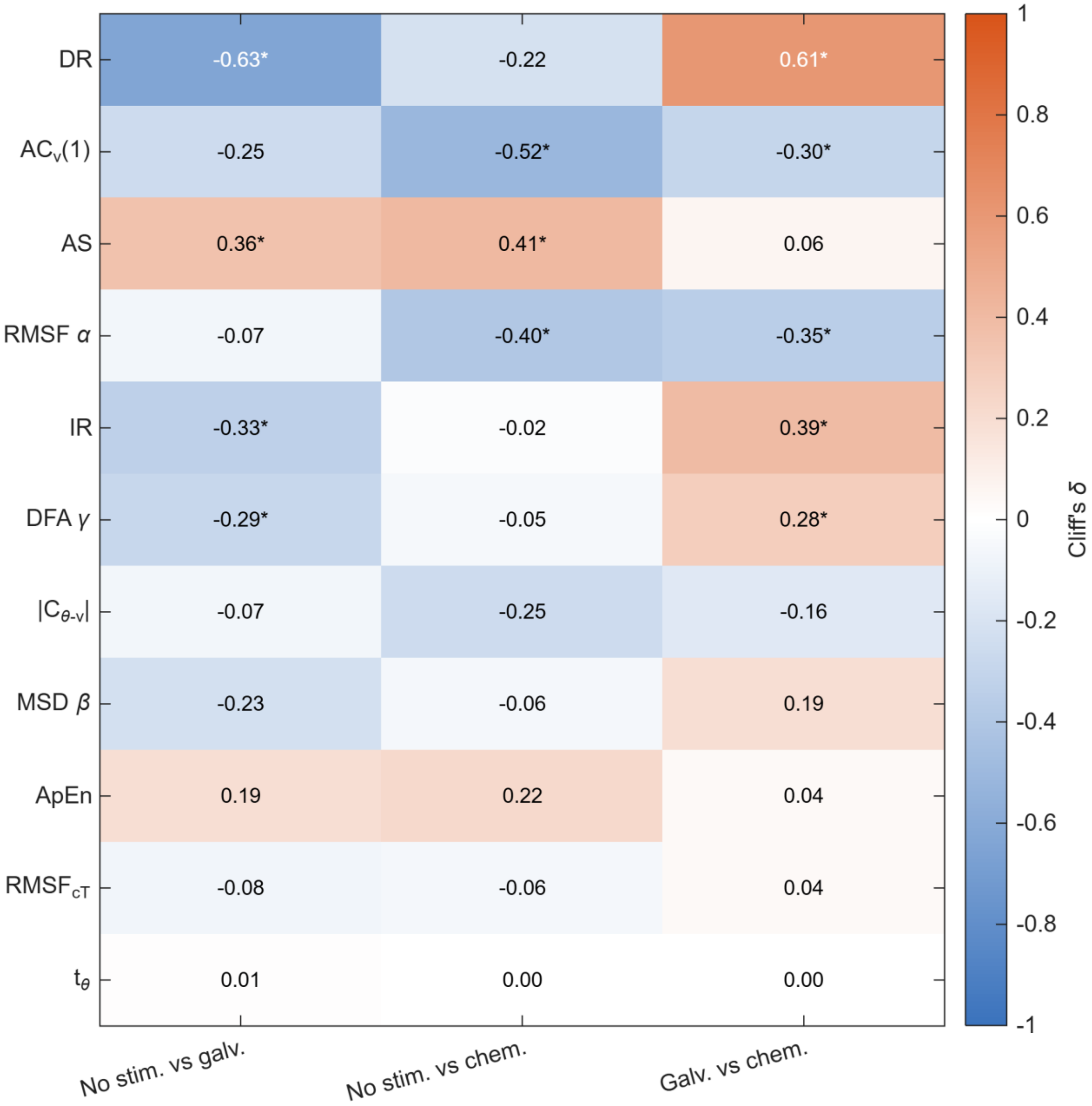


**Fig S3. Stimulus effect sizes: heatmap.** Cliff's δ for every metric and pair of scenarios on a diverging scale; an asterisk marks Mann–Whitney $q < 0.05$ ($q$, the p-value adjusted by the Benjamini–Hochberg procedure over the 33 pairwise tests), and rows are ordered by the Kruskal–Wallis effect size $\varepsilon^2$.

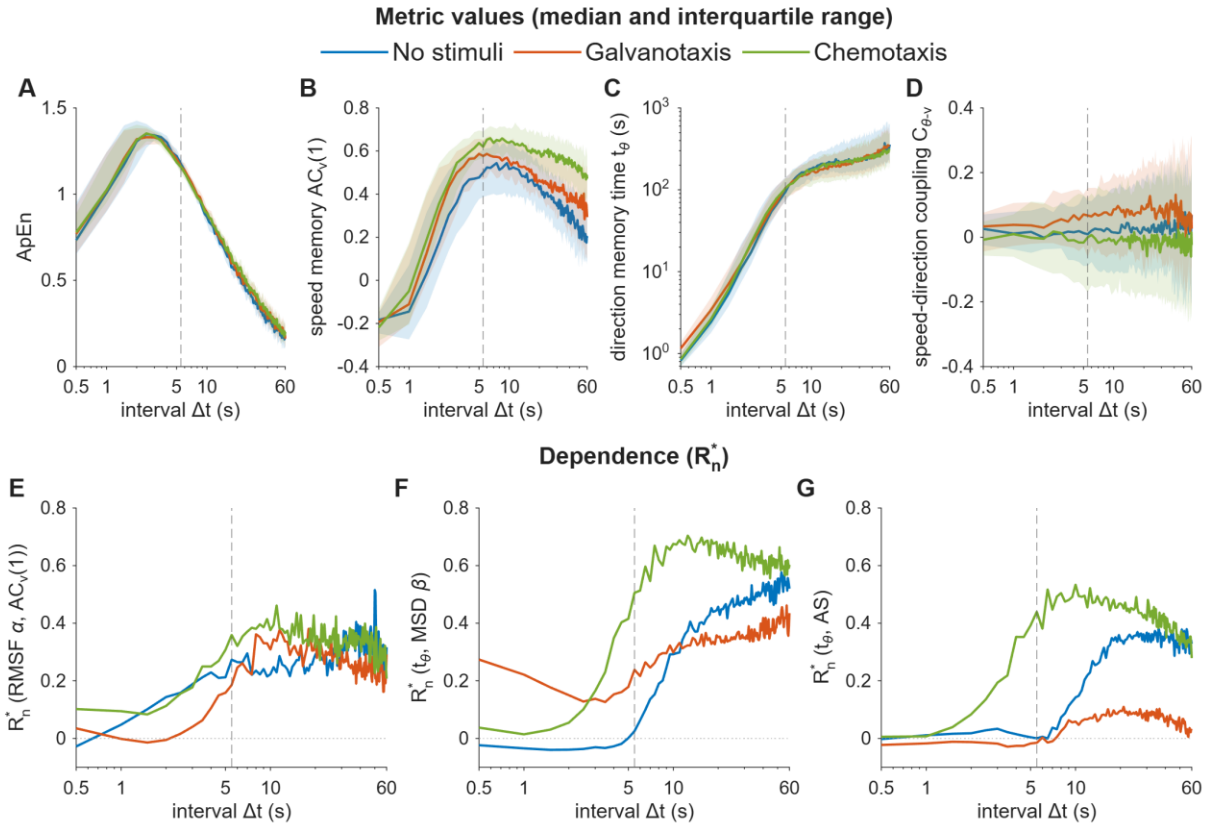


**Fig S4. Dependence on the sampling interval of the four metrics computed at the 5.5 s interval and of the dependences involving them on which the paper draws conclusions.** For every sampling interval Δt from 0.5 to 60 s, in steps of 0.5 s, and each scenario: the median and interquartile range over the 50 cells of **(A)** ApEn, **(B)** the speed memory $AC_v(1)$, **(C)** the direction-memory time $t_\theta$ (cells whose mean cosine of the turning angle is not positive at a given interval have no $t_\theta$ there and are left out) and **(D)** the speed-direction coupling $C_{\theta\text{-}v}$; and the bias-corrected squared distance correlation $R^*_n$ **(E)** of $AC_v(1)$ with RMSF α, one of the six couplings, and of $t_\theta$ with **(F)** MSD β and **(G)** AS, the two dependences compared between scenarios in the Results. In (E-G), RMSF α, MSD β and AS are computed from every frame and are not subsampled, so they keep their main-text values, computed at Δt = 0.5 s, and only ACv(1) and tθ vary with Δt. ApEn falls at long intervals mainly because the series shorten, from 327 steps at 5.5 s to 29 at 60 s. The dashed line marks the 5.5 s interval used in the paper, the shortest interval at which more than 99% of the steps were displacements; at shorter intervals zero-length steps become more frequent, reaching 56 to 58% at 0.5 s, because a manually tracked cell that moves less than one pixel between frames keeps its coordinates [72].

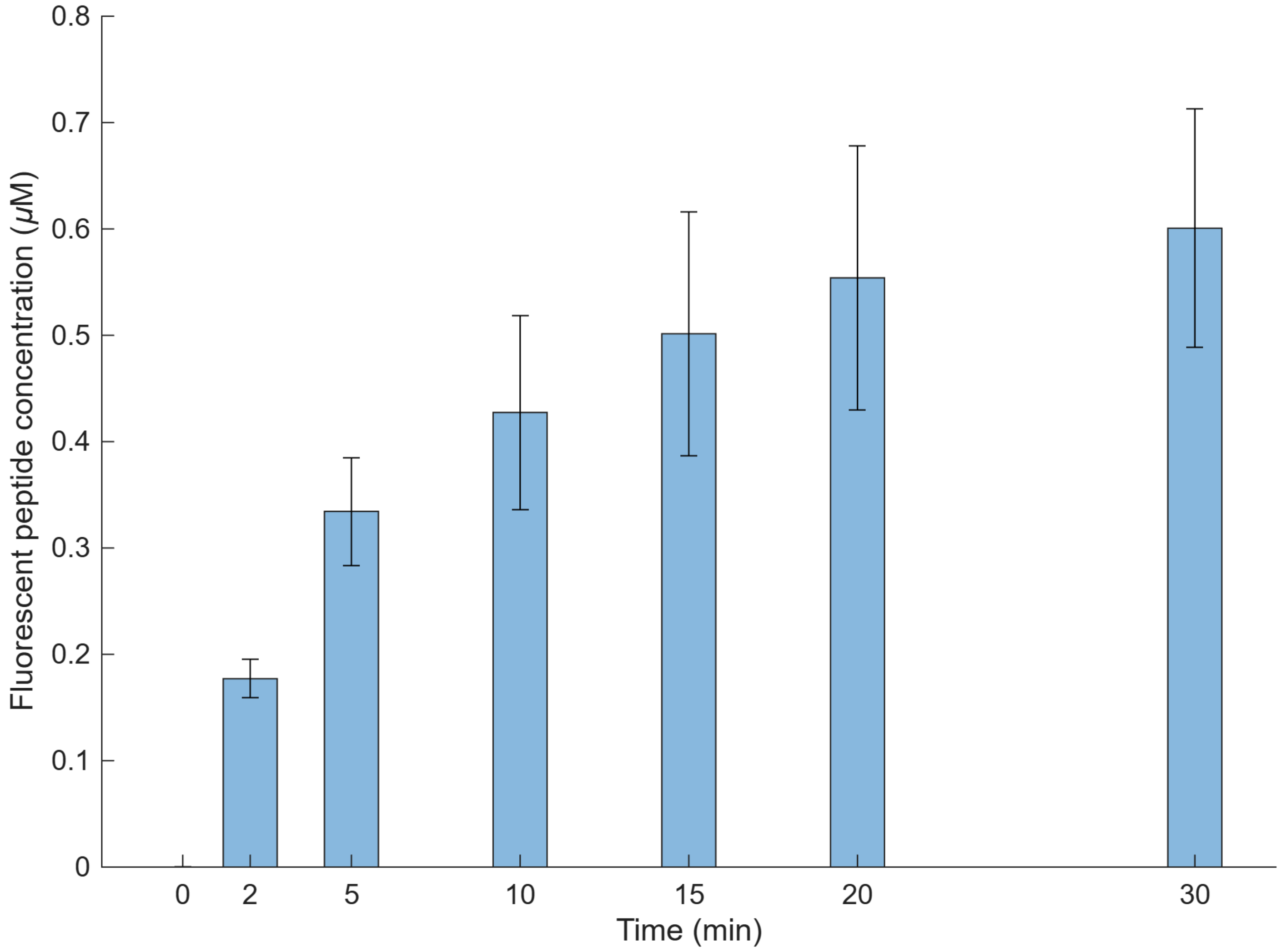


**Fig S5. Temporal dynamics of the fluorescein-tagged chemotactic peptide gradient.** Peptide concentration at the center of the experimental glass chamber, where the amoebae were placed at the start of each experiment, measured at 0, 2, 5, 10, 15, 20 and 30 min in two aliquots in each of three experiments (mean ± standard deviation of the six measurements). The concentration rises to about 0.2 μM within 2 min of establishing the laminar flow and to about 0.6 μM by 30 min, so the cells experience a sustained gradient throughout the assay.

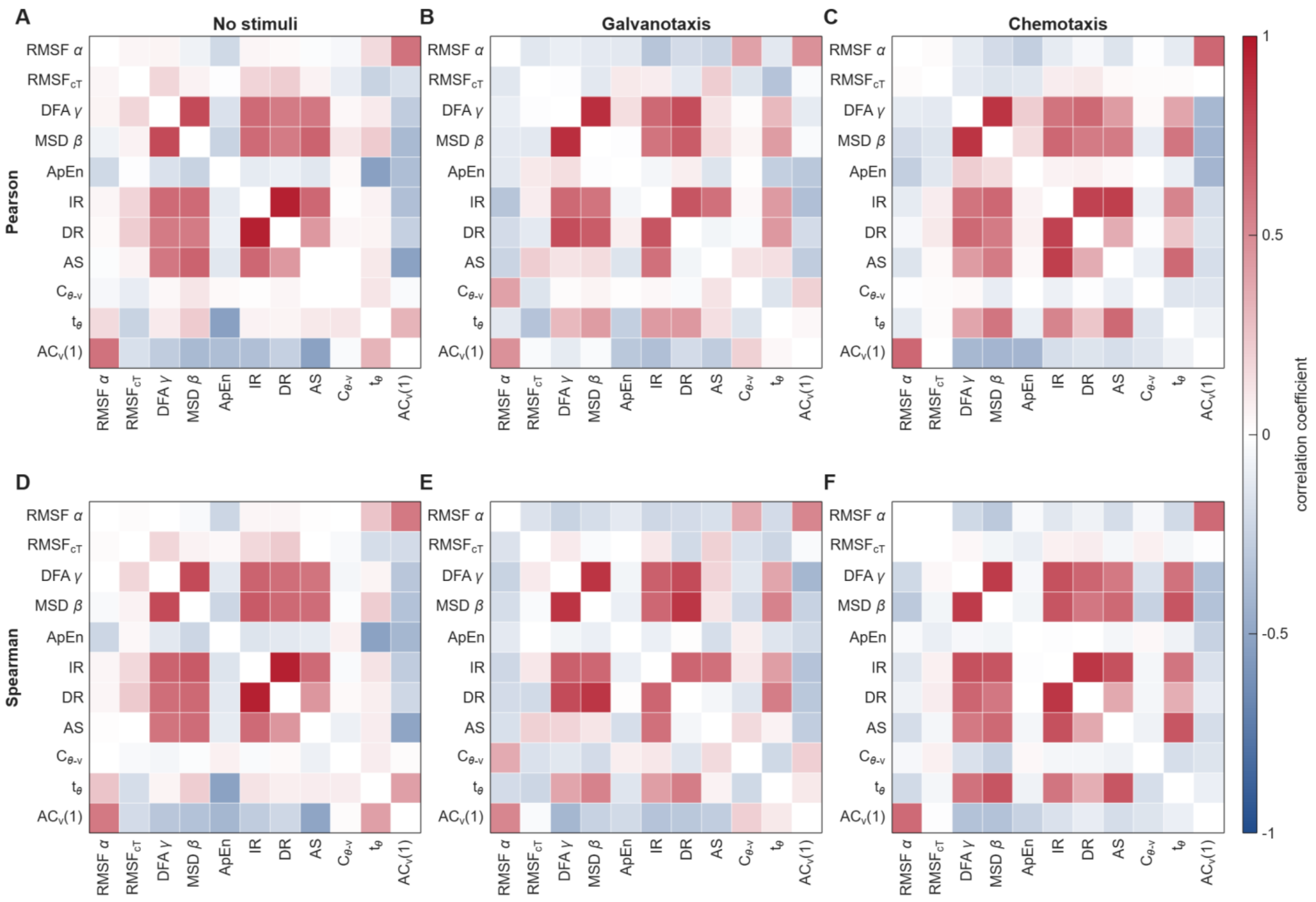


**Fig S6. Pearson and Spearman correlation matrices of the eleven metrics in each scenario.** Pearson (top row: **A**, no stimuli; **B**, galvanotaxis; **C**, chemotaxis) and Spearman (bottom row: **D to F**, same order) correlation coefficients computed on the 50 cells of each scenario, on a common diverging scale from −1 to 1; the unit diagonal is left blank. Pearson measures the linear component of a relation and Spearman any monotonic one, so a Spearman coefficient clearly larger in magnitude than its Pearson counterpart indicates a monotonic but curved relation.

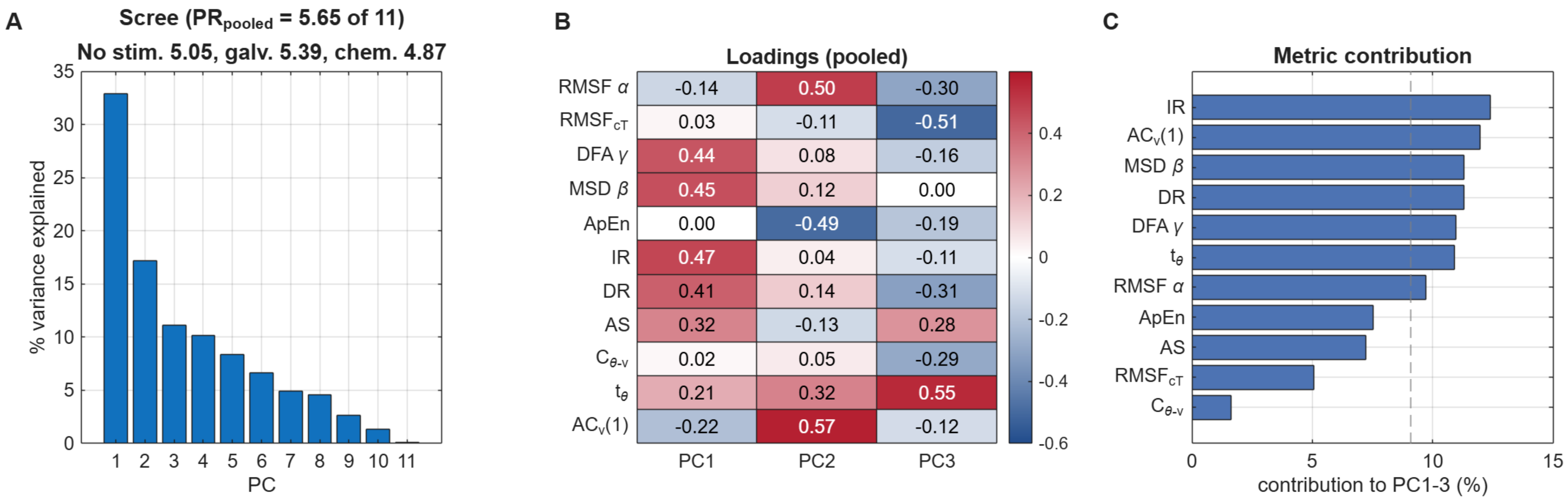


**Fig S7. Principal component analysis of the eleven metrics.** The metrics were z-scored over the 150 cells and decomposed on their correlation matrix. **(A)** variance explained by successive principal components, with the participation ratio PR, pooled and within each scenario. **(B)** loadings of the metrics on the first three components. **(C)** contribution of each metric to the first three components, the variance-weighted mean of its squared loadings; the dashed line marks the contribution of every metric if all contributed equally.

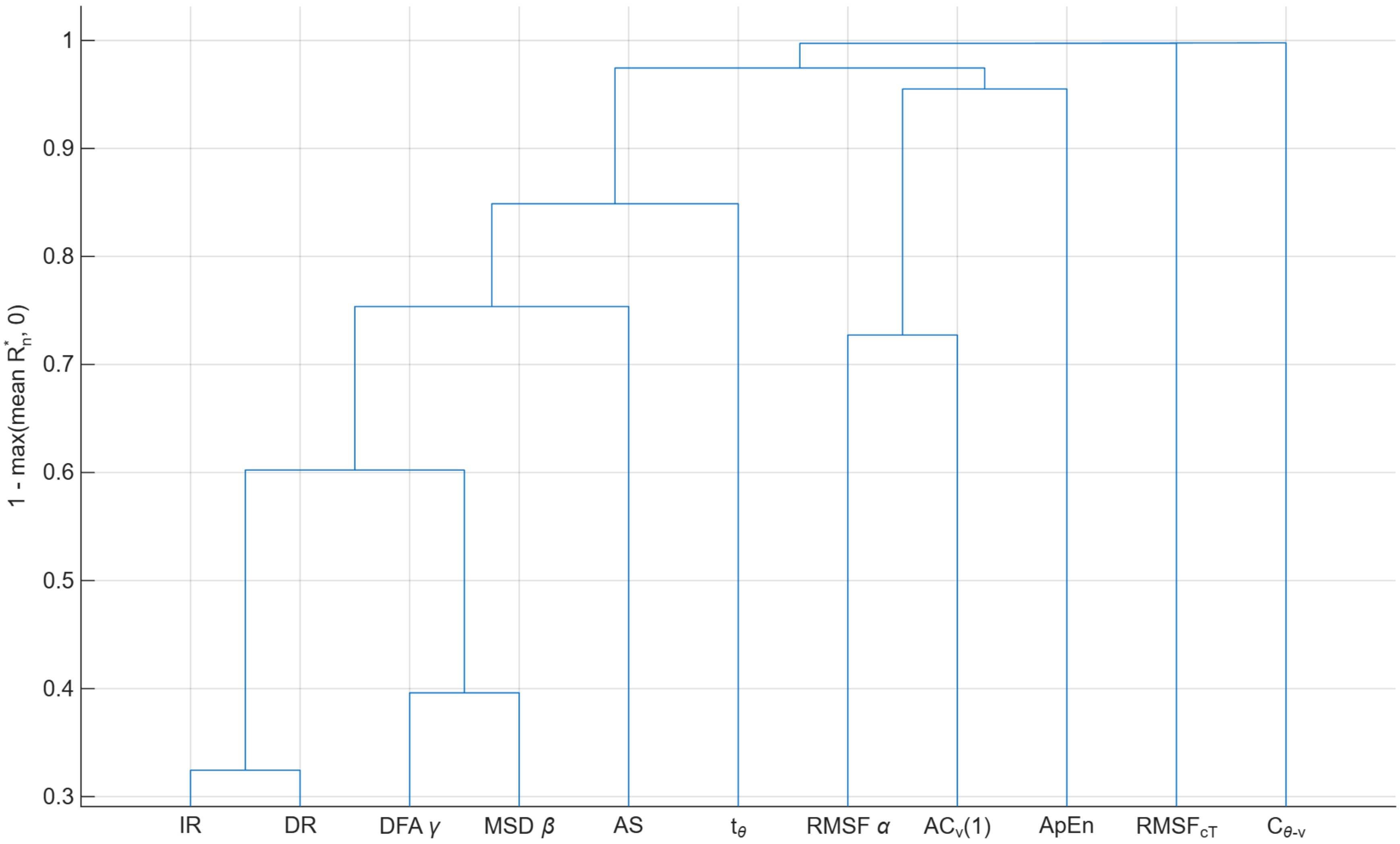


**Fig S8. Dendrogram of the metrics by statistical dependence.** Average linkage on 1 − max($R^*_n$, 0), with $R^*_n$ the bias-corrected squared distance correlation averaged over the three scenarios. A shorter merge height marks a stronger, possibly nonlinear, dependence, not redundancy between the metrics.

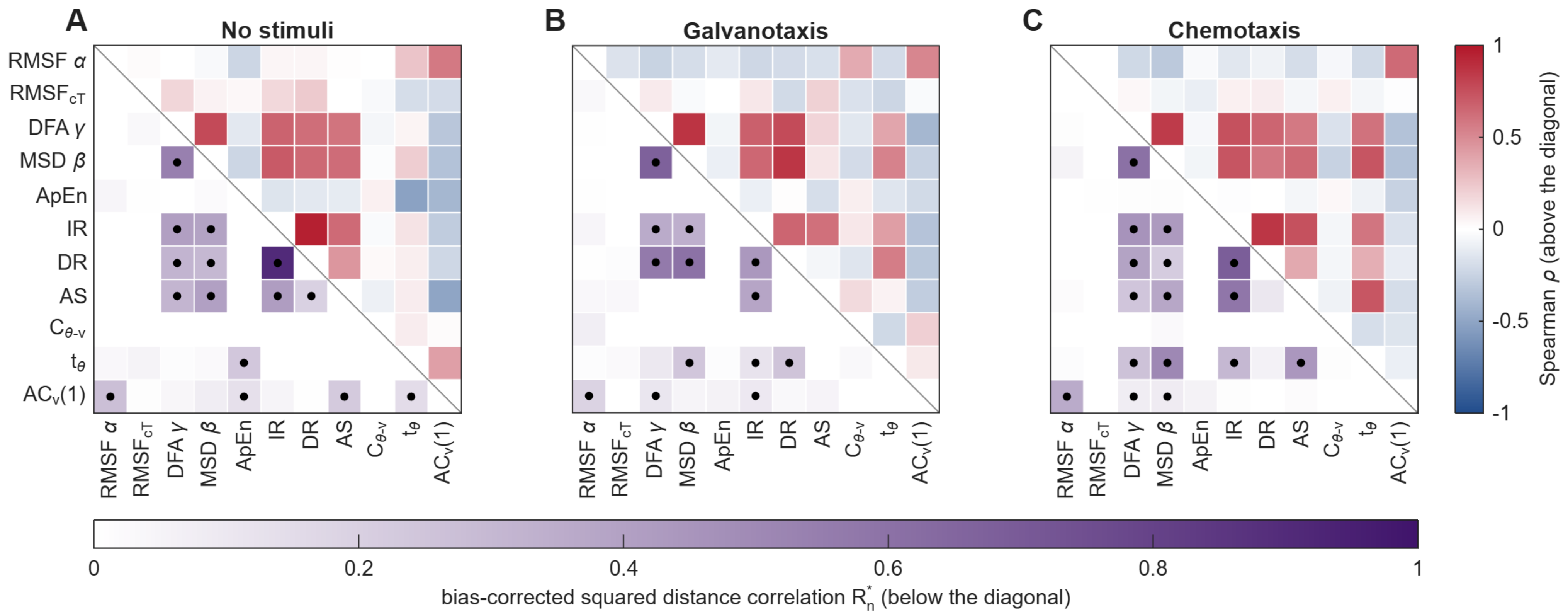


**Fig S9. Dependence matrices of the eleven metrics in each scenario.** **(A)** No stimuli, **(B)** galvanotaxis, **(C)** chemotaxis. Below the diagonal, the bias-corrected squared distance correlation $R^{*}_{n}$; above it, the Spearman correlation ρ, each with its own color scale. A dot marks a distance covariance test with Benjamini–Hochberg $q < 0.05$ in that scenario, and a pair is dependent when it carries the dot in all three panels. Negative $R^{*}_{n}$ values, possible under independence, are drawn as zero. The values are those of Table S13.

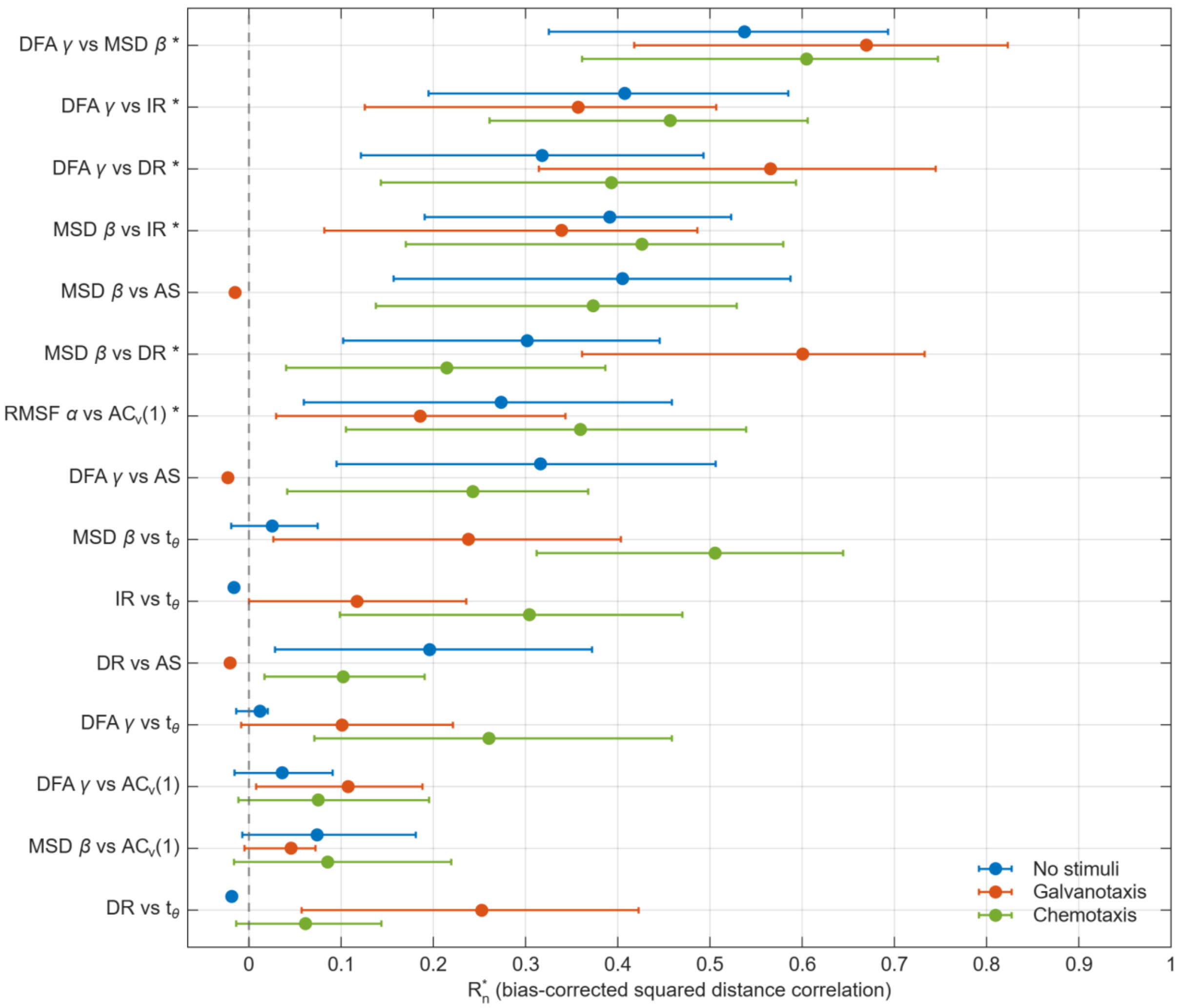


**Fig S10. Forest plot of the strongest pairwise dependences.** $R^*{}_n$ with its bias-corrected and accelerated (BCa) bootstrap 95% confidence interval in each scenario, for the 15 non-definitional pairs with the largest median $R^*{}_n$. An asterisk marks the pairs dependent in all three scenarios. Near independence the BCa interval degenerates (see Table S13) and is not drawn.

**Table S1. Approximate Entropy (ApEn) of the 150 trajectories.** Last row: p-value of the Lilliefors normality test in each scenario.

| Cell | No stimuli | Galvanotaxis | Chemotaxis |
|---|---|---|---|
| 1 | 1.190 | 1.139 | 1.033 |
| 2 | 1.159 | 1.147 | 1.190 |
| 3 | 1.152 | 1.167 | 1.163 |
| 4 | 1.181 | 1.152 | 1.114 |
| 5 | 1.173 | 1.203 | 1.139 |
| 6 | 1.135 | 1.160 | 1.286 |
| 7 | 1.157 | 1.212 | 1.096 |
| 8 | 1.128 | 1.207 | 1.171 |
| 9 | 1.226 | 1.117 | 1.256 |
| 10 | 1.165 | 1.281 | 1.135 |
| 11 | 1.198 | 1.254 | 1.300 |
| 12 | 1.119 | 1.135 | 1.161 |
| 13 | 1.134 | 1.066 | 1.101 |
| 14 | 1.128 | 1.172 | 1.162 |
| 15 | 1.226 | 1.134 | 1.152 |
| 16 | 1.244 | 1.163 | 1.092 |
| 17 | 1.176 | 1.225 | 1.201 |
| 18 | 1.222 | 1.155 | 1.175 |
| 19 | 1.300 | 1.254 | 1.088 |
| 20 | 1.084 | 1.298 | 1.157 |
| 21 | 1.248 | 1.191 | 1.139 |
| 22 | 1.209 | 1.189 | 1.199 |
| 23 | 1.186 | 1.136 | 1.185 |
| 24 | 1.224 | 1.175 | 1.145 |
| 25 | 1.122 | 1.157 | 1.116 |
| 26 | 1.175 | 1.125 | 1.188 |
| 27 | 1.115 | 1.116 | 1.164 |
| 28 | 1.188 | 1.152 | 1.194 |
| 29 | 1.132 | 1.167 | 1.120 |
| 30 | 1.171 | 1.071 | 1.179 |
| 31 | 1.149 | 1.169 | 1.151 |
| 32 | 1.284 | 1.140 | 1.114 |
| 33 | 1.302 | 1.124 | 1.155 |
| 34 | 1.178 | 1.176 | 1.206 |
| 35 | 1.129 | 1.110 | 1.255 |
| 36 | 1.098 | 1.165 | 1.148 |
| 37 | 1.155 | 1.118 | 1.088 |
| 38 | 1.191 | 1.156 | 1.156 |
| 39 | 1.170 | 1.148 | 1.135 |
| 40 | 1.199 | 1.121 | 1.089 |
| 41 | 1.211 | 1.132 | 1.187 |
| 42 | 1.033 | 1.196 | 1.159 |
| 43 | 1.209 | 1.203 | 1.178 |
| 44 | 1.159 | 1.070 | 1.082 |
| 45 | 1.138 | 1.162 | 1.170 |
| 46 | 1.259 | 1.146 | 1.221 |
| 47 | 1.192 | 1.188 | 1.185 |
| 48 | 1.084 | 1.180 | 1.184 |
| 49 | 1.230 | 1.120 | 1.081 |
| 50 | 1.252 | 1.167 | 1.138 |
| Lilliefors p | > 0.5 | 0.1700 | 0.3612 |

**Table S2. MSD diffusion exponent β of the 150 trajectories.** Last row: p-value of the Lilliefors normality test in each scenario.

| Cell | No stimuli | Galvanotaxis | Chemotaxis |
|---|---|---|---|
| 1 | 1.557 | 1.910 | 1.710 |
| 2 | 1.898 | 1.870 | 1.820 |
| 3 | 1.838 | 1.897 | 1.827 |
| 4 | 1.892 | 1.917 | 1.887 |
| 5 | 1.911 | 1.886 | 1.930 |
| 6 | 1.942 | 1.916 | 1.920 |
| 7 | 1.900 | 1.958 | 1.753 |
| 8 | 1.874 | 1.864 | 1.922 |
| 9 | 1.713 | 1.887 | 1.882 |
| 10 | 1.507 | 1.806 | 1.917 |
| 11 | 1.793 | 1.854 | 1.801 |
| 12 | 1.815 | 1.807 | 1.932 |
| 13 | 1.680 | 1.855 | 1.898 |
| 14 | 1.850 | 1.860 | 1.818 |
| 15 | 1.851 | 1.930 | 1.950 |
| 16 | 1.664 | 1.779 | 1.818 |
| 17 | 1.719 | 1.879 | 1.825 |
| 18 | 1.818 | 1.986 | 1.839 |
| 19 | 1.648 | 1.840 | 1.917 |
| 20 | 1.828 | 1.932 | 1.850 |
| 21 | 1.705 | 1.955 | 1.881 |
| 22 | 1.748 | 1.875 | 1.868 |
| 23 | 1.894 | 1.873 | 1.819 |
| 24 | 1.923 | 1.895 | 1.881 |
| 25 | 1.936 | 1.941 | 1.868 |
| 26 | 1.704 | 1.856 | 1.864 |
| 27 | 1.924 | 1.917 | 1.880 |
| 28 | 1.862 | 1.796 | 1.885 |
| 29 | 1.851 | 1.904 | 1.931 |
| 30 | 1.908 | 1.914 | 1.789 |
| 31 | 1.914 | 1.923 | 1.934 |
| 32 | 1.921 | 1.408 | 1.961 |
| 33 | 1.822 | 1.669 | 1.926 |
| 34 | 1.878 | 1.966 | 1.907 |
| 35 | 1.859 | 1.954 | 1.948 |
| 36 | 1.926 | 1.833 | 1.902 |
| 37 | 1.860 | 1.883 | 1.910 |
| 38 | 1.958 | 1.814 | 1.908 |
| 39 | 1.841 | 1.871 | 1.811 |
| 40 | 1.942 | 1.916 | 1.946 |
| 41 | 1.864 | 1.906 | 1.856 |
| 42 | 1.921 | 1.894 | 1.924 |
| 43 | 1.920 | 1.833 | 1.727 |
| 44 | 1.808 | 1.975 | 1.514 |
| 45 | 1.965 | 1.838 | 1.816 |
| 46 | 1.820 | 1.905 | 1.794 |
| 47 | 1.900 | 1.887 | 1.760 |
| 48 | 1.871 | 1.927 | 1.854 |
| 49 | 1.921 | 1.942 | 1.791 |
| 50 | 1.894 | 1.935 | 1.771 |
| Lilliefors p | 0.0014 | $< 0.001$ | 0.1144 |

**Table S3. DFA scaling exponent γ of the 150 trajectories.** Last row: p-value of the Lilliefors normality test in each scenario.

| Cell | No stimuli | Galvanotaxis | Chemotaxis |
|---|---|---|---|
| 1 | 1.870 | 1.995 | 1.839 |
| 2 | 1.930 | 1.955 | 1.880 |
| 3 | 1.856 | 1.981 | 1.970 |
| 4 | 1.920 | 1.977 | 1.997 |
| 5 | 1.910 | 1.947 | 1.989 |
| 6 | 1.948 | 1.973 | 1.963 |
| 7 | 1.989 | 1.994 | 1.830 |
| 8 | 1.953 | 1.977 | 1.992 |
| 9 | 1.899 | 1.989 | 1.947 |
| 10 | 1.792 | 1.943 | 1.959 |
| 11 | 1.910 | 1.984 | 1.948 |
| 12 | 1.921 | 1.939 | 1.971 |
| 13 | 1.853 | 1.818 | 1.963 |
| 14 | 1.860 | 1.948 | 1.934 |
| 15 | 1.965 | 1.991 | 1.991 |
| 16 | 1.873 | 1.908 | 1.955 |
| 17 | 1.927 | 1.966 | 1.908 |
| 18 | 1.840 | 2.012 | 1.928 |
| 19 | 1.842 | 1.938 | 1.969 |
| 20 | 1.964 | 1.989 | 1.955 |
| 21 | 1.858 | 1.995 | 1.941 |
| 22 | 1.953 | 1.966 | 1.946 |
| 23 | 1.973 | 1.951 | 1.946 |
| 24 | 1.981 | 1.937 | 1.930 |
| 25 | 1.956 | 1.993 | 1.945 |
| 26 | 1.835 | 1.928 | 1.870 |
| 27 | 1.995 | 1.985 | 1.960 |
| 28 | 1.977 | 1.886 | 1.984 |
| 29 | 1.967 | 1.986 | 1.987 |
| 30 | 1.990 | 1.992 | 1.852 |
| 31 | 2.000 | 1.953 | 1.985 |
| 32 | 1.987 | 1.702 | 2.003 |
| 33 | 1.955 | 1.849 | 1.975 |
| 34 | 1.953 | 2.007 | 1.960 |
| 35 | 1.952 | 1.997 | 1.992 |
| 36 | 1.989 | 1.911 | 1.980 |
| 37 | 1.962 | 1.951 | 1.950 |
| 38 | 1.998 | 1.916 | 1.968 |
| 39 | 1.914 | 1.974 | 1.901 |
| 40 | 1.977 | 1.979 | 1.978 |
| 41 | 1.963 | 1.987 | 1.973 |
| 42 | 1.965 | 1.974 | 1.961 |
| 43 | 1.979 | 1.943 | 1.904 |
| 44 | 1.830 | 1.995 | 1.556 |
| 45 | 2.001 | 1.943 | 1.909 |
| 46 | 1.888 | 1.985 | 1.904 |
| 47 | 1.952 | 1.971 | 1.938 |
| 48 | 1.946 | 1.990 | 1.928 |
| 49 | 1.982 | 1.988 | 1.930 |
| 50 | 1.952 | 1.980 | 1.885 |
| Lilliefors p | < 0.001 | < 0.001 | < 0.001 |

**Table S4. RMSF exponent α of the 150 trajectories.** Last row: p-value of the Lilliefors normality test in each scenario.

| Cell | No stimuli | Galvanotaxis | Chemotaxis |
|---|---|---|---|
| 1 | 0.704 | 0.611 | 0.805 |
| 2 | 0.646 | 0.659 | 0.684 |
| 3 | 0.518 | 0.595 | 0.806 |
| 4 | 0.720 | 0.502 | 0.691 |
| 5 | 0.579 | 0.598 | 0.693 |
| 6 | 0.676 | 0.688 | 0.685 |
| 7 | 0.537 | 0.622 | 0.752 |
| 8 | 0.727 | 0.668 | 0.677 |
| 9 | 0.734 | 0.701 | 0.639 |
| 10 | 0.768 | 0.640 | 0.577 |
| 11 | 0.573 | 0.605 | 0.499 |
| 12 | 0.655 | 0.612 | 0.676 |
| 13 | 0.655 | 0.652 | 0.653 |
| 14 | 0.725 | 0.673 | 0.781 |
| 15 | 0.589 | 0.650 | 0.675 |
| 16 | 0.653 | 0.707 | 0.670 |
| 17 | 0.654 | 0.746 | 0.744 |
| 18 | 0.640 | 0.509 | 0.741 |
| 19 | 0.592 | 0.656 | 0.642 |
| 20 | 0.645 | 0.653 | 0.698 |
| 21 | 0.531 | 0.647 | 0.755 |
| 22 | 0.732 | 0.690 | 0.716 |
| 23 | 0.709 | 0.727 | 0.722 |
| 24 | 0.601 | 0.691 | 0.680 |
| 25 | 0.637 | 0.690 | 0.766 |
| 26 | 0.590 | 0.735 | 0.709 |
| 27 | 0.707 | 0.691 | 0.601 |
| 28 | 0.681 | 0.561 | 0.746 |
| 29 | 0.690 | 0.656 | 0.737 |
| 30 | 0.641 | 0.786 | 0.685 |
| 31 | 0.517 | 0.698 | 0.682 |
| 32 | 0.603 | 0.702 | 0.634 |
| 33 | 0.620 | 0.590 | 0.673 |
| 34 | 0.754 | 0.602 | 0.698 |
| 35 | 0.735 | 0.579 | 0.716 |
| 36 | 0.713 | 0.675 | 0.737 |
| 37 | 0.742 | 0.738 | 0.735 |
| 38 | 0.733 | 0.763 | 0.699 |
| 39 | 0.685 | 0.795 | 0.669 |
| 40 | 0.611 | 0.740 | 0.750 |
| 41 | 0.634 | 0.571 | 0.767 |
| 42 | 0.611 | 0.788 | 0.727 |
| 43 | 0.681 | 0.704 | 0.720 |
| 44 | 0.659 | 0.711 | 0.689 |
| 45 | 0.593 | 0.703 | 0.697 |
| 46 | 0.603 | 0.661 | 0.772 |
| 47 | 0.696 | 0.627 | 0.718 |
| 48 | 0.749 | 0.617 | 0.608 |
| 49 | 0.728 | 0.651 | 0.740 |
| 50 | 0.646 | 0.691 | 0.762 |
| Lilliefors p | 0.4196 | > 0.5 | 0.0609 |

**Table S5. RMSF correlation time (minutes) of the 150 trajectories.** Last row: p-value of the Lilliefors normality test in each scenario.

| Cell | No stimuli | Galvanotaxis | Chemotaxis |
|---|---|---|---|
| 1 | 3.125 | 16.667 | 11.458 |
| 2 | 4.167 | 15.625 | 7.292 |
| 3 | 9.375 | 14.583 | 15.625 |
| 4 | 3.125 | 12.500 | 15.625 |
| 5 | 4.167 | 13.542 | 4.167 |
| 6 | 13.542 | 15.625 | 10.417 |
| 7 | 1.042 | 1.042 | 9.375 |
| 8 | 9.375 | 15.625 | 7.292 |
| 9 | 3.125 | 15.625 | 11.458 |
| 10 | 6.250 | 8.333 | 13.542 |
| 11 | 11.458 | 14.583 | 8.333 |
| 12 | 2.083 | 11.458 | 14.583 |
| 13 | 12.500 | 4.167 | 12.500 |
| 14 | 3.125 | 9.375 | 11.458 |
| 15 | 13.542 | 10.417 | 7.292 |
| 16 | 2.083 | 10.417 | 13.542 |
| 17 | 15.625 | 10.417 | 12.500 |
| 18 | 2.083 | 12.500 | 3.125 |
| 19 | 13.542 | 7.292 | 9.375 |
| 20 | 9.375 | 16.667 | 7.292 |
| 21 | 14.583 | 13.542 | 11.458 |
| 22 | 14.583 | 2.083 | 13.542 |
| 23 | 11.458 | 11.458 | 11.458 |
| 24 | 2.083 | 2.083 | 3.125 |
| 25 | 15.625 | 1.042 | 8.333 |
| 26 | 12.500 | 7.292 | 4.167 |
| 27 | 3.125 | 6.250 | 6.250 |
| 28 | 12.500 | 14.583 | 15.625 |
| 29 | 12.500 | 2.083 | 11.458 |
| 30 | 15.625 | 10.417 | 10.417 |
| 31 | 6.250 | 10.417 | 10.417 |
| 32 | 3.125 | 15.625 | 10.417 |
| 33 | 14.583 | 8.333 | 12.500 |
| 34 | 14.583 | 10.417 | 12.500 |
| 35 | 10.417 | 15.625 | 10.417 |
| 36 | 11.458 | 12.500 | 3.125 |
| 37 | 10.417 | 11.458 | 9.375 |
| 38 | 9.375 | 6.250 | 7.292 |
| 39 | 5.208 | 14.583 | 7.292 |
| 40 | 4.167 | 12.500 | 9.375 |
| 41 | 14.583 | 11.458 | 9.375 |
| 42 | 8.333 | 14.583 | 8.333 |
| 43 | 15.625 | 3.125 | 11.458 |
| 44 | 12.500 | 1.042 | 15.625 |
| 45 | 16.667 | 2.083 | 10.417 |
| 46 | 2.083 | 6.250 | 3.125 |
| 47 | 12.500 | 6.250 | 4.167 |
| 48 | 9.375 | 7.292 | 13.542 |
| 49 | 14.583 | 3.125 | 10.417 |
| 50 | 6.250 | 14.583 | 15.625 |
| Lilliefors p | 0.0072 | 0.0124 | 0.1076 |

**Table S6. Speed memory $AC_v(1)$ of the 150 trajectories.** Last row: p-value of the Lilliefors normality test in each scenario.

| Cell | No stimuli | Galvanotaxis | Chemotaxis |
|---|---|---|---|
| 1 | 0.699 | 0.584 | 0.739 |
| 2 | 0.466 | 0.650 | 0.603 |
| 3 | 0.411 | 0.468 | 0.664 |
| 4 | 0.607 | 0.546 | 0.520 |
| 5 | 0.424 | 0.629 | 0.586 |
| 6 | 0.537 | 0.645 | 0.448 |
| 7 | 0.292 | 0.303 | 0.623 |
| 8 | 0.701 | 0.306 | 0.669 |
| 9 | 0.782 | 0.496 | 0.520 |
| 10 | 0.810 | 0.354 | 0.447 |
| 11 | 0.542 | 0.325 | 0.430 |
| 12 | 0.590 | 0.565 | 0.521 |
| 13 | 0.732 | 0.493 | 0.547 |
| 14 | 0.668 | 0.595 | 0.727 |
| 15 | 0.616 | 0.404 | 0.603 |
| 16 | 0.577 | 0.556 | 0.613 |
| 17 | 0.408 | 0.496 | 0.676 |
| 18 | 0.505 | 0.391 | 0.690 |
| 19 | 0.387 | 0.652 | 0.599 |
| 20 | 0.607 | 0.437 | 0.652 |
| 21 | 0.221 | 0.380 | 0.674 |
| 22 | 0.495 | 0.499 | 0.656 |
| 23 | 0.423 | 0.791 | 0.690 |
| 24 | 0.437 | 0.607 | 0.583 |
| 25 | 0.559 | 0.389 | 0.717 |
| 26 | 0.558 | 0.753 | 0.744 |
| 27 | 0.646 | 0.551 | 0.619 |
| 28 | 0.456 | 0.682 | 0.622 |
| 29 | 0.565 | 0.581 | 0.688 |
| 30 | 0.343 | 0.684 | 0.617 |
| 31 | 0.296 | 0.641 | 0.622 |
| 32 | 0.356 | 0.464 | 0.552 |
| 33 | 0.353 | 0.413 | 0.523 |
| 34 | 0.579 | 0.434 | 0.609 |
| 35 | 0.588 | 0.531 | 0.568 |
| 36 | 0.555 | 0.712 | 0.527 |
| 37 | 0.658 | 0.741 | 0.677 |
| 38 | 0.406 | 0.844 | 0.606 |
| 39 | 0.665 | 0.656 | 0.695 |
| 40 | 0.242 | 0.642 | 0.749 |
| 41 | 0.535 | 0.493 | 0.786 |
| 42 | 0.449 | 0.684 | 0.728 |
| 43 | 0.439 | 0.644 | 0.725 |
| 44 | 0.316 | 0.582 | 0.821 |
| 45 | 0.169 | 0.690 | 0.506 |
| 46 | 0.295 | 0.617 | 0.589 |
| 47 | 0.454 | 0.681 | 0.603 |
| 48 | 0.746 | 0.561 | 0.476 |
| 49 | 0.336 | 0.573 | 0.736 |
| 50 | 0.311 | 0.693 | 0.770 |
| Lilliefors p | > 0.5 | 0.3952 | > 0.5 |

**Table S7. Direction-memory time $t_\theta$ (s) of the 150 trajectories.** Last row: p-value of the Lilliefors normality test in each scenario.

| Cell | No stimuli | Galvanotaxis | Chemotaxis |
|---|---|---|---|
| 1 | 95.067 | 123.238 | 54.064 |
| 2 | 264.611 | 111.802 | 40.546 |
| 3 | 68.580 | 143.245 | 80.847 |
| 4 | 155.108 | 122.735 | 214.098 |
| 5 | 259.917 | 137.191 | 110.071 |
| 6 | 131.823 | 77.958 | 103.202 |
| 7 | 71.544 | 235.941 | 95.377 |
| 8 | 159.233 | 38.818 | 143.275 |
| 9 | 115.995 | 64.521 | 107.075 |
| 10 | 90.690 | 62.337 | 88.677 |
| 11 | 132.340 | 55.477 | 57.898 |
| 12 | 149.633 | 65.292 | 117.458 |
| 13 | 172.481 | 107.816 | 162.012 |
| 14 | 155.562 | 48.795 | 96.457 |
| 15 | 87.704 | 97.313 | 176.063 |
| 16 | 49.486 | 59.608 | 70.197 |
| 17 | 72.785 | 92.101 | 101.986 |
| 18 | 84.803 | 169.454 | 73.506 |
| 19 | 44.634 | 53.982 | 118.119 |
| 20 | 249.573 | 55.557 | 116.639 |
| 21 | 75.575 | 81.036 | 105.681 |
| 22 | 86.822 | 103.612 | 91.364 |
| 23 | 106.673 | 80.431 | 82.869 |
| 24 | 89.990 | 87.789 | 155.070 |
| 25 | 129.478 | 118.593 | 126.713 |
| 26 | 64.105 | 140.042 | 177.050 |
| 27 | 191.752 | 116.918 | 119.942 |
| 28 | 130.563 | 80.873 | 138.309 |
| 29 | 124.473 | 94.319 | 99.951 |
| 30 | 92.910 | 37.998 | 90.933 |
| 31 | 61.451 | 97.272 | 138.025 |
| 32 | 71.726 | 48.404 | 173.081 |
| 33 | 53.017 | 65.009 | 133.023 |
| 34 | 69.820 | 169.597 | 148.615 |
| 35 | 59.121 | 175.083 | 142.331 |
| 36 | 78.418 | 201.134 | 86.293 |
| 37 | 199.327 | 73.793 | 171.639 |
| 38 | 92.843 | 74.253 | 104.648 |
| 39 | 207.164 | 114.264 | 77.120 |
| 40 | 137.219 | 114.567 | 140.897 |
| 41 | 42.369 | 106.426 | 105.908 |
| 42 | 198.059 | 136.995 | 129.991 |
| 43 | 107.109 | 84.809 | 96.957 |
| 44 | 90.610 | 384.791 | 75.282 |
| 45 | 106.167 | 146.736 | 33.386 |
| 46 | 50.619 | 178.479 | 41.336 |
| 47 | 66.858 | 139.249 | 66.506 |
| 48 | 143.003 | 165.156 | 80.748 |
| 49 | 89.037 | 171.135 | 39.717 |
| 50 | 103.553 | 145.282 | 54.320 |
| Lilliefors p | 0.0038 | 0.1085 | > 0.5 |

**Table S8. Speed-direction coupling $C_{\theta\text{-}v}$ of the 150 trajectories.** Last row: p-value of the Lilliefors normality test in each scenario.

| Cell | No stimuli | Galvanotaxis | Chemotaxis |
|---|---|---|---|
| 1 | 0.014 | 0.085 | 0.142 |
| 2 | 0.197 | 0.086 | 0.261 |
| 3 | 0.088 | -0.127 | -0.038 |
| 4 | 0.013 | -0.021 | 0.190 |
| 5 | 0.128 | 0.065 | -0.107 |
| 6 | 0.006 | -0.104 | -0.256 |
| 7 | 0.103 | 0.055 | -0.172 |
| 8 | -0.036 | -0.075 | -0.361 |
| 9 | 0.113 | 0.175 | 0.326 |
| 10 | -0.272 | -0.016 | 0.259 |
| 11 | -0.128 | 0.316 | -0.036 |
| 12 | 0.187 | 0.162 | 0.061 |
| 13 | -0.088 | 0.211 | 0.110 |
| 14 | 0.198 | 0.231 | 0.339 |
| 15 | 0.211 | 0.084 | 0.043 |
| 16 | 0.073 | 0.109 | 0.120 |
| 17 | 0.138 | 0.146 | -0.213 |
| 18 | -0.015 | -0.225 | 0.119 |
| 19 | -0.226 | -0.015 | -0.093 |
| 20 | -0.014 | 0.153 | 0.338 |
| 21 | 0.057 | -0.010 | 0.238 |
| 22 | 0.086 | 0.076 | -0.143 |
| 23 | 0.131 | -0.196 | -0.241 |
| 24 | -0.088 | -0.081 | 0.009 |
| 25 | -0.268 | 0.145 | -0.048 |
| 26 | 0.123 | 0.374 | -0.006 |
| 27 | -0.008 | 0.058 | -0.262 |
| 28 | -0.217 | 0.122 | 0.099 |
| 29 | -0.051 | 0.241 | -0.064 |
| 30 | 0.054 | 0.476 | 0.251 |
| 31 | -0.089 | 0.341 | -0.100 |
| 32 | 0.024 | -0.148 | -0.091 |
| 33 | -0.038 | -0.064 | 0.311 |
| 34 | -0.295 | -0.094 | 0.033 |
| 35 | -0.154 | -0.149 | -0.286 |
| 36 | -0.007 | 0.013 | -0.042 |
| 37 | -0.239 | -0.059 | -0.333 |
| 38 | -0.012 | 0.193 | 0.005 |
| 39 | 0.226 | -0.061 | 0.107 |
| 40 | -0.008 | 0.084 | -0.333 |
| 41 | 0.093 | -0.084 | 0.195 |
| 42 | -0.143 | 0.333 | -0.332 |
| 43 | 0.249 | 0.308 | 0.015 |
| 44 | 0.019 | 0.138 | -0.213 |
| 45 | -0.043 | 0.084 | -0.077 |
| 46 | -0.134 | -0.019 | 0.060 |
| 47 | 0.094 | -0.001 | 0.200 |
| 48 | 0.187 | 0.013 | -0.170 |
| 49 | 0.236 | -0.181 | 0.090 |
| 50 | -0.095 | 0.070 | -0.182 |
| Lilliefors p | > 0.5 | 0.4804 | > 0.5 |

**Table S9. Intensity of the Response (IR, mm) of the 150 trajectories.** Last row: p-value of the Lilliefors normality test in each scenario.

| Cell | No stimuli | Galvanotaxis | Chemotaxis |
|---|---|---|---|
| 1 | 0.608 | 8.438 | 3.047 |
| 2 | 6.139 | 7.834 | 4.012 |
| 3 | 3.745 | 9.259 | 3.689 |
| 4 | 6.128 | 9.606 | 5.694 |
| 5 | 3.828 | 7.686 | 7.366 |
| 6 | 6.320 | 7.491 | 5.266 |
| 7 | 1.054 | 9.981 | 1.057 |
| 8 | 6.457 | 5.322 | 6.902 |
| 9 | 2.657 | 10.640 | 4.810 |
| 10 | 1.172 | 5.730 | 5.578 |
| 11 | 2.532 | 6.985 | 5.260 |
| 12 | 5.420 | 9.674 | 5.890 |
| 13 | 3.831 | 2.473 | 7.018 |
| 14 | 3.565 | 4.928 | 6.924 |
| 15 | 6.264 | 11.513 | 8.440 |
| 16 | 0.991 | 5.992 | 5.589 |
| 17 | 5.677 | 8.776 | 5.481 |
| 18 | 3.458 | 11.591 | 4.271 |
| 19 | 2.713 | 4.523 | 3.390 |
| 20 | 4.823 | 7.707 | 3.959 |
| 21 | 3.399 | 9.129 | 5.439 |
| 22 | 3.722 | 5.882 | 3.350 |
| 23 | 8.702 | 3.350 | 4.962 |
| 24 | 5.514 | 5.661 | 4.832 |
| 25 | 5.129 | 7.832 | 4.376 |
| 26 | 2.949 | 7.913 | 1.448 |
| 27 | 9.955 | 8.135 | 1.833 |
| 28 | 1.447 | 4.547 | 6.502 |
| 29 | 3.775 | 9.656 | 5.392 |
| 30 | 9.440 | 6.890 | 1.784 |
| 31 | 10.139 | 9.700 | 10.808 |
| 32 | 10.198 | 1.608 | 13.079 |
| 33 | 4.252 | 4.732 | 9.371 |
| 34 | 3.525 | 11.052 | 9.105 |
| 35 | 7.388 | 12.779 | 10.720 |
| 36 | 10.055 | 5.821 | 6.279 |
| 37 | 11.523 | 2.544 | 6.491 |
| 38 | 11.337 | 2.274 | 8.566 |
| 39 | 3.128 | 6.530 | 4.282 |
| 40 | 8.675 | 7.543 | 8.435 |
| 41 | 3.526 | 5.680 | 6.951 |
| 42 | 6.377 | 6.595 | 7.956 |
| 43 | 9.764 | 5.993 | 6.115 |
| 44 | 3.114 | 11.179 | 0.894 |
| 45 | 13.411 | 6.179 | 2.577 |
| 46 | 5.433 | 6.754 | 1.202 |
| 47 | 8.207 | 5.641 | 3.341 |
| 48 | 0.736 | 5.985 | 4.038 |
| 49 | 8.677 | 6.813 | 3.556 |
| 50 | 4.866 | 6.286 | 3.268 |
| Lilliefors p | 0.0136 | > 0.5 | > 0.5 |

**Table S10. Directionality Ratio (DR) of the 150 trajectories.** Last row: p-value of the Lilliefors normality test in each scenario.

| Cell | No stimuli | Galvanotaxis | Chemotaxis |
|---|---|---|---|
| 1 | 0.052 | 0.696 | 0.356 |
| 2 | 0.507 | 0.628 | 0.474 |
| 3 | 0.307 | 0.758 | 0.477 |
| 4 | 0.504 | 0.782 | 0.449 |
| 5 | 0.270 | 0.680 | 0.771 |
| 6 | 0.556 | 0.659 | 0.436 |
| 7 | 0.072 | 0.844 | 0.099 |
| 8 | 0.470 | 0.612 | 0.656 |
| 9 | 0.288 | 0.722 | 0.461 |
| 10 | 0.184 | 0.443 | 0.491 |
| 11 | 0.240 | 0.539 | 0.558 |
| 12 | 0.501 | 0.563 | 0.586 |
| 13 | 0.386 | 0.187 | 0.616 |
| 14 | 0.337 | 0.573 | 0.670 |
| 15 | 0.609 | 0.780 | 0.739 |
| 16 | 0.102 | 0.424 | 0.623 |
| 17 | 0.469 | 0.724 | 0.587 |
| 18 | 0.302 | 0.875 | 0.520 |
| 19 | 0.302 | 0.493 | 0.327 |
| 20 | 0.361 | 0.833 | 0.336 |
| 21 | 0.424 | 0.798 | 0.450 |
| 22 | 0.328 | 0.721 | 0.384 |
| 23 | 0.567 | 0.598 | 0.525 |
| 24 | 0.440 | 0.792 | 0.388 |
| 25 | 0.504 | 0.814 | 0.532 |
| 26 | 0.310 | 0.705 | 0.167 |
| 27 | 0.718 | 0.838 | 0.172 |
| 28 | 0.124 | 0.554 | 0.579 |
| 29 | 0.319 | 0.731 | 0.605 |
| 30 | 0.581 | 0.702 | 0.174 |
| 31 | 0.677 | 0.706 | 0.576 |
| 32 | 0.703 | 0.197 | 0.660 |
| 33 | 0.325 | 0.441 | 0.625 |
| 34 | 0.245 | 0.761 | 0.558 |
| 35 | 0.601 | 0.782 | 0.672 |
| 36 | 0.735 | 0.429 | 0.527 |
| 37 | 0.718 | 0.189 | 0.512 |
| 38 | 0.721 | 0.261 | 0.621 |
| 39 | 0.265 | 0.667 | 0.448 |
| 40 | 0.543 | 0.723 | 0.607 |
| 41 | 0.339 | 0.715 | 0.590 |
| 42 | 0.460 | 0.691 | 0.567 |
| 43 | 0.712 | 0.695 | 0.487 |
| 44 | 0.173 | 0.867 | 0.124 |
| 45 | 0.768 | 0.678 | 0.433 |
| 46 | 0.427 | 0.780 | 0.155 |
| 47 | 0.556 | 0.764 | 0.439 |
| 48 | 0.071 | 0.780 | 0.520 |
| 49 | 0.626 | 0.795 | 0.519 |
| 50 | 0.347 | 0.776 | 0.401 |
| Lilliefors p | 0.3694 | < 0.001 | 0.0292 |

**Table S11. Average Speed (AS, µm/s) of the 150 trajectories.** Last row: p-value of the Lilliefors normality test in each scenario.

| Cell | No stimuli | Galvanotaxis | Chemotaxis |
|---|---|---|---|
| 1 | 6.462 | 6.735 | 4.761 |
| 2 | 6.725 | 6.931 | 4.704 |
| 3 | 6.766 | 6.787 | 4.294 |
| 4 | 6.751 | 6.827 | 7.041 |
| 5 | 7.869 | 6.277 | 5.305 |
| 6 | 6.316 | 6.312 | 6.714 |
| 7 | 8.081 | 6.569 | 5.908 |
| 8 | 7.638 | 4.831 | 5.846 |
| 9 | 5.126 | 8.190 | 5.792 |
| 10 | 3.536 | 7.188 | 6.318 |
| 11 | 5.856 | 7.200 | 5.233 |
| 12 | 6.007 | 9.548 | 5.587 |
| 13 | 5.520 | 7.327 | 6.333 |
| 14 | 5.874 | 4.774 | 5.746 |
| 15 | 5.717 | 8.198 | 6.341 |
| 16 | 5.392 | 7.851 | 4.986 |
| 17 | 6.727 | 6.733 | 5.186 |
| 18 | 6.363 | 7.359 | 4.564 |
| 19 | 4.999 | 5.097 | 5.766 |
| 20 | 7.415 | 5.143 | 6.547 |
| 21 | 4.457 | 6.351 | 6.719 |
| 22 | 6.299 | 4.534 | 4.845 |
| 23 | 8.533 | 3.111 | 5.255 |
| 24 | 6.957 | 3.971 | 6.917 |
| 25 | 5.650 | 5.345 | 4.567 |
| 26 | 5.280 | 6.239 | 4.831 |
| 27 | 7.706 | 5.390 | 5.916 |
| 28 | 6.463 | 4.561 | 6.237 |
| 29 | 6.564 | 7.338 | 4.951 |
| 30 | 9.032 | 5.453 | 5.692 |
| 31 | 8.318 | 7.634 | 10.415 |
| 32 | 8.055 | 4.539 | 11.001 |
| 33 | 7.268 | 5.960 | 8.325 |
| 34 | 8.002 | 8.065 | 9.063 |
| 35 | 6.828 | 9.074 | 8.866 |
| 36 | 7.602 | 7.539 | 6.619 |
| 37 | 8.917 | 7.458 | 7.046 |
| 38 | 8.732 | 4.844 | 7.659 |
| 39 | 6.565 | 5.439 | 5.314 |
| 40 | 8.872 | 5.800 | 7.721 |
| 41 | 5.782 | 4.416 | 6.545 |
| 42 | 7.709 | 5.303 | 7.794 |
| 43 | 7.617 | 4.788 | 6.980 |
| 44 | 10.027 | 7.163 | 4.002 |
| 45 | 9.697 | 5.059 | 3.306 |
| 46 | 7.063 | 4.808 | 4.314 |
| 47 | 8.203 | 4.104 | 4.224 |
| 48 | 5.784 | 4.261 | 4.310 |
| 49 | 7.704 | 4.758 | 3.810 |
| 50 | 7.784 | 4.502 | 4.524 |
| Lilliefors p | > 0.5 | 0.0459 | 0.1829 |

**Table S12. Stimulus effect sizes.** For each metric, the Kruskal–Wallis statistic H with its p-value, q-value and effect size $\varepsilon^2$; and for each pair of scenarios, Cliff's δ with its BCa confidence interval and the Mann–Whitney p- and q-values. The q-values are p-values adjusted by the Benjamini–Hochberg procedure, separately over the 11 Kruskal–Wallis tests (one per metric, across the three scenarios) and over the 33 Mann–Whitney tests (one per metric and pair of scenarios). The confidence level of δ, 98.2%, keeps the false coverage rate of the intervals of the contrasts with Mann–Whitney q < 0.05 at or below 0.05 (Materials and Methods). A negative δ indicates higher values in the second scenario of the contrast. $C_{\theta\text{-}v}$ enters as its magnitude $|C_{\theta\text{-}v}|$, since a mirror reflection of a trajectory reverses its sign.

| Metric | Contrast (first vs second) | Cliff's δ | δ 98.2% CI, lower | δ 98.2% CI, upper | Mann–Whitney p | Mann–Whitney q | Kruskal–Wallis $\varepsilon^2$ | Kruskal–Wallis H | Kruskal–Wallis p | Kruskal–Wallis q |
|---|---|---|---|---|---|---|---|---|---|---|
| RMSF α | No stimuli vs galvanotaxis | -0.065 | -0.326 | 0.218 | 0.5790 | 0.7890 | 0.095 | 14.154 | $8.00 \times 10^{-4}$ | 0.0020 |
| RMSF α | No stimuli vs chemotaxis | -0.398 | -0.622 | -0.126 | $6.00 \times 10^{-4}$ | 0.0040 | 0.095 | 14.154 | $8.00 \times 10^{-4}$ | 0.0020 |
| RMSF α | Galvanotaxis vs chemotaxis | -0.350 | -0.594 | -0.089 | 0.0030 | 0.0110 | 0.095 | 14.154 | $8.00 \times 10^{-4}$ | 0.0020 |
| RMSF correlation time | No stimuli vs galvanotaxis | -0.076 | -0.344 | 0.223 | 0.5110 | 0.7890 | 0.003 | 0.505 | 0.7770 | 0.8550 |
| RMSF correlation time | No stimuli vs chemotaxis | -0.056 | -0.338 | 0.244 | 0.6330 | 0.8040 | 0.003 | 0.505 | 0.7770 | 0.8550 |
| RMSF correlation time | Galvanotaxis vs chemotaxis | 0.035 | -0.247 | 0.318 | 0.7630 | 0.8690 | 0.003 | 0.505 | 0.7770 | 0.8550 |
| DFA γ | No stimuli vs galvanotaxis | -0.286 | -0.536 | -0.016 | 0.0140 | 0.0410 | 0.054 | 8.118 | 0.0170 | 0.0320 |
| DFA γ | No stimuli vs chemotaxis | -0.049 | -0.327 | 0.210 | 0.6770 | 0.8270 | 0.054 | 8.118 | 0.0170 | 0.0320 |
| DFA γ | Galvanotaxis vs chemotaxis | 0.283 | 0.010 | 0.519 | 0.0150 | 0.0410 | 0.054 | 8.118 | 0.0170 | 0.0320 |
| MSD β | No stimuli vs galvanotaxis | -0.225 | -0.496 | 0.036 | 0.0530 | 0.1170 | 0.030 | 4.487 | 0.1060 | 0.1410 |
| MSD β | No stimuli vs chemotaxis | -0.062 | -0.335 | 0.210 | 0.5980 | 0.7890 | 0.030 | 4.487 | 0.1060 | 0.1410 |
| MSD β | Galvanotaxis vs chemotaxis | 0.190 | -0.078 | 0.456 | 0.1020 | 0.1760 | 0.030 | 4.487 | 0.1060 | 0.1410 |
| ApEn | No stimuli vs galvanotaxis | 0.194 | -0.099 | 0.461 | 0.0960 | 0.1760 | 0.029 | 4.321 | 0.1150 | 0.1410 |
| ApEn | No stimuli vs chemotaxis | 0.219 | -0.062 | 0.456 | 0.0590 | 0.1220 | 0.029 | 4.321 | 0.1150 | 0.1410 |
| ApEn | Galvanotaxis vs chemotaxis | 0.039 | -0.243 | 0.298 | 0.7380 | 0.8690 | 0.029 | 4.321 | 0.1150 | 0.1410 |
| IR | No stimuli vs galvanotaxis | -0.330 | -0.588 | -0.034 | 0.0050 | 0.0170 | 0.087 | 12.935 | 0.0020 | 0.0030 |
| IR | No stimuli vs chemotaxis | -0.022 | -0.301 | 0.235 | 0.8550 | 0.9410 | 0.087 | 12.935 | 0.0020 | 0.0030 |
| IR | Galvanotaxis vs chemotaxis | 0.392 | 0.109 | 0.623 | $7.00 \times 10^{-4}$ | 0.0040 | 0.087 | 12.935 | 0.0020 | 0.0030 |
| DR | No stimuli vs galvanotaxis | -0.634 | -0.805 | -0.377 | $5.00 \times 10^{-8}$ | $2.00 \times 10^{-6}$ | 0.268 | 39.919 | $2.00 \times 10^{-9}$ | $2.00 \times 10^{-8}$ |
| DR | No stimuli vs chemotaxis | -0.215 | -0.480 | 0.060 | 0.0640 | 0.1250 | 0.268 | 39.919 | $2.00 \times 10^{-9}$ | $2.00 \times 10^{-8}$ |
| DR | Galvanotaxis vs chemotaxis | 0.606 | 0.344 | 0.790 | $2.00 \times 10^{-7}$ | $3.00 \times 10^{-6}$ | 0.268 | 39.919 | $2.00 \times 10^{-9}$ | $2.00 \times 10^{-8}$ |
| AS | No stimuli vs galvanotaxis | 0.356 | 0.083 | 0.587 | 0.0020 | 0.0100 | 0.099 | 14.787 | $6.00 \times 10^{-4}$ | 0.0020 |
| AS | No stimuli vs chemotaxis | 0.409 | 0.138 | 0.625 | $4.00 \times 10^{-4}$ | 0.0040 | 0.099 | 14.787 | $6.00 \times 10^{-4}$ | 0.0020 |
| AS | Galvanotaxis vs chemotaxis | 0.062 | -0.226 | 0.332 | 0.5930 | 0.7890 | 0.099 | 14.787 | $6.00 \times 10^{-4}$ | 0.0020 |
| $\|C_{\theta\text{-}v}\|$ | No stimuli vs galvanotaxis | -0.069 | -0.330 | 0.187 | 0.5560 | 0.7890 | 0.032 | 4.739 | 0.0940 | 0.1410 |
| $\|C_{\theta\text{-}v}\|$ | No stimuli vs chemotaxis | -0.253 | -0.488 | 0.041 | 0.0300 | 0.0720 | 0.032 | 4.739 | 0.0940 | 0.1410 |
| $\|C_{\theta\text{-}v}\|$ | Galvanotaxis vs chemotaxis | -0.164 | -0.426 | 0.122 | 0.1590 | 0.2620 | 0.032 | 4.739 | 0.0940 | 0.1410 |
| $t_\theta$ | No stimuli vs galvanotaxis | 0.009 | -0.257 | 0.264 | 0.9420 | 0.9970 | 0.000 | 0.002 | 0.9990 | 0.9990 |
| $t_\theta$ | No stimuli vs chemotaxis | -0.001 | -0.282 | 0.264 | 0.9970 | 0.9970 | 0.000 | 0.002 | 0.9990 | 0.9990 |
| $t_\theta$ | Galvanotaxis vs chemotaxis | 0.002 | -0.272 | 0.280 | 0.9920 | 0.9970 | 0.000 | 0.002 | 0.9990 | 0.9990 |
| $AC_v(1)$ | No stimuli vs galvanotaxis | -0.251 | -0.492 | 0.045 | 0.0310 | 0.0720 | 0.138 | 20.630 | $3.00 \times 10^{-5}$ | $2.00 \times 10^{-4}$ |
| $AC_v(1)$ | No stimuli vs chemotaxis | -0.515 | -0.718 | -0.259 | $9.00 \times 10^{-6}$ | $1.00 \times 10^{-4}$ | 0.138 | 20.630 | $3.00 \times 10^{-5}$ | $2.00 \times 10^{-4}$ |
| $AC_v(1)$ | Galvanotaxis vs chemotaxis | -0.295 | -0.546 | -0.020 | 0.0110 | 0.0370 | 0.138 | 20.630 | $3.00 \times 10^{-5}$ | $2.00 \times 10^{-4}$ |

**Table S13. Pairwise dependence of the eleven metrics in each scenario.** The same 55 pairs occupy the same rows of the three subtables, ordered by the median of $R^*_n$ over the scenarios (Rank); the six couplings, dependent in all three scenarios and not definitional, are in bold, and † marks the two definitional pairs. $R^*_n$: bias-corrected squared distance correlation, with its BCa 95% confidence interval (CI); p and q: permutation p-value and Benjamini–Hochberg q-value of the distance covariance test (165 tests); r and ρ: Pearson and Spearman correlations, with their q-values; q RESET: linearity test of the regression of B on A and of A on B, A being the first metric of the pair and B the second (330 tests), a pair being nonlinear when any of its six q-values is below 0.05. No pair reversed its trend (Lind–Mehlum q ≥ 0.35 and two-lines q = 1 over the same 330 tests). The p-value $1.00 \times 10^{-4}$ is the smallest that 9,999 permutations can give, (1 + 0)/(9,999 + 1), so the 28 tests that reached it share q = $1.00 \times 10^{-4} \times 165/28 = 5.89 \times 10^{-4}$. In 72 of the 165 pair and scenario combinations, all with q ≥ 0.62, the BCa interval excludes the estimate: resampling cells with replacement duplicates some of them and adds spurious dependence, so those intervals carry no information.

**No stimuli**

| Rank | Pair (A-B) | $R^*_n$ | 95% CI | p (dCov test) | q | Pearson r | q (r) | Spearman ρ | q (ρ) | q RESET, B on A | q RESET, A on B |
|---|---|---|---|---|---|---|---|---|---|---|---|
| 1 | IR-DR† | 0.903 | [0.829, 0.940] | $1.00 \times 10^{-4}$ | $5.89 \times 10^{-4}$ | 0.955 | $1.15 \times 10^{-24}$ | 0.961 | $3.73 \times 10^{-26}$ | 0.0225 | 0.0513 |
| **2** | **DFA γ-MSD β** | **0.537** | **[0.325, 0.693]** | **$1.00 \times 10^{-4}$** | **$5.89 \times 10^{-4}$** | **0.775** | **$1.14 \times 10^{-9}$** | **0.777** | **$8.68 \times 10^{-10}$** | **0.7286** | **0.7286** |
| 3 | IR-AS† | 0.416 | [0.167, 0.608] | $1.00 \times 10^{-4}$ | $5.89 \times 10^{-4}$ | 0.654 | $3.52 \times 10^{-6}$ | 0.639 | $4.58 \times 10^{-6}$ | 0.9812 | 0.4738 |
| **4** | **DFA γ-IR** | **0.407** | **[0.195, 0.585]** | **$1.00 \times 10^{-4}$** | **$5.89 \times 10^{-4}$** | **0.636** | **$6.65 \times 10^{-6}$** | **0.668** | **$1.25 \times 10^{-6}$** | **0.2422** | **0.6223** |
| **5** | **DFA γ-DR** | **0.318** | **[0.121, 0.493]** | **$1.00 \times 10^{-4}$** | **$5.89 \times 10^{-4}$** | **0.563** | **$1.23 \times 10^{-4}$** | **0.621** | **$1.03 \times 10^{-5}$** | **0.6038** | **0.4841** |
| **6** | **MSD β-IR** | **0.392** | **[0.190, 0.523]** | **$1.00 \times 10^{-4}$** | **$5.89 \times 10^{-4}$** | **0.633** | **$7.44 \times 10^{-6}$** | **0.706** | **$1.26 \times 10^{-7}$** | **0.1276** | **0.7245** |
| 7 | MSD β-AS | 0.405 | [0.157, 0.587] | $1.00 \times 10^{-4}$ | $5.89 \times 10^{-4}$ | 0.664 | $2.39 \times 10^{-6}$ | 0.630 | $6.91 \times 10^{-6}$ | 0.9554 | 0.1276 |
| **8** | **MSD β-DR** | **0.302** | **[0.102, 0.445]** | **$1.00 \times 10^{-4}$** | **$5.89 \times 10^{-4}$** | **0.569** | **$1.07 \times 10^{-4}$** | **0.641** | **$4.34 \times 10^{-6}$** | **0.2952** | **0.9979** |
| **9** | **RMSF α-$AC_v(1)$** | **0.273** | **[0.059, 0.459]** | **$1.00 \times 10^{-4}$** | **$5.89 \times 10^{-4}$** | **0.608** | **$2.33 \times 10^{-5}$** | **0.577** | **$6.46 \times 10^{-5}$** | **0.9304** | **0.9908** |
| 10 | DFA γ-AS | 0.316 | [0.095, 0.506] | $2.00 \times 10^{-4}$ | 0.0010 | 0.581 | $6.62 \times 10^{-5}$ | 0.598 | $2.81 \times 10^{-5}$ | 0.5002 | 0.0225 |
| 11 | MSD β-$t_\theta$ | 0.025 | [-0.019, 0.075] | 0.2157 | 0.4810 | 0.222 | 0.3216 | 0.215 | 0.3247 | 0.9444 | 0.9505 |
| 12 | IR-$t_\theta$ | -0.016 | [-0.022, -0.022] | 0.8233 | 0.8722 | 0.060 | 0.8773 | 0.123 | 0.6400 | 0.8821 | 0.7590 |
| 13 | DR-AS | 0.196 | [0.028, 0.372] | $4.00 \times 10^{-4}$ | 0.0018 | 0.442 | 0.0066 | 0.451 | 0.0045 | 0.5952 | 0.5691 |
| 14 | DFA γ-$t_\theta$ | 0.012 | [-0.014, 0.021] | 0.2977 | 0.6016 | 0.090 | 0.7525 | 0.050 | 0.8785 | 0.4951 | 0.9271 |
| 15 | DFA γ-$AC_v(1)$ | 0.036 | [-0.015, 0.091] | 0.0714 | 0.2110 | -0.283 | 0.1414 | -0.318 | 0.0801 | 0.2085 | 0.5985 |
| 16 | MSD β-$AC_v(1)$ | 0.074 | [-0.007, 0.181] | 0.0154 | 0.0565 | -0.387 | 0.0213 | -0.338 | 0.0567 | 0.4951 | 0.3033 |
| 17 | DR-$t_\theta$ | -0.019 | [-0.016, -0.016] | 0.8509 | 0.8775 | 0.051 | 0.9090 | 0.077 | 0.8259 | 0.9306 | 0.8802 |
| 18 | ApEn-$AC_v(1)$ | 0.131 | [0.011, 0.277] | 0.0052 | 0.0220 | -0.363 | 0.0357 | -0.403 | 0.0154 | 0.8802 | 0.5952 |
| 19 | AS-$AC_v(1)$ | 0.224 | [0.022, 0.419] | $2.00 \times 10^{-4}$ | 0.0010 | -0.520 | $5.73 \times 10^{-4}$ | -0.501 | $9.76 \times 10^{-4}$ | 0.8802 | 0.6048 |
| 20 | IR-$AC_v(1)$ | 0.046 | [-0.025, 0.151] | 0.0528 | 0.1675 | -0.352 | 0.0423 | -0.285 | 0.1389 | 0.8821 | 0.7963 |
| 21 | RMSF correlation time-$t_\theta$ | 0.055 | [-0.003, 0.135] | 0.0605 | 0.1883 | -0.244 | 0.2400 | -0.190 | 0.3839 | 0.1806 | 0.9925 |
| 22 | RMSF α-AS | -0.035 | [-0.026, -0.026] | 0.9278 | 0.9335 | -0.023 | 0.9543 | 0.010 | 0.9796 | 0.7245 | 0.7770 |
| 23 | ApEn-$t_\theta$ | 0.237 | [0.069, 0.388] | $2.00 \times 10^{-4}$ | 0.0010 | -0.527 | $4.64 \times 10^{-4}$ | -0.521 | $4.97 \times 10^{-4}$ | 0.9979 | 0.5205 |
| 24 | RMSF α-$t_\theta$ | 0.034 | [-0.013, 0.079] | 0.1321 | 0.3302 | 0.159 | 0.5722 | 0.250 | 0.2270 | 0.8630 | 0.5205 |
| 25 | $C_{\theta\text{-}v}$-$t_\theta$ | -0.016 | [-0.019, -0.019] | 0.6290 | 0.8154 | 0.111 | 0.6852 | 0.083 | 0.8147 | 0.8568 | 0.8014 |
| 26 | MSD β-ApEn | 0.023 | [-0.023, 0.071] | 0.1792 | 0.4165 | -0.246 | 0.2382 | -0.228 | 0.2931 | 0.8362 | 0.9979 |

| Rank | Pair (A-B) | $R^*_n$ | 95% CI | p (dCov test) | q | Pearson r | q (r) | Spearman ρ | q (ρ) | q RESET, B on A | q RESET, A on B |
|---|---|---|---|---|---|---|---|---|---|---|---|
| 27 | RMSF α-DFA γ | -0.012 | [-0.024, -0.017] | 0.4515 | 0.7163 | 0.052 | 0.9090 | -0.005 | 0.9964 | 0.5952 | 0.2362 |
| 28 | RMSF α-ApEn | 0.041 | [-0.008, 0.093] | 0.0934 | 0.2526 | -0.218 | 0.3361 | -0.232 | 0.2832 | 0.7770 | 0.8821 |
| 29 | DR-$AC_v(1)$ | 0.005 | [-0.027, 0.036] | 0.2587 | 0.5544 | -0.261 | 0.1939 | -0.222 | 0.3070 | 0.9979 | 0.7286 |
| 30 | RMSF correlation time-ApEn | 0.004 | [-0.017, 0.008] | 0.3290 | 0.6240 | -0.015 | 0.9621 | 0.032 | 0.9235 | 0.0283 | 0.8568 |
| 31 | MSD β-$C_{\theta\text{-}v}$ | -0.015 | [-0.022, -0.022] | 0.5659 | 0.7717 | 0.111 | 0.6852 | -0.018 | 0.9673 | 0.7247 | 0.5952 |
| 32 | RMSF correlation time-AS | -0.021 | [-0.023, -0.023] | 0.8054 | 0.8722 | 0.061 | 0.8773 | 0.000 | 0.9981 | 0.4951 | 0.6967 |
| 33 | AS-$t_\theta$ | 0.000 | [-0.015, -0.009] | 0.5083 | 0.7494 | 0.096 | 0.7320 | 0.084 | 0.8147 | 0.8821 | 0.5597 |
| 34 | DFA γ-ApEn | -0.006 | [-0.023, -0.004] | 0.5132 | 0.7494 | -0.165 | 0.5482 | -0.126 | 0.6339 | 0.9505 | 0.9913 |
| 35 | RMSF correlation time-$AC_v(1)$ | 0.012 | [-0.012, 0.023] | 0.2123 | 0.4799 | -0.176 | 0.4954 | -0.199 | 0.3642 | 0.4772 | 0.5952 |
| 36 | ApEn-AS | -0.001 | [-0.009, -0.005] | 0.4769 | 0.7416 | -0.138 | 0.6222 | -0.113 | 0.6915 | 0.8677 | 0.8630 |
| 37 | AS-$C_{\theta\text{-}v}$ | -0.004 | [-0.023, -0.005] | 0.3961 | 0.7069 | 0.003 | 0.9850 | -0.079 | 0.8259 | 0.5952 | 0.4132 |
| 38 | RMSF correlation time-DR | -0.005 | [-0.022, 0.004] | 0.4206 | 0.7082 | 0.214 | 0.3461 | 0.227 | 0.2931 | 0.5985 | 0.8821 |
| 39 | RMSF correlation time-$C_{\theta\text{-}v}$ | -0.009 | [-0.025, -0.015] | 0.5257 | 0.7505 | -0.102 | 0.7228 | -0.039 | 0.9118 | 0.8164 | 0.8176 |
| 40 | RMSF α-MSD β | -0.024 | [-0.016, -0.016] | 0.7408 | 0.8722 | -0.077 | 0.8144 | -0.031 | 0.9235 | 0.5544 | 0.5952 |
| 41 | RMSF correlation time-IR | -0.010 | [-0.020, -0.011] | 0.5126 | 0.7494 | 0.187 | 0.4510 | 0.171 | 0.4399 | 0.3996 | 0.7245 |
| 42 | RMSF correlation time-DFA γ | 0.029 | [-0.012, 0.085] | 0.1248 | 0.3181 | 0.174 | 0.4965 | 0.175 | 0.4364 | 0.6122 | 0.3996 |
| 43 | IR-$C_{\theta\text{-}v}$ | -0.015 | [-0.017, -0.017] | 0.6794 | 0.8366 | 0.013 | 0.9627 | -0.027 | 0.9262 | 0.8821 | 0.4180 |
| 44 | $t_\theta$-$AC_v(1)$ | 0.146 | [0.034, 0.315] | 0.0036 | 0.0156 | 0.325 | 0.0719 | 0.408 | 0.0137 | 0.4951 | 0.8778 |
| 45 | RMSF α-IR | -0.014 | [-0.025, -0.017] | 0.5322 | 0.7505 | 0.046 | 0.9090 | 0.046 | 0.8889 | 0.8411 | 0.7286 |
| 46 | RMSF α-$C_{\theta\text{-}v}$ | -0.014 | [-0.031, -0.018] | 0.5786 | 0.7825 | -0.041 | 0.9302 | 0.002 | 0.9981 | 0.2674 | 0.4841 |
| 47 | DFA γ-$C_{\theta\text{-}v}$ | -0.014 | [-0.025, -0.021] | 0.6058 | 0.8055 | 0.037 | 0.9415 | -0.055 | 0.8785 | 0.5575 | 0.6048 |
| 48 | ApEn-$C_{\theta\text{-}v}$ | -0.014 | [-0.015, -0.015] | 0.6424 | 0.8154 | 0.029 | 0.9484 | 0.070 | 0.8447 | 0.6054 | 0.9979 |
| 49 | $C_{\theta\text{-}v}$-$AC_v(1)$ | -0.021 | [-0.022, -0.022] | 0.7977 | 0.8722 | -0.025 | 0.9543 | 0.020 | 0.9617 | 0.5952 | 0.9271 |
| 50 | ApEn-DR | -0.012 | [-0.025, -0.017] | 0.6555 | 0.8194 | -0.091 | 0.7525 | -0.129 | 0.6209 | 0.9935 | 0.8362 |
| 51 | RMSF α-RMSF correlation time | -0.018 | [-0.019, -0.019] | 0.7600 | 0.8722 | 0.045 | 0.9090 | 0.016 | 0.9715 | 0.9979 | 0.7288 |
| 52 | RMSF α-DR | -0.018 | [-0.025, -0.025] | 0.6525 | 0.8194 | 0.028 | 0.9484 | 0.044 | 0.8904 | 0.8176 | 0.8362 |
| 53 | ApEn-IR | -0.021 | [-0.024, -0.024] | 0.8410 | 0.8727 | -0.111 | 0.6852 | -0.153 | 0.5133 | 0.9979 | 0.8014 |
| 54 | DR-$C_{\theta\text{-}v}$ | -0.021 | [-0.021, -0.021] | 0.7654 | 0.8722 | 0.045 | 0.9090 | 0.027 | 0.9262 | 0.8568 | 0.4915 |
| 55 | RMSF correlation time-MSD β | -0.024 | [-0.023, -0.023] | 0.8218 | 0.8722 | 0.060 | 0.8773 | 0.060 | 0.8705 | 0.7366 | 0.6592 |

**Galvanotaxis**

| Rank | Pair (A-B) | $R^*_n$ | 95% CI | p (dCov test) | q | Pearson r | q (r) | Spearman ρ | q (ρ) | q RESET, B on A | q RESET, A on B |
|---|---|---|---|---|---|---|---|---|---|---|---|
| 1 | IR-DR[†] | 0.437 | [0.209, 0.640] | $1.00 \times 10^{-4}$ | $5.89 \times 10^{-4}$ | 0.732 | $3.18 \times 10^{-8}$ | 0.659 | $1.91 \times 10^{-6}$ | 0.0170 | 0.9304 |
| **2** | **DFA γ-MSD β** | **0.670** | **[0.418, 0.823]** | $\mathbf{1.00 \times 10^{-4}}$ | $\mathbf{5.89 \times 10^{-4}}$ | **0.898** | $\mathbf{8.20 \times 10^{-17}}$ | **0.868** | $\mathbf{2.83 \times 10^{-14}}$ | **0.0106** | **0.9908** |
| 3 | IR-AS[†] | 0.383 | [0.150, 0.594] | $1.00 \times 10^{-4}$ | $5.89 \times 10^{-4}$ | 0.614 | $1.88 \times 10^{-5}$ | 0.617 | $1.21 \times 10^{-5}$ | 0.2479 | 0.9979 |

| Rank | Pair (A-B) | $R^*_n$ | 95% CI | p (dCov test) | q | Pearson r | q (r) | Spearman ρ | q (ρ) | q RESET, B on A | q RESET, A on B |
|---|---|---|---|---|---|---|---|---|---|---|---|
| **4** | **DFA γ-IR** | **0.357** | **[0.126, 0.507]** | **$1.00 \times 10^{-4}$** | **$5.89 \times 10^{-4}$** | **0.643** | **$5.16 \times 10^{-6}$** | **0.684** | **$5.19 \times 10^{-7}$** | **0.1593** | **0.0225** |
| **5** | **DFA γ-DR** | **0.566** | **[0.315, 0.745]** | **$1.00 \times 10^{-4}$** | **$5.89 \times 10^{-4}$** | **0.765** | **$2.25 \times 10^{-9}$** | **0.767** | **$2.02 \times 10^{-9}$** | **0.4951** | **0.9163** |
| **6** | **MSD β-IR** | **0.339** | **[0.081, 0.486]** | **$1.00 \times 10^{-4}$** | **$5.89 \times 10^{-4}$** | **0.595** | **$3.83 \times 10^{-5}$** | **0.654** | **$2.43 \times 10^{-6}$** | **0.1276** | **0.1309** |
| 7 | MSD β-AS | -0.015 | [-0.024, -0.021] | 0.6102 | 0.8055 | 0.154 | 0.5722 | 0.113 | 0.6915 | 0.5544 | 0.9505 |
| **8** | **MSD β-DR** | **0.601** | **[0.361, 0.733]** | **$1.00 \times 10^{-4}$** | **$5.89 \times 10^{-4}$** | **0.702** | **$2.46 \times 10^{-7}$** | **0.862** | **$3.74 \times 10^{-14}$** | **0.1331** | **0.9505** |
| **9** | **RMSF α-AC$_v$(1)** | **0.186** | **[0.029, 0.343]** | **$5.00 \times 10^{-4}$** | **0.0022** | **0.470** | **0.0030** | **0.521** | **$4.97 \times 10^{-4}$** | **0.5544** | **0.8677** |
| 10 | DFA γ-AS | -0.023 | [-0.029, -0.029] | 0.7928 | 0.8722 | 0.124 | 0.6725 | 0.184 | 0.3985 | 0.5952 | 0.9979 |
| 11 | MSD β-t$_θ$ | 0.238 | [0.026, 0.403] | $4.00 \times 10^{-4}$ | 0.0018 | 0.427 | 0.0090 | 0.532 | $3.44 \times 10^{-4}$ | 0.0767 | 0.7288 |
| 12 | IR-t$_θ$ | 0.117 | [0.000, 0.236] | 0.0056 | 0.0231 | 0.432 | 0.0082 | 0.422 | 0.0099 | 0.9591 | 0.9935 |
| 13 | DR-AS | -0.020 | [-0.024, -0.024] | 0.7735 | 0.8722 | -0.062 | 0.8773 | -0.052 | 0.8785 | 0.9395 | 0.9979 |
| 14 | DFA γ-t$_θ$ | 0.101 | [-0.008, 0.221] | 0.0193 | 0.0663 | 0.304 | 0.1007 | 0.389 | 0.0207 | 0.5952 | 0.8927 |
| 15 | DFA γ-AC$_v$(1) | 0.108 | [0.008, 0.188] | 0.0099 | 0.0398 | -0.111 | 0.6852 | -0.401 | 0.0156 | 0.0225 | 0.8723 |
| 16 | MSD β-AC$_v$(1) | 0.046 | [-0.005, 0.072] | 0.0716 | 0.2110 | -0.030 | 0.9484 | -0.252 | 0.2270 | 0.1990 | 0.9238 |
| 17 | DR-t$_θ$ | 0.252 | [0.057, 0.423] | $1.00 \times 10^{-4}$ | $5.89 \times 10^{-4}$ | 0.441 | 0.0066 | 0.555 | $1.47 \times 10^{-4}$ | 0.1276 | 0.7197 |
| 18 | ApEn-AC$_v$(1) | 0.056 | [-0.009, 0.154] | 0.0429 | 0.1388 | -0.309 | 0.0932 | -0.208 | 0.3436 | 0.8176 | 0.5823 |
| 19 | AS-AC$_v$(1) | 0.048 | [-0.014, 0.148] | 0.0774 | 0.2205 | -0.286 | 0.1371 | -0.278 | 0.1547 | 0.8768 | 0.9124 |
| 20 | IR-AC$_v$(1) | 0.094 | [-0.002, 0.213] | 0.0123 | 0.0472 | -0.354 | 0.0420 | -0.335 | 0.0581 | 0.9979 | 0.5952 |
| 21 | RMSF correlation time-t$_θ$ | 0.027 | [-0.014, 0.077] | 0.0917 | 0.2522 | -0.334 | 0.0616 | -0.234 | 0.2806 | 0.4159 | 0.7286 |
| 22 | RMSF α-AS | 0.033 | [-0.016, 0.089] | 0.1253 | 0.3181 | -0.240 | 0.2526 | -0.174 | 0.4364 | 0.9979 | 0.9863 |
| 23 | ApEn-t$_θ$ | 0.014 | [-0.028, 0.029] | 0.1102 | 0.2933 | -0.268 | 0.1760 | -0.147 | 0.5317 | 0.7288 | 0.6223 |
| 24 | RMSF α-t$_θ$ | 0.013 | [-0.021, 0.031] | 0.2858 | 0.5895 | -0.132 | 0.6393 | -0.196 | 0.3717 | 0.9979 | 0.7288 |
| 25 | C$_{θ\text{-}v}$-t$_θ$ | 0.029 | [-0.015, 0.048] | 0.1395 | 0.3435 | -0.153 | 0.5722 | -0.217 | 0.3221 | 0.9101 | 0.8014 |
| 26 | MSD β-ApEn | -0.015 | [-0.020, -0.020] | 0.7301 | 0.8722 | -0.010 | 0.9627 | -0.088 | 0.8024 | 0.8014 | 0.8821 |
| 27 | RMSF α-DFA γ | 0.007 | [-0.021, 0.015] | 0.3511 | 0.6509 | -0.100 | 0.7228 | -0.245 | 0.2410 | 0.8411 | 0.1802 |
| 28 | RMSF α-ApEn | 0.005 | [-0.016, 0.007] | 0.4130 | 0.7082 | -0.129 | 0.6536 | -0.124 | 0.6360 | 0.5952 | 0.9304 |
| 29 | DR-AC$_v$(1) | 0.029 | [0.000, 0.035] | 0.0891 | 0.2492 | -0.200 | 0.3938 | -0.308 | 0.0954 | 0.5327 | 0.5327 |
| 30 | RMSF correlation time-ApEn | -0.025 | [-0.024, -0.024] | 0.7814 | 0.8722 | 0.099 | 0.7235 | -0.001 | 0.9981 | 0.9271 | 0.5544 |
| 31 | MSD β-C$_{θ\text{-}v}$ | 0.002 | [-0.020, 0.001] | 0.3104 | 0.6171 | 0.049 | 0.9090 | -0.203 | 0.3568 | 0.4841 | 0.8821 |
| 32 | RMSF correlation time-AS | 0.033 | [-0.018, 0.102] | 0.1212 | 0.3174 | 0.213 | 0.3461 | 0.198 | 0.3642 | 0.5952 | 0.9935 |
| 33 | AS-t$_θ$ | -0.015 | [-0.023, -0.023] | 0.6754 | 0.8366 | 0.135 | 0.6332 | 0.061 | 0.8705 | 0.9505 | 0.9163 |
| 34 | DFA γ-ApEn | 0.000 | [-0.015, -0.002] | 0.3947 | 0.7069 | 0.141 | 0.6157 | -0.061 | 0.8705 | 0.4915 | 0.9979 |
| 35 | RMSF correlation time-AC$_v$(1) | -0.025 | [-0.024, -0.024] | 0.8748 | 0.8965 | -0.030 | 0.9484 | -0.030 | 0.9252 | 0.5304 | 0.8802 |
| 36 | ApEn-AS | 0.002 | [-0.013, 0.001] | 0.4461 | 0.7146 | -0.149 | 0.5840 | -0.185 | 0.3981 | 0.8362 | 0.9306 |
| 37 | AS-C$_{θ\text{-}v}$ | -0.021 | [-0.033, -0.024] | 0.7580 | 0.8722 | 0.118 | 0.6852 | 0.162 | 0.4729 | 0.5952 | 0.9505 |
| 38 | RMSF correlation time-DR | 0.015 | [-0.019, 0.039] | 0.2307 | 0.5009 | -0.121 | 0.6751 | -0.203 | 0.3568 | 0.5611 | 0.5594 |

| Rank | Pair (A-B) | $R^*_n$ | 95% CI | p (dCov test) | q | Pearson r | q (r) | Spearman ρ | q (ρ) | q RESET, B on A | q RESET, A on B |
|---|---|---|---|---|---|---|---|---|---|---|---|
| 39 | RMSF correlation time-$C_{\theta\text{-}v}$ | -0.001 | [-0.024, 0.012] | 0.4356 | 0.7116 | -0.154 | 0.5722 | -0.170 | 0.4407 | 0.9908 | 0.9505 |
| 40 | RMSF α-MSD β | -0.006 | [-0.023, -0.009] | 0.5046 | 0.7494 | -0.118 | 0.6852 | -0.193 | 0.3723 | 0.8927 | 0.4180 |
| 41 | RMSF correlation time-IR | -0.008 | [-0.017, -0.009] | 0.5230 | 0.7505 | 0.082 | 0.7895 | 0.107 | 0.7121 | 0.3068 | 0.8014 |
| 42 | RMSF correlation time-DFA γ | -0.009 | [-0.017, -0.014] | 0.5292 | 0.7505 | -0.010 | 0.9627 | 0.093 | 0.7811 | 0.7286 | 0.8176 |
| 43 | IR-$C_{\theta\text{-}v}$ | -0.007 | [-0.018, -0.013] | 0.4451 | 0.7146 | 0.036 | 0.9439 | 0.108 | 0.7121 | 0.5257 | 0.8927 |
| 44 | $t_\theta$-$AC_v(1)$ | -0.013 | [-0.025, -0.018] | 0.5642 | 0.7717 | 0.037 | 0.9415 | 0.095 | 0.7780 | 0.5952 | 0.8821 |
| 45 | RMSF α-IR | 0.043 | [-0.012, 0.128] | 0.0641 | 0.1959 | -0.323 | 0.0729 | -0.227 | 0.2931 | 0.9505 | 0.7197 |
| 46 | RMSF α-$C_{\theta\text{-}v}$ | 0.072 | [-0.023, 0.188] | 0.0175 | 0.0614 | 0.401 | 0.0161 | 0.363 | 0.0360 | 0.9960 | 0.9159 |
| 47 | DFA γ-$C_{\theta\text{-}v}$ | -0.031 | [-0.028, -0.028] | 0.8187 | 0.8722 | 0.020 | 0.9565 | -0.139 | 0.5732 | 0.5952 | 0.8927 |
| 48 | ApEn-$C_{\theta\text{-}v}$ | -0.007 | [-0.021, -0.016] | 0.4185 | 0.7082 | 0.011 | 0.9627 | 0.074 | 0.8293 | 0.3767 | 0.5544 |
| 49 | $C_{\theta\text{-}v}$-$AC_v(1)$ | 0.000 | [-0.028, 0.014] | 0.3356 | 0.6292 | 0.202 | 0.3911 | 0.208 | 0.3436 | 0.9908 | 0.9163 |
| 50 | ApEn-DR | -0.018 | [-0.023, -0.023] | 0.4752 | 0.7416 | 0.074 | 0.8235 | 0.011 | 0.9796 | 0.8821 | 0.4738 |
| 51 | RMSF α-RMSF correlation time | 0.027 | [-0.006, 0.049] | 0.1862 | 0.4267 | -0.138 | 0.6222 | -0.157 | 0.4979 | 0.5544 | 0.9979 |
| 52 | RMSF α-DR | 0.007 | [-0.010, 0.007] | 0.3189 | 0.6190 | -0.199 | 0.3938 | -0.200 | 0.3642 | 0.8821 | 0.6223 |
| 53 | ApEn-IR | -0.006 | [-0.021, -0.015] | 0.4933 | 0.7467 | -0.050 | 0.9090 | -0.086 | 0.8069 | 0.9979 | 0.4937 |
| 54 | DR-$C_{\theta\text{-}v}$ | 0.000 | [-0.018, -0.018] | 0.4279 | 0.7116 | -0.029 | 0.9484 | -0.148 | 0.5312 | 0.7286 | 0.8821 |
| 55 | RMSF correlation time-MSD β | -0.024 | [-0.029, -0.029] | 0.8034 | 0.8722 | -0.125 | 0.6720 | -0.028 | 0.9262 | 0.8176 | 0.9071 |

**Chemotaxis**

| Rank | Pair (A-B) | $R^*_n$ | 95% CI | p (dCov test) | q | Pearson r | q (r) | Spearman ρ | q (ρ) | q RESET, B on A | q RESET, A on B |
|---|---|---|---|---|---|---|---|---|---|---|---|
| 1 | IR-DR[†] | 0.686 | [0.541, 0.800] | $1.00 \times 10^{-4}$ | $5.89 \times 10^{-4}$ | 0.810 | $3.55 \times 10^{-11}$ | 0.865 | $3.01 \times 10^{-14}$ | $7.17 \times 10^{-5}$ | 0.4841 |
| **2** | **DFA γ-MSD β** | **0.605** | **[0.361, 0.747]** | **$1.00 \times 10^{-4}$** | **$5.89 \times 10^{-4}$** | **0.871** | **$1.11 \times 10^{-14}$** | **0.842** | **$6.14 \times 10^{-13}$** | **0.4951** | **$5.05 \times 10^{-4}$** |
| 3 | IR-AS[†] | 0.579 | [0.319, 0.749] | $1.00 \times 10^{-4}$ | $5.89 \times 10^{-4}$ | 0.824 | $8.62 \times 10^{-12}$ | 0.750 | $7.56 \times 10^{-9}$ | 0.0767 | 0.9488 |
| **4** | **DFA γ-IR** | **0.457** | **[0.261, 0.606]** | **$1.00 \times 10^{-4}$** | **$5.89 \times 10^{-4}$** | **0.599** | **$3.39 \times 10^{-5}$** | **0.743** | **$1.15 \times 10^{-8}$** | **0.0225** | **0.0225** |
| **5** | **DFA γ-DR** | **0.393** | **[0.143, 0.594]** | **$1.00 \times 10^{-4}$** | **$5.89 \times 10^{-4}$** | **0.642** | **$5.16 \times 10^{-6}$** | **0.661** | **$1.77 \times 10^{-6}$** | **0.2020** | **0.3060** |
| **6** | **MSD β-IR** | **0.426** | **[0.170, 0.580]** | **$1.00 \times 10^{-4}$** | **$5.89 \times 10^{-4}$** | **0.654** | **$3.52 \times 10^{-6}$** | **0.730** | **$2.77 \times 10^{-8}$** | **0.0225** | **0.6054** |
| 7 | MSD β-AS | 0.374 | [0.137, 0.529] | $1.00 \times 10^{-4}$ | $5.89 \times 10^{-4}$ | 0.566 | $1.13 \times 10^{-4}$ | 0.648 | $3.10 \times 10^{-6}$ | 0.0677 | 0.8014 |
| **8** | **MSD β-DR** | **0.215** | **[0.041, 0.386]** | **$2.00 \times 10^{-4}$** | **0.0010** | **0.568** | **$1.07 \times 10^{-4}$** | **0.582** | **$5.21 \times 10^{-5}$** | **0.7288** | **0.5544** |
| **9** | **RMSF α-$AC_v(1)$** | **0.359** | **[0.105, 0.539]** | **$1.00 \times 10^{-4}$** | **$5.89 \times 10^{-4}$** | **0.652** | **$3.75 \times 10^{-6}$** | **0.634** | **$5.83 \times 10^{-6}$** | **0.9713** | **0.1276** |
| 10 | DFA γ-AS | 0.243 | [0.041, 0.368] | $3.00 \times 10^{-4}$ | 0.0015 | 0.426 | 0.0090 | 0.575 | $6.68 \times 10^{-5}$ | 0.1276 | 0.8464 |
| 11 | MSD β-$t_\theta$ | 0.506 | [0.312, 0.644] | $1.00 \times 10^{-4}$ | $5.89 \times 10^{-4}$ | 0.590 | $4.67 \times 10^{-5}$ | 0.731 | $2.65 \times 10^{-8}$ | 0.1331 | 0.2085 |
| 12 | IR-$t_\theta$ | 0.304 | [0.099, 0.470] | $1.00 \times 10^{-4}$ | $5.89 \times 10^{-4}$ | 0.529 | $4.49 \times 10^{-4}$ | 0.587 | $4.42 \times 10^{-5}$ | 0.5985 | 0.5952 |
| 13 | DR-AS | 0.102 | [0.017, 0.191] | 0.0161 | 0.0578 | 0.355 | 0.0420 | 0.367 | 0.0334 | 0.5967 | 0.9979 |
| 14 | DFA γ-$t_\theta$ | 0.260 | [0.071, 0.459] | $2.00 \times 10^{-4}$ | 0.0010 | 0.387 | 0.0213 | 0.603 | $2.30 \times 10^{-5}$ | 0.0650 | 0.6406 |
| 15 | DFA γ-$AC_v(1)$ | 0.075 | [-0.011, 0.195] | 0.0108 | 0.0424 | -0.398 | 0.0168 | -0.338 | 0.0567 | 0.9979 | $5.05 \times 10^{-4}$ |

| Rank | Pair (A-B) | $R^*_n$ | 95% CI | p (dCov test) | q | Pearson r | q (r) | Spearman ρ | q (ρ) | q RESET, B on A | q RESET, A on B |
|---|---|---|---|---|---|---|---|---|---|---|---|
| 16 | MSD β-$AC_v(1)$ | 0.086 | [-0.016, 0.220] | 0.0128 | 0.0480 | -0.411 | 0.0127 | -0.338 | 0.0567 | 0.9979 | 0.0283 |
| 17 | DR-$t_θ$ | 0.062 | [-0.014, 0.143] | 0.0428 | 0.1388 | 0.249 | 0.2310 | 0.345 | 0.0521 | 0.5952 | 0.8821 |
| 18 | ApEn-$AC_v(1)$ | 0.056 | [-0.013, 0.170] | 0.0265 | 0.0892 | -0.412 | 0.0127 | -0.258 | 0.2127 | 0.4951 | 0.5952 |
| 19 | AS-$AC_v(1)$ | 0.016 | [-0.019, 0.041] | 0.2301 | 0.5009 | -0.181 | 0.4702 | -0.188 | 0.3906 | 0.8821 | 0.8362 |
| 20 | IR-$AC_v(1)$ | 0.012 | [-0.009, 0.016] | 0.2840 | 0.5895 | -0.183 | 0.4635 | -0.171 | 0.4399 | 0.8927 | 0.8176 |
| 21 | RMSF correlation time-$t_θ$ | -0.029 | [-0.025, -0.025] | 0.8305 | 0.8727 | 0.021 | 0.9543 | -0.051 | 0.8785 | 0.9488 | 0.6223 |
| 22 | RMSF α-AS | 0.016 | [-0.011, 0.027] | 0.2990 | 0.6016 | -0.156 | 0.5722 | -0.193 | 0.3723 | 0.7770 | 0.8176 |
| 23 | ApEn-$t_θ$ | -0.015 | [-0.024, -0.023] | 0.5641 | 0.7717 | -0.101 | 0.7228 | -0.077 | 0.8259 | 0.5952 | 0.9071 |
| 24 | RMSF α-$t_θ$ | 0.012 | [-0.016, 0.026] | 0.2763 | 0.5845 | -0.114 | 0.6852 | -0.208 | 0.3436 | 0.4951 | 0.9979 |
| 25 | $C_{θ-v}$-$t_θ$ | 0.009 | [-0.013, 0.015] | 0.3176 | 0.6190 | -0.147 | 0.5934 | -0.182 | 0.4032 | 0.5952 | 0.8014 |
| 26 | MSD β-ApEn | 0.009 | [-0.017, 0.010] | 0.1682 | 0.4022 | 0.155 | 0.5722 | -0.057 | 0.8785 | 0.4951 | 0.5205 |
| 27 | RMSF α-DFA γ | 0.011 | [-0.022, 0.022] | 0.4317 | 0.7116 | -0.115 | 0.6852 | -0.217 | 0.3221 | 0.9979 | 0.8104 |
| 28 | RMSF α-ApEn | -0.020 | [-0.020, -0.020] | 0.3237 | 0.6211 | -0.271 | 0.1709 | -0.032 | 0.9235 | 0.4841 | 0.0767 |
| 29 | DR-$AC_v(1)$ | -0.007 | [-0.012, -0.011] | 0.5569 | 0.7717 | -0.155 | 0.5722 | -0.106 | 0.7178 | 0.9505 | 0.5985 |
| 30 | RMSF correlation time-ApEn | 0.004 | [-0.009, 0.003] | 0.4832 | 0.7416 | -0.134 | 0.6332 | -0.094 | 0.7780 | 0.9935 | 0.9069 |
| 31 | MSD β-$C_{θ-v}$ | 0.025 | [-0.013, 0.056] | 0.1580 | 0.3834 | -0.109 | 0.6911 | -0.251 | 0.2270 | 0.5104 | 0.5544 |
| 32 | RMSF correlation time-AS | 0.002 | [-0.024, 0.010] | 0.4854 | 0.7416 | 0.024 | 0.9543 | -0.053 | 0.8785 | 0.5048 | 0.9979 |
| 33 | AS-$t_θ$ | 0.440 | [0.190, 0.597] | $1.00 \times 10^{-4}$ | $5.89 \times 10^{-4}$ | 0.647 | $4.64 \times 10^{-6}$ | 0.723 | $4.29 \times 10^{-8}$ | 0.4951 | 0.5594 |
| 34 | DFA γ-ApEn | 0.008 | [-0.016, 0.011] | 0.1760 | 0.4149 | 0.213 | 0.3461 | -0.045 | 0.8904 | 0.5544 | 0.5985 |
| 35 | RMSF correlation time-$AC_v(1)$ | -0.001 | [-0.015, -0.001] | 0.3995 | 0.7069 | 0.005 | 0.9803 | -0.010 | 0.9796 | 0.6851 | 0.6048 |
| 36 | ApEn-AS | -0.023 | [-0.021, -0.021] | 0.8192 | 0.8722 | 0.034 | 0.9461 | -0.049 | 0.8785 | 0.8821 | 0.8677 |
| 37 | AS-$C_{θ-v}$ | 0.003 | [-0.010, -0.001] | 0.4076 | 0.7079 | -0.091 | 0.7525 | -0.071 | 0.8446 | 0.9304 | 0.8176 |
| 38 | RMSF correlation time-DR | -0.006 | [-0.024, -0.004] | 0.3673 | 0.6734 | 0.100 | 0.7228 | 0.078 | 0.8259 | 0.7286 | 0.5544 |
| 39 | RMSF correlation time-$C_{θ-v}$ | -0.006 | [-0.024, -0.008] | 0.6010 | 0.8055 | 0.022 | 0.9543 | 0.065 | 0.8638 | 0.7770 | 0.5952 |
| 40 | RMSF α-MSD β | 0.052 | [-0.009, 0.127] | 0.0775 | 0.2205 | -0.198 | 0.3938 | -0.301 | 0.1068 | 0.6888 | 0.8014 |
| 41 | RMSF correlation time-IR | 0.003 | [-0.015, 0.002] | 0.4027 | 0.7069 | 0.081 | 0.7915 | 0.069 | 0.8452 | 0.5952 | 0.9908 |
| 42 | RMSF correlation time-DFA γ | -0.036 | [-0.031, -0.031] | 0.7576 | 0.8722 | -0.123 | 0.6744 | 0.033 | 0.9235 | 0.7963 | 0.7720 |
| 43 | IR-$C_{θ-v}$ | -0.012 | [-0.030, -0.015] | 0.6270 | 0.8154 | -0.084 | 0.7874 | -0.051 | 0.8785 | 0.8162 | 0.5985 |
| 44 | $t_θ$-$AC_v(1)$ | -0.018 | [-0.021, -0.021] | 0.6888 | 0.8419 | -0.072 | 0.8304 | -0.090 | 0.7913 | 0.8821 | 0.5985 |
| 45 | RMSF α-IR | -0.016 | [-0.020, -0.020] | 0.7691 | 0.8722 | -0.112 | 0.6852 | -0.133 | 0.5992 | 0.8802 | 0.9261 |
| 46 | RMSF α-$C_{θ-v}$ | -0.027 | [-0.026, -0.026] | 0.9166 | 0.9278 | -0.014 | 0.9621 | -0.042 | 0.9000 | 0.8802 | 0.9870 |
| 47 | DFA γ-$C_{θ-v}$ | -0.014 | [-0.020, -0.020] | 0.6378 | 0.8154 | 0.026 | 0.9541 | -0.164 | 0.4667 | 0.7770 | 0.5487 |
| 48 | ApEn-$C_{θ-v}$ | -0.025 | [-0.023, -0.023] | 0.8207 | 0.8722 | 0.008 | 0.9699 | 0.037 | 0.9126 | 0.8014 | 0.8677 |
| 49 | $C_{θ-v}$-$AC_v(1)$ | -0.016 | [-0.025, -0.020] | 0.6401 | 0.8154 | -0.145 | 0.5958 | -0.149 | 0.5274 | 0.9124 | 0.9271 |
| 50 | ApEn-DR | -0.029 | [-0.024, -0.024] | 0.9007 | 0.9174 | 0.066 | 0.8643 | 0.002 | 0.9981 | 0.8162 | 0.9812 |

| Rank | Pair (A-B) | $R^*_n$ | 95% CI | p (dCov test) | q | Pearson r | q (r) | Spearman ρ | q (ρ) | q RESET, B on A | q RESET, A on B |
|---|---|---|---|---|---|---|---|---|---|---|---|
| 51 | RMSF α-RMSF correlation time | -0.020 | [-0.022, -0.022] | 0.8059 | 0.8722 | 0.016 | 0.9621 | 0.007 | 0.9894 | 0.7770 | 0.3502 |
| 52 | RMSF α-DR | -0.033 | [-0.024, -0.024] | 0.9798 | 0.9798 | -0.046 | 0.9090 | -0.075 | 0.8293 | 0.9505 | 0.9908 |
| 53 | ApEn-IR | -0.023 | [-0.021, -0.021] | 0.7914 | 0.8722 | 0.058 | 0.8809 | -0.012 | 0.9796 | 0.8677 | 0.9271 |
| 54 | DR-$C_{\theta\text{-}v}$ | -0.023 | [-0.028, -0.028] | 0.8246 | 0.8722 | -0.014 | 0.9621 | -0.050 | 0.8785 | 0.7770 | 0.5544 |
| 55 | RMSF correlation time-MSD β | -0.028 | [-0.024, -0.024] | 0.8358 | 0.8727 | -0.149 | 0.5840 | -0.060 | 0.8705 | 0.6954 | 0.8821 |

**Table S14. Ranking of the four candidate forms by AICc.** Borda count over the 36 contests (6 couplings × 3 scenarios × 2 fit directions), with the number of first places, the mean AICc difference from the best-ranked candidate, the mean Akaike weight and the mean rank of each form.

| Rank | Candidate form | Mean Borda points per contest | Contests entered | Total Borda points | First places | Mean AICc difference from the best | Mean Akaike weight | Mean rank |
|---|---|---|---|---|---|---|---|---|
| 1 | One-term exponential | 1.917 | 36 | 69 | 22 | 1.848 | 0.323 | 2.083 |
| 2 | Power law | 1.778 | 36 | 64 | 2 | 1.228 | 0.306 | 2.222 |
| 3 | Straight line | 1.417 | 36 | 51 | 3 | 4.109 | 0.164 | 2.583 |
| 4 | Logarithm | 0.889 | 36 | 32 | 9 | 3.896 | 0.207 | 3.111 |

**Table S15. Between-scenario differences in dependence.** For each of the 53 metric pairs not linked by definition and each pair of scenarios: the bias-corrected squared distance correlation $R^*_n$ in the first and in the second scenario, their difference, the permutation p-value and the Benjamini–Hochberg q-value over the 159 contrasts. Each metric was standardized within its scenario before the permutations (Materials and Methods). Rows are ordered by q.

| Pair | Contrast (first vs second) | $R^*_n$, first | $R^*_n$, second | Difference | p | q |
|---|---|---|---|---|---|---|
| MSD β-$t_\theta$ | No stimuli vs chemotaxis | 0.025 | 0.506 | -0.480 | 0.0001 | 0.016 |
| AS-$t_\theta$ | No stimuli vs chemotaxis | 0.001 | 0.440 | -0.439 | 0.0002 | 0.016 |
| AS-$t_\theta$ | Galvanotaxis vs chemotaxis | -0.015 | 0.440 | -0.455 | 0.0006 | 0.032 |
| MSD β-AS | No stimuli vs galvanotaxis | 0.405 | -0.015 | 0.421 | 0.0010 | 0.052 |
| IR-$t_\theta$ | No stimuli vs chemotaxis | -0.016 | 0.304 | -0.321 | 0.0020 | 0.060 |
| MSD β-AS | Galvanotaxis vs chemotaxis | -0.015 | 0.374 | -0.389 | 0.0030 | 0.080 |
| DR-$t_\theta$ | No stimuli vs galvanotaxis | -0.019 | 0.252 | -0.271 | 0.0050 | 0.087 |
| $t_\theta$-$AC_v(1)$ | No stimuli vs chemotaxis | 0.146 | -0.018 | 0.164 | 0.0050 | 0.087 |
| DR-AS | No stimuli vs galvanotaxis | 0.196 | -0.020 | 0.216 | 0.0050 | 0.087 |
| DFA γ-$t_\theta$ | No stimuli vs chemotaxis | 0.012 | 0.260 | -0.248 | 0.0060 | 0.087 |
| DFA γ-AS | No stimuli vs galvanotaxis | 0.316 | -0.023 | 0.339 | 0.0070 | 0.097 |
| DFA γ-AS | Galvanotaxis vs chemotaxis | -0.023 | 0.243 | -0.266 | 0.0100 | 0.129 |
| DR-AS | Galvanotaxis vs chemotaxis | -0.020 | 0.102 | -0.122 | 0.0120 | 0.150 |
| MSD β-DR | Galvanotaxis vs chemotaxis | 0.601 | 0.215 | 0.386 | 0.0140 | 0.159 |
| ApEn-$t_\theta$ | No stimuli vs chemotaxis | 0.237 | -0.015 | 0.252 | 0.0230 | 0.244 |
| MSD β-DR | No stimuli vs galvanotaxis | 0.302 | 0.601 | -0.299 | 0.0420 | 0.402 |
| IR-$t_\theta$ | No stimuli vs galvanotaxis | -0.016 | 0.117 | -0.134 | 0.0430 | 0.402 |
| MSD β-$t_\theta$ | No stimuli vs galvanotaxis | 0.025 | 0.238 | -0.213 | 0.0470 | 0.414 |
| ApEn-$t_\theta$ | No stimuli vs galvanotaxis | 0.237 | 0.014 | 0.224 | 0.0590 | 0.490 |
| MSD β-$t_\theta$ | Galvanotaxis vs chemotaxis | 0.238 | 0.506 | -0.267 | 0.0640 | 0.492 |
| RMSF α-AS | No stimuli vs chemotaxis | -0.035 | 0.016 | -0.051 | 0.0650 | 0.492 |
| RMSF α-IR | No stimuli vs galvanotaxis | -0.014 | 0.043 | -0.056 | 0.0760 | 0.551 |
| RMSF α-RMSF correlation time | No stimuli vs galvanotaxis | -0.018 | 0.027 | -0.046 | 0.0850 | 0.588 |
| RMSF α-$C_{\theta\text{-}v}$ | Galvanotaxis vs chemotaxis | 0.072 | -0.027 | 0.099 | 0.0890 | 0.588 |
| $t_\theta$-$AC_v(1)$ | No stimuli vs galvanotaxis | 0.146 | -0.013 | 0.159 | 0.0990 | 0.629 |
| RMSF α-AS | No stimuli vs galvanotaxis | -0.035 | 0.033 | -0.068 | 0.1040 | 0.636 |
| AS-$AC_v(1)$ | No stimuli vs chemotaxis | 0.224 | 0.016 | 0.208 | 0.1130 | 0.663 |
| DFA γ-DR | No stimuli vs galvanotaxis | 0.318 | 0.566 | -0.248 | 0.1290 | 0.732 |
| RMSF α-MSD β | No stimuli vs chemotaxis | -0.024 | 0.052 | -0.076 | 0.1370 | 0.753 |
| RMSF α-DFA γ | No stimuli vs chemotaxis | -0.012 | 0.011 | -0.023 | 0.1450 | 0.766 |
| DR-$t_\theta$ | Galvanotaxis vs chemotaxis | 0.252 | 0.062 | 0.191 | 0.1740 | 0.857 |
| RMSF correlation time-$t_\theta$ | No stimuli vs chemotaxis | 0.055 | -0.029 | 0.084 | 0.1770 | 0.857 |

| Pair | Contrast (first vs second) | $R^*_n$, first | $R^*_n$, second | Difference | p | q |
|---|---|---|---|---|---|---|
| DR-$t_\theta$ | No stimuli vs chemotaxis | -0.019 | 0.062 | -0.080 | 0.1940 | 0.857 |
| AS-$AC_v(1)$ | No stimuli vs galvanotaxis | 0.224 | 0.048 | 0.176 | 0.1980 | 0.857 |
| $C_{\theta\text{-}v}$-$t_\theta$ | No stimuli vs galvanotaxis | -0.016 | 0.029 | -0.045 | 0.2010 | 0.857 |
| RMSF α-$C_{\theta\text{-}v}$ | No stimuli vs galvanotaxis | -0.014 | 0.072 | -0.086 | 0.2130 | 0.857 |
| RMSF α-DR | Galvanotaxis vs chemotaxis | 0.007 | -0.033 | 0.041 | 0.2140 | 0.857 |
| DFA γ-$t_\theta$ | No stimuli vs galvanotaxis | 0.012 | 0.101 | -0.089 | 0.2170 | 0.857 |
| IR-$t_\theta$ | Galvanotaxis vs chemotaxis | 0.117 | 0.304 | -0.187 | 0.2180 | 0.857 |
| RMSF α-RMSF correlation time | Galvanotaxis vs chemotaxis | 0.027 | -0.020 | 0.048 | 0.2200 | 0.857 |
| RMSF α-DR | No stimuli vs galvanotaxis | -0.018 | 0.007 | -0.026 | 0.2240 | 0.857 |
| RMSF correlation time-AS | No stimuli vs galvanotaxis | -0.021 | 0.033 | -0.054 | 0.2310 | 0.857 |
| RMSF correlation time-ApEn | No stimuli vs galvanotaxis | 0.004 | -0.025 | 0.029 | 0.2370 | 0.857 |
| RMSF α-$AC_v(1)$ | Galvanotaxis vs chemotaxis | 0.186 | 0.359 | -0.174 | 0.2370 | 0.857 |
| RMSF correlation time-DFA γ | No stimuli vs chemotaxis | 0.029 | -0.036 | 0.065 | 0.2580 | 0.900 |
| DFA γ-$t_\theta$ | Galvanotaxis vs chemotaxis | 0.101 | 0.260 | -0.159 | 0.2690 | 0.900 |
| AS-$C_{\theta\text{-}v}$ | Galvanotaxis vs chemotaxis | -0.021 | 0.003 | -0.024 | 0.2740 | 0.900 |
| DR-$C_{\theta\text{-}v}$ | No stimuli vs galvanotaxis | -0.021 | 0.000 | -0.021 | 0.2900 | 0.900 |
| RMSF correlation time-ApEn | Galvanotaxis vs chemotaxis | -0.025 | 0.004 | -0.029 | 0.2900 | 0.900 |
| RMSF α-$t_\theta$ | No stimuli vs galvanotaxis | 0.034 | 0.013 | 0.021 | 0.2930 | 0.900 |
| RMSF correlation time-$t_\theta$ | Galvanotaxis vs chemotaxis | 0.027 | -0.029 | 0.056 | 0.2980 | 0.900 |
| RMSF α-ApEn | No stimuli vs chemotaxis | 0.041 | -0.020 | 0.061 | 0.3000 | 0.900 |
| MSD β-$C_{\theta\text{-}v}$ | No stimuli vs chemotaxis | -0.015 | 0.025 | -0.040 | 0.3040 | 0.900 |
| DFA γ-DR | Galvanotaxis vs chemotaxis | 0.566 | 0.393 | 0.173 | 0.3110 | 0.900 |
| RMSF correlation time-$AC_v(1)$ | No stimuli vs galvanotaxis | 0.012 | -0.025 | 0.036 | 0.3170 | 0.900 |
| RMSF α-$C_{\theta\text{-}v}$ | No stimuli vs chemotaxis | -0.014 | -0.027 | 0.013 | 0.3180 | 0.900 |
| RMSF α-IR | Galvanotaxis vs chemotaxis | 0.043 | -0.016 | 0.059 | 0.3410 | 0.900 |
| DFA γ-MSD β | No stimuli vs galvanotaxis | 0.537 | 0.670 | -0.132 | 0.3430 | 0.900 |
| $C_{\theta\text{-}v}$-$AC_v(1)$ | Galvanotaxis vs chemotaxis | 0.000 | -0.016 | 0.015 | 0.3460 | 0.900 |
| RMSF correlation time-DFA γ | Galvanotaxis vs chemotaxis | -0.009 | -0.036 | 0.027 | 0.3480 | 0.900 |
| RMSF correlation time-DR | No stimuli vs galvanotaxis | -0.005 | 0.015 | -0.019 | 0.3510 | 0.900 |
| IR-$AC_v(1)$ | Galvanotaxis vs chemotaxis | 0.094 | 0.012 | 0.082 | 0.3510 | 0.900 |
| RMSF α-DR | No stimuli vs chemotaxis | -0.018 | -0.033 | 0.015 | 0.3570 | 0.901 |
| RMSF correlation time-AS | No stimuli vs chemotaxis | -0.021 | 0.002 | -0.023 | 0.3850 | 0.955 |
| DR-$C_{\theta\text{-}v}$ | Galvanotaxis vs chemotaxis | 0.000 | -0.023 | 0.023 | 0.4070 | 0.969 |
| $C_{\theta\text{-}v}$-$t_\theta$ | No stimuli vs chemotaxis | -0.016 | 0.009 | -0.025 | 0.4130 | 0.969 |
| RMSF correlation time-$AC_v(1)$ | Galvanotaxis vs chemotaxis | -0.025 | -0.001 | -0.024 | 0.4320 | 0.969 |
| ApEn-$C_{\theta\text{-}v}$ | Galvanotaxis vs chemotaxis | -0.007 | -0.025 | 0.018 | 0.4330 | 0.969 |
| RMSF α-MSD β | Galvanotaxis vs chemotaxis | -0.006 | 0.052 | -0.058 | 0.4490 | 0.969 |
| DFA γ-$AC_v(1)$ | No stimuli vs galvanotaxis | 0.036 | 0.108 | -0.072 | 0.4520 | 0.969 |
| DFA γ-IR | Galvanotaxis vs chemotaxis | 0.357 | 0.457 | -0.100 | 0.4680 | 0.969 |
| RMSF correlation time-DR | Galvanotaxis vs chemotaxis | 0.015 | -0.006 | 0.020 | 0.4720 | 0.969 |
| DR-AS | No stimuli vs chemotaxis | 0.196 | 0.102 | 0.094 | 0.4790 | 0.969 |
| RMSF α-$t_\theta$ | No stimuli vs chemotaxis | 0.034 | 0.012 | 0.022 | 0.4790 | 0.969 |
| DR-$AC_v(1)$ | Galvanotaxis vs chemotaxis | 0.029 | -0.007 | 0.035 | 0.4880 | 0.969 |
| RMSF α-MSD β | No stimuli vs galvanotaxis | -0.024 | -0.006 | -0.018 | 0.4930 | 0.969 |
| AS-$C_{\theta\text{-}v}$ | No stimuli vs galvanotaxis | -0.004 | -0.021 | 0.017 | 0.4970 | 0.969 |
| RMSF α-DFA γ | No stimuli vs galvanotaxis | -0.012 | 0.007 | -0.019 | 0.5000 | 0.969 |
| MSD β-ApEn | No stimuli vs galvanotaxis | 0.023 | -0.015 | 0.038 | 0.5080 | 0.969 |
| ApEn-AS | No stimuli vs chemotaxis | -0.001 | -0.023 | 0.022 | 0.5090 | 0.969 |
| ApEn-$AC_v(1)$ | No stimuli vs galvanotaxis | 0.131 | 0.056 | 0.075 | 0.5140 | 0.969 |
| ApEn-AS | Galvanotaxis vs chemotaxis | 0.002 | -0.023 | 0.025 | 0.5150 | 0.969 |

| Pair | Contrast (first vs second) | $R^*_n$, first | $R^*_n$, second | Difference | p | q |
|---|---|---|---|---|---|---|
| MSD β-$C_{θ-v}$ | No stimuli vs galvanotaxis | -0.015 | 0.002 | -0.017 | 0.5190 | 0.969 |
| ApEn-$AC_v(1)$ | No stimuli vs chemotaxis | 0.131 | 0.056 | 0.075 | 0.5210 | 0.969 |
| MSD β-ApEn | Galvanotaxis vs chemotaxis | -0.015 | 0.009 | -0.024 | 0.5230 | 0.969 |
| AS-$t_θ$ | No stimuli vs galvanotaxis | 0.001 | -0.015 | 0.016 | 0.5270 | 0.969 |
| ApEn-DR | No stimuli vs chemotaxis | -0.012 | -0.029 | 0.017 | 0.5300 | 0.969 |
| RMSF correlation time-DFA γ | No stimuli vs galvanotaxis | 0.029 | -0.009 | 0.039 | 0.5420 | 0.969 |
| MSD β-DR | No stimuli vs chemotaxis | 0.302 | 0.215 | 0.087 | 0.5430 | 0.969 |
| RMSF α-ApEn | Galvanotaxis vs chemotaxis | 0.005 | -0.020 | 0.025 | 0.5600 | 0.972 |
| RMSF correlation time-AS | Galvanotaxis vs chemotaxis | 0.033 | 0.002 | 0.031 | 0.5650 | 0.972 |
| RMSF α-$AC_v(1)$ | No stimuli vs galvanotaxis | 0.273 | 0.186 | 0.088 | 0.5700 | 0.972 |
| MSD β-IR | Galvanotaxis vs chemotaxis | 0.339 | 0.426 | -0.087 | 0.5720 | 0.972 |
| RMSF α-$AC_v(1)$ | No stimuli vs chemotaxis | 0.273 | 0.359 | -0.086 | 0.5870 | 0.972 |
| RMSF α-ApEn | No stimuli vs galvanotaxis | 0.041 | 0.005 | 0.036 | 0.6010 | 0.972 |
| DFA γ-$C_{θ-v}$ | No stimuli vs galvanotaxis | -0.014 | -0.031 | 0.018 | 0.6030 | 0.972 |
| ApEn-$t_θ$ | Galvanotaxis vs chemotaxis | 0.014 | -0.015 | 0.028 | 0.6100 | 0.972 |
| $C_{θ-v}$-$AC_v(1)$ | No stimuli vs galvanotaxis | -0.021 | 0.000 | -0.021 | 0.6130 | 0.972 |
| ApEn-IR | Galvanotaxis vs chemotaxis | -0.006 | -0.023 | 0.016 | 0.6140 | 0.972 |
| DFA γ-ApEn | No stimuli vs chemotaxis | -0.006 | 0.008 | -0.014 | 0.6180 | 0.972 |
| DFA γ-$C_{θ-v}$ | Galvanotaxis vs chemotaxis | -0.031 | -0.014 | -0.017 | 0.6270 | 0.972 |
| DFA γ-MSD β | No stimuli vs chemotaxis | 0.537 | 0.605 | -0.068 | 0.6310 | 0.972 |
| DFA γ-DR | No stimuli vs chemotaxis | 0.318 | 0.393 | -0.076 | 0.6460 | 0.972 |
| DFA γ-MSD β | Galvanotaxis vs chemotaxis | 0.670 | 0.605 | 0.065 | 0.6560 | 0.972 |
| IR-$AC_v(1)$ | No stimuli vs galvanotaxis | 0.046 | 0.094 | -0.048 | 0.6770 | 0.972 |
| DFA γ-AS | No stimuli vs chemotaxis | 0.316 | 0.243 | 0.074 | 0.6780 | 0.972 |
| IR-$AC_v(1)$ | No stimuli vs chemotaxis | 0.046 | 0.012 | 0.035 | 0.6810 | 0.972 |
| ApEn-DR | Galvanotaxis vs chemotaxis | -0.018 | -0.029 | 0.012 | 0.6930 | 0.972 |
| MSD β-$AC_v(1)$ | Galvanotaxis vs chemotaxis | 0.046 | 0.086 | -0.040 | 0.6990 | 0.972 |
| IR-$C_{θ-v}$ | No stimuli vs galvanotaxis | -0.015 | -0.007 | -0.008 | 0.6990 | 0.972 |
| ApEn-$C_{θ-v}$ | No stimuli vs chemotaxis | -0.014 | -0.025 | 0.011 | 0.7100 | 0.972 |
| MSD β-IR | No stimuli vs galvanotaxis | 0.392 | 0.339 | 0.053 | 0.7130 | 0.972 |
| MSD β-$C_{θ-v}$ | Galvanotaxis vs chemotaxis | 0.002 | 0.025 | -0.023 | 0.7220 | 0.972 |
| DFA γ-$AC_v(1)$ | No stimuli vs chemotaxis | 0.036 | 0.075 | -0.039 | 0.7220 | 0.972 |
| AS-$AC_v(1)$ | Galvanotaxis vs chemotaxis | 0.048 | 0.016 | 0.032 | 0.7280 | 0.972 |
| DFA γ-IR | No stimuli vs galvanotaxis | 0.407 | 0.357 | 0.050 | 0.7320 | 0.972 |
| DFA γ-IR | No stimuli vs chemotaxis | 0.407 | 0.457 | -0.050 | 0.7340 | 0.972 |
| ApEn-IR | No stimuli vs galvanotaxis | -0.021 | -0.006 | -0.014 | 0.7340 | 0.972 |
| RMSF correlation time-$t_θ$ | No stimuli vs galvanotaxis | 0.055 | 0.027 | 0.028 | 0.7370 | 0.972 |
| DR-$AC_v(1)$ | No stimuli vs galvanotaxis | 0.005 | 0.029 | -0.024 | 0.7440 | 0.972 |
| RMSF correlation time-$AC_v(1)$ | No stimuli vs chemotaxis | 0.012 | -0.001 | 0.012 | 0.7470 | 0.972 |
| MSD β-$AC_v(1)$ | No stimuli vs galvanotaxis | 0.074 | 0.046 | 0.028 | 0.7480 | 0.972 |
| $C_{θ-v}$-$t_θ$ | Galvanotaxis vs chemotaxis | 0.029 | 0.009 | 0.020 | 0.7650 | 0.972 |
| DFA γ-$AC_v(1)$ | Galvanotaxis vs chemotaxis | 0.108 | 0.075 | 0.032 | 0.7730 | 0.972 |
| RMSF correlation time-IR | No stimuli vs chemotaxis | -0.010 | 0.003 | -0.013 | 0.7750 | 0.972 |
| MSD β-ApEn | No stimuli vs chemotaxis | 0.023 | 0.009 | 0.014 | 0.7980 | 0.972 |
| RMSF correlation time-IR | Galvanotaxis vs chemotaxis | -0.008 | 0.003 | -0.011 | 0.8040 | 0.972 |
| RMSF α-AS | Galvanotaxis vs chemotaxis | 0.033 | 0.016 | 0.017 | 0.8050 | 0.972 |
| MSD β-IR | No stimuli vs chemotaxis | 0.392 | 0.426 | -0.034 | 0.8080 | 0.972 |
| ApEn-$C_{θ-v}$ | No stimuli vs galvanotaxis | -0.014 | -0.007 | -0.007 | 0.8280 | 0.972 |
| DFA γ-ApEn | No stimuli vs galvanotaxis | -0.006 | 0.000 | -0.006 | 0.8310 | 0.972 |
| AS-$C_{θ-v}$ | No stimuli vs chemotaxis | -0.004 | 0.003 | -0.006 | 0.8350 | 0.972 |

| Pair | Contrast (first vs second) | $R^*_n$, first | $R^*_n$, second | Difference | p | q |
|---|---|---|---|---|---|---|
| IR-$C_{\theta\text{-}v}$ | Galvanotaxis vs chemotaxis | -0.007 | -0.012 | 0.006 | 0.8370 | 0.972 |
| MSD β-AS | No stimuli vs chemotaxis | 0.405 | 0.374 | 0.032 | 0.8450 | 0.972 |
| $t_\theta$-$AC_v(1)$ | Galvanotaxis vs chemotaxis | -0.013 | -0.018 | 0.005 | 0.8460 | 0.972 |
| RMSF correlation time-MSD β | No stimuli vs chemotaxis | -0.024 | -0.028 | 0.004 | 0.8470 | 0.972 |
| DR-$AC_v(1)$ | No stimuli vs chemotaxis | 0.005 | -0.007 | 0.011 | 0.8480 | 0.972 |
| RMSF correlation time-MSD β | Galvanotaxis vs chemotaxis | -0.024 | -0.028 | 0.004 | 0.8510 | 0.972 |
| DFA γ-ApEn | Galvanotaxis vs chemotaxis | 0.000 | 0.008 | -0.008 | 0.8520 | 0.972 |
| $C_{\theta\text{-}v}$-$AC_v(1)$ | No stimuli vs chemotaxis | -0.021 | -0.016 | -0.006 | 0.8580 | 0.972 |
| ApEn-DR | No stimuli vs galvanotaxis | -0.012 | -0.018 | 0.005 | 0.8680 | 0.972 |
| RMSF correlation time-$C_{\theta\text{-}v}$ | No stimuli vs galvanotaxis | -0.009 | -0.001 | -0.008 | 0.8680 | 0.972 |
| RMSF correlation time-$C_{\theta\text{-}v}$ | Galvanotaxis vs chemotaxis | -0.001 | -0.006 | 0.005 | 0.8790 | 0.972 |
| DR-$C_{\theta\text{-}v}$ | No stimuli vs chemotaxis | -0.021 | -0.023 | 0.002 | 0.8910 | 0.972 |
| IR-$C_{\theta\text{-}v}$ | No stimuli vs chemotaxis | -0.015 | -0.012 | -0.003 | 0.8920 | 0.972 |
| RMSF correlation time-$C_{\theta\text{-}v}$ | No stimuli vs chemotaxis | -0.009 | -0.006 | -0.003 | 0.8940 | 0.972 |
| RMSF α-IR | No stimuli vs chemotaxis | -0.014 | -0.016 | 0.003 | 0.8990 | 0.972 |
| RMSF α-RMSF correlation time | No stimuli vs chemotaxis | -0.018 | -0.020 | 0.002 | 0.9050 | 0.972 |
| MSD β-$AC_v(1)$ | No stimuli vs chemotaxis | 0.074 | 0.086 | -0.011 | 0.9290 | 0.991 |
| ApEn-IR | No stimuli vs chemotaxis | -0.021 | -0.023 | 0.002 | 0.9410 | 0.991 |
| ApEn-AS | No stimuli vs galvanotaxis | -0.001 | 0.002 | -0.003 | 0.9410 | 0.991 |
| RMSF α-DFA γ | Galvanotaxis vs chemotaxis | 0.007 | 0.011 | -0.004 | 0.9540 | 0.998 |
| RMSF correlation time-IR | No stimuli vs galvanotaxis | -0.010 | -0.008 | -0.002 | 0.9650 | 0.999 |
| RMSF correlation time-MSD β | No stimuli vs galvanotaxis | -0.024 | -0.024 | 0.000 | 0.9810 | 0.999 |
| DFA γ-$C_{\theta\text{-}v}$ | No stimuli vs chemotaxis | -0.014 | -0.014 | 0.001 | 0.9820 | 0.999 |
| RMSF α-$t_\theta$ | Galvanotaxis vs chemotaxis | 0.013 | 0.012 | 0.001 | 0.9890 | 0.999 |
| RMSF correlation time-DR | No stimuli vs chemotaxis | -0.005 | -0.006 | 0.001 | 0.9900 | 0.999 |
| RMSF correlation time-ApEn | No stimuli vs chemotaxis | 0.004 | 0.004 | 0.000 | 0.9960 | 0.999 |
| ApEn-$AC_v(1)$ | Galvanotaxis vs chemotaxis | 0.056 | 0.056 | 0.000 | 0.9990 | 0.999 |
